# Dynamics of *mcr-1* prevalence and *mcr-1*-positive *Escherichia coli* after the cessation of colistin use as a feed additive for animals in China: a prospective cross-sectional and whole genome sequencing based molecular epidemiological study

**DOI:** 10.1101/2020.02.03.923607

**Authors:** Cong Shen, Lan-Lan Zhong, Yongqiang Yang, Yohei Doi, David L Paterson, Nicole Stoesser, Furong Ma, Mohamed Abd El-Gawad El-Sayed Ahmed, Siyuan Feng, Songying Huang, Hong-Yu Li, Xi Huang, Xin Wen, Zihan Zhao, Minmin Lin, Guanping Chen, Wanfei Liang, Yingjian Liang, Yong Xia, Min Dai, Ding-Qiang Chen, Liyan Zhang, Kang Liao, Guo-Bao Tian

**Affiliations:** Department of Microbiology, Zhongshan School of Medicine, Sun Yat-sen University, Guangzhou 510080, China; Key Laboratory of Tropical Diseases Control (Sun Yat-sen University), Ministry of Education, Guangzhou 510080, China; School of Pharmaceutical Sciences (Shenzhen), Sun Yat-sen University, Guangzhou 510006, China; University of Pittsburgh School of Medicine, Pittsburgh 15261, Pennsylvania; Departments of Microbiology and Infectious Diseases, Fujita Health University School of Medicine, Aichi 470-1192, Japan; University of Queensland Centre for Clinical Research, The University of Queensland, Herston, Brisbane, QLD, Australia; Modernising Medical Microbiology, Nuffield Department of Medicine, University of Oxford, Oxford, United Kingdom; Department of Clinical Laboratory Medicine, Third Affiliated Hospital of Guangzhou Medical University, Guangzhou, China; Department of Microbiology and Immunology, Faculty of Pharmaceutical Sciences and Drug Manufacturing, Misr University for Science and Technology (MUST), Cairo, 6^th^ of October City, Egypt; Department of Clinical Laboratory, Sun Yat-sen Memorial Hospital, Sun Yat-sen University, Guangzhou, China; Program of Pathobiology and Immunology, the Fifth Affiliated Hospital of Sun Yat-sen University, Sun Yat-sen University, Zhuhai 519000, China; Department of Respiratory Medicine, the Fifth Affiliated Hospital of Sun Yat-sen University, Zhuhai 519000, China; School of Laboratory Medicine, Chengdu Medical College, Chengdu 610500, China; Division of Laboratory Medicine, Zhujiang Hospital, Southern Medical University, Guangzhou, Guangdong, 510282, China; Department of Clinical Laboratory, Guangdong Provincial People’s Hospital / Guangdong Academy of Medical Sciences, Guangzhou, Guangdong, 510080, China; Department of Clinical Laboratory, the First Affiliated Hospital of Sun Yat-Sen University, Guangzhou 510080, China

**Keywords:** Colistin, antimicrobial stewardship, intervention, *mcr-1*, *Escherichia coli*, genomic epidemiology

## Abstract

**Background:** The global dissemination of colistin resistance encoded by *mcr-1* has been attributed to extensive use of colistin in livestock, threatening colistin efficacy in medicine. The emergence of *mcr-1* in common pathogens, such as *Escherichia coli*, is of particular concern. Therefore, China banned the use of colistin in animal feed from May 1^ST^ 2017. We investigated subsequent changes in *mcr-1* prevalence, and the genomic epidemiology of *mcr-1*-positive *Escherichia coli* (MCRPEC).

**Methods:** Sampling was conducted pre- (October-December 2016) and post-colistin ban (October-December, 2017 and 2018, respectively). 3675 non-duplicate pig fecal samples were collected from 14 provinces (66 farms) in China to determine intervention-related changes in *mcr-1* prevalence. 15193 samples were collected from pigs, healthy human volunteers, colonized and infected hospital inpatients, food and the environment in Guangzhou, to characterize source-specific *mcr-1* prevalence and the wider ecological impact of the ban. From these samples, 688 MCRPEC were analyzed with whole genome sequencing (WGS), plasmid conjugation and S1-PFGE/Southern blots to characterize associated genomic changes.

**Findings:** After the ban, *mcr-1* prevalence decreased significantly in national pig farms, from 45·0% (308/684 samples) in 2016, to 19·4% (274/1416) in 2018 (p<0·0001). This trend was mirrored in samples from most sources in Guangzhou (overall 19·2% [959/5003 samples] in 2016; 5·3% [238/4489] in 2018; p<0·0001). The population structure of MCRPEC was diverse (23 sequence clusters [SCs]); ST10 clonal complex isolates were predominant (247/688 [36%]). MCRPEC causing infection in hospitalized inpatients were genetically more distinct and appeared less affected by the ban. *mcr-1* was predominantly found on plasmids (632/688 [92%]). Common *mcr-1* plasmid types included IncX4, IncI2 and IncHI2 (502/656 [76.5%]); significant increases in IncI2-associated *mcr-1* and a distinct lineage of *mcr-1*-associated IncHI2 were observed post-ban. Changes in the frequency of *mcr-1*-associated flanking sequences (IS*Apl1*-negative MCRPEC), 63 core genome SNPs and 30 accessory genes were also significantly different after the ban, consistent with rapid genetic adaptation in response to changing selection pressures.

**Interpretation:** A rapid, ecosystem-wide, decline in *mcr-1* was observed after banning the use of colistin in animal feed, with associated genetic changes in MCRPEC. Genomic surveillance is key to assessing and monitoring stewardship interventions.

**Funding:** National Natural Science Foundation of China

## Introduction

The emergence of antimicrobial resistant pathogens, particularly carbapenem-resistant Enterobacteriaceae (CRE), *Acinetobacter baumannii* (CRAB) and *Pseudomonas aeruginosa* (CRPA), is a global public health crisis.^1^ Colistin is one of the antibiotics of last resort for managing multidrug-resistant Gram-negative infections, especially carbapenem-resistant Gram-negative infections.^2^ However, its use has been threatened by the emergence of plasmid-mediated colistin resistance encoded by *mcr-1*, which was first identified in China in 2016.^3^ In just four years, *mcr-1* has been reported in more than 70 countries across five continents, in numerous species from multiple ecological niches, and in association with diverse plasmids such as IncX4, IncI2 and IncHI2.^4,5^ Moreover, *mcr-1* is associated with other resistance genes, including ESBLs and carbapenemases, and its dissemination amongst common human pathogens, such as *Escherichia coli*, could considerably compromise clinical treatments.^6–8^ Therefore, efficient interventions to limit the spread of *mcr-1* are urgently needed.

In the past two decades, with the global expansion of industrial livestock production to fulfill increasing demand for meat products, the proportion of antimicrobial resistance (AMR) in food animals has been increasing dramatically.^9^ China is the world’s largest poultry and pig producer, one of the world’s highest users of colistin in animal husbandry, and has the highest prevalence of *mcr-1*-positive Enterobacterales globally.^9^ The widespread dissemination of *mcr-1* could be attributable to selective pressures exerted by the heavy use of colistin in animal husbandry in China, which may encourage the spread of *mcr-1*-positive isolates and enhance horizontal transfer of *mcr-1*-harboring plasmids.^3,10^ Evidence to date strongly suggests that *mcr-1* has spread to humans and other ecological niches from farm animal reservoirs.^3,5,11^ As a result, China banned the use of colistin use as a feed additive for animals in May 1^st^ 2017, which led to a decline in usage of more than 8000 tons per year.^12^ The impact of this intervention on *mcr-1* prevalence and associated genetic changes in *mcr-1*-positive isolates remain to be evaluated.

The primary aim of this study was to investigate changes in *mcr-1* prevalence in pigs and other source populations in response to the ban, using whole genome sequencing to track the associated genomic changes in *mcr-1*-positive *Escherichia coli* (MCRPEC) populations. We first investigated changes in *mcr-1* prevalence in farmed pigs across China, and source-specific changes in *mcr-1* prevalence in pigs and other sources (humans, food, the environment) in a single center, namely Guangzhou, the capital city of Guangdong province with catchment of 15 million people. To describe the genetic dynamics of MCRPEC populations in response to the ban and in the context of various ecological niches, we carried out a large-scale, detailed, genome-based analysis.

## Methods

### Study setting and design

We undertook a prospective cross-sectional and molecular epidemiological study across two time periods, occurring pre-and post-the implementation of a national ban on the use of colistin in animal feed in China from May 1^st^ 2017. The pre-intervention period spanned from October 1^st^ to December 31^st^ 2016, with follow-up, post-intervention annual surveillance from October 1^st^ to December 31^st^ 2017 and October 1^st^ to December 31^st^ 2018, respectively.

To determine national changes in *mcr-1* prevalence in farmed pigs pre-and post-ban, a total of 3675 non-duplicate fecal samples were collected from pigs on 66 farms in 14 provinces in China (figure 1 and table 1). These provinces accounted for 65.6%-70% of total pork slaughtered and 65.0%-66.6% of total pork consumption during 2016-2018 in China (http://www.stats.gov.cn/tjsj/ndsj/).

**Figure 1.**
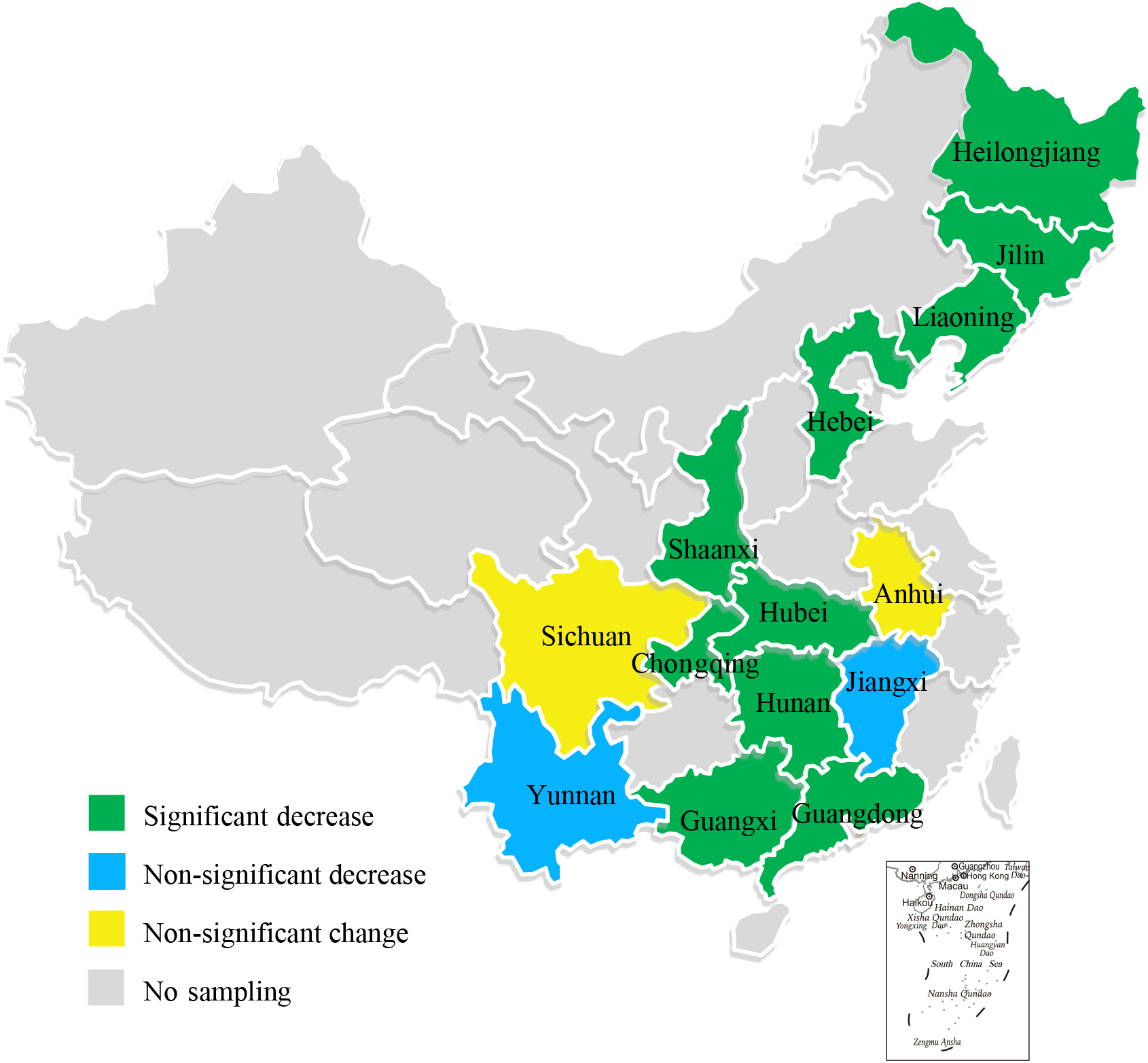
Map of 14 provinces sampled for the national pig farming study, and mcr-1 prevalence trends pre-and post-colistin ban.

**Table 1.**
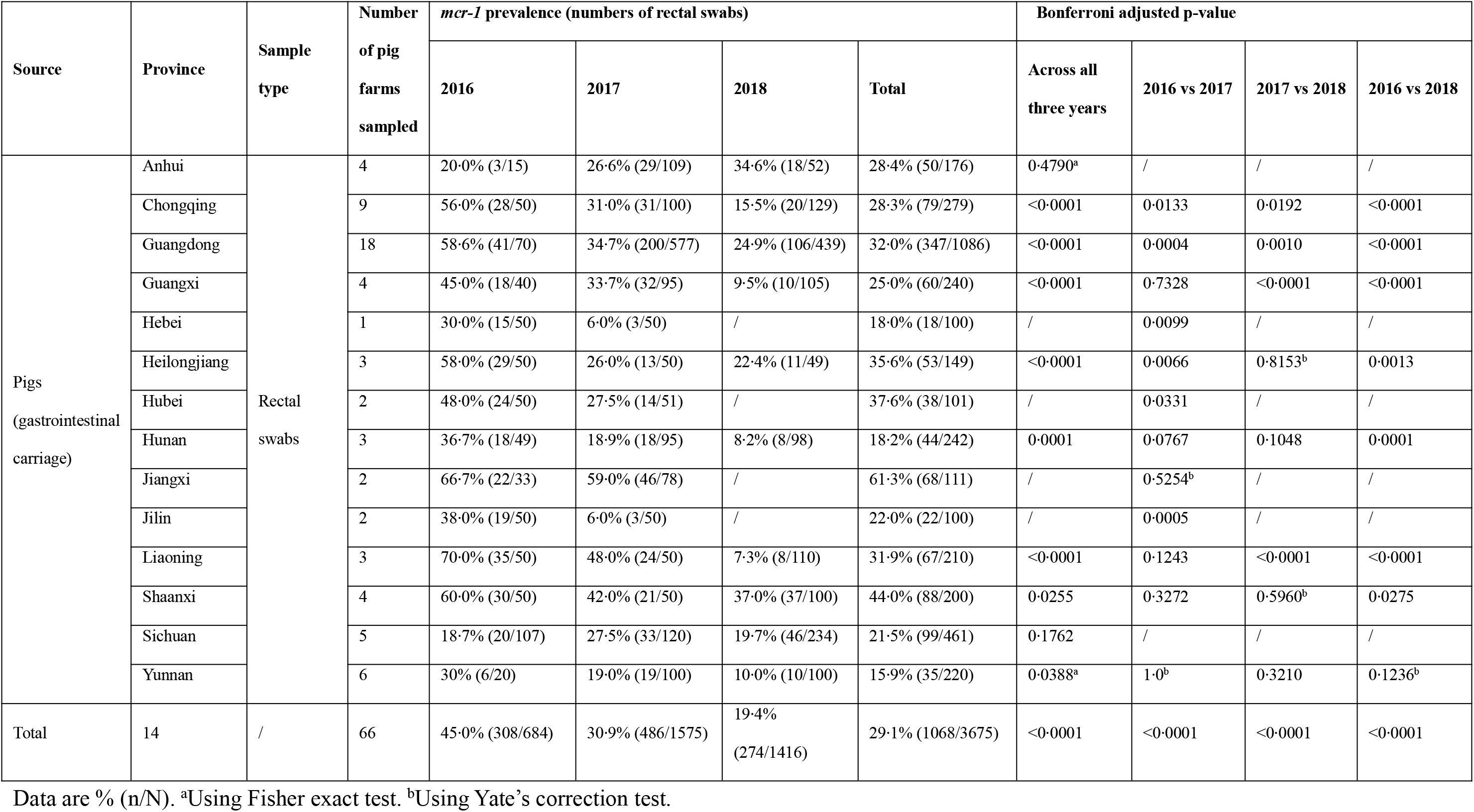
Characteristics of sample collection and *mcr-1* prevalence for national prospective epidemiological study.

To investigate changes in *mcr-1* prevalence and the molecular epidemiology of *mcr-1* in more detail across a regional ecosystem (including human and porcine gastrointestinal carriage, human infection, foodstuffs, and environmental samples) in response to the colistin ban, we performed sampling in Guangzhou, the capital city of Guangdong province, with a catchment of 15 million people over ~10000 km^2^. In total, 15193 non-duplicate samples were collected in Guangzhou from 2016-2018, including 2214 rectal swabs from pigs across 23 farms and two slaughterhouses, 3422 rectal swabs from healthy volunteers, 2395 rectal swabs from inpatients in two tertiary general hospitals containing over 6000 beds, 3724 clinical isolates from inpatients with infections caused by Enterobacterales in eight hospitals, 2632 environmental samples from residential areas and slaughterhouses, and 806 food samples from markets and slaughterhouses (figure 2 and table 2). All samples were collected over the same timeframe as for the national pig farm study, except clinical samples from patients with Enterobacterales infections which were collected monthly from January 1^st^, 2016 to December 31^st^, 2018.

**Figure 2.**
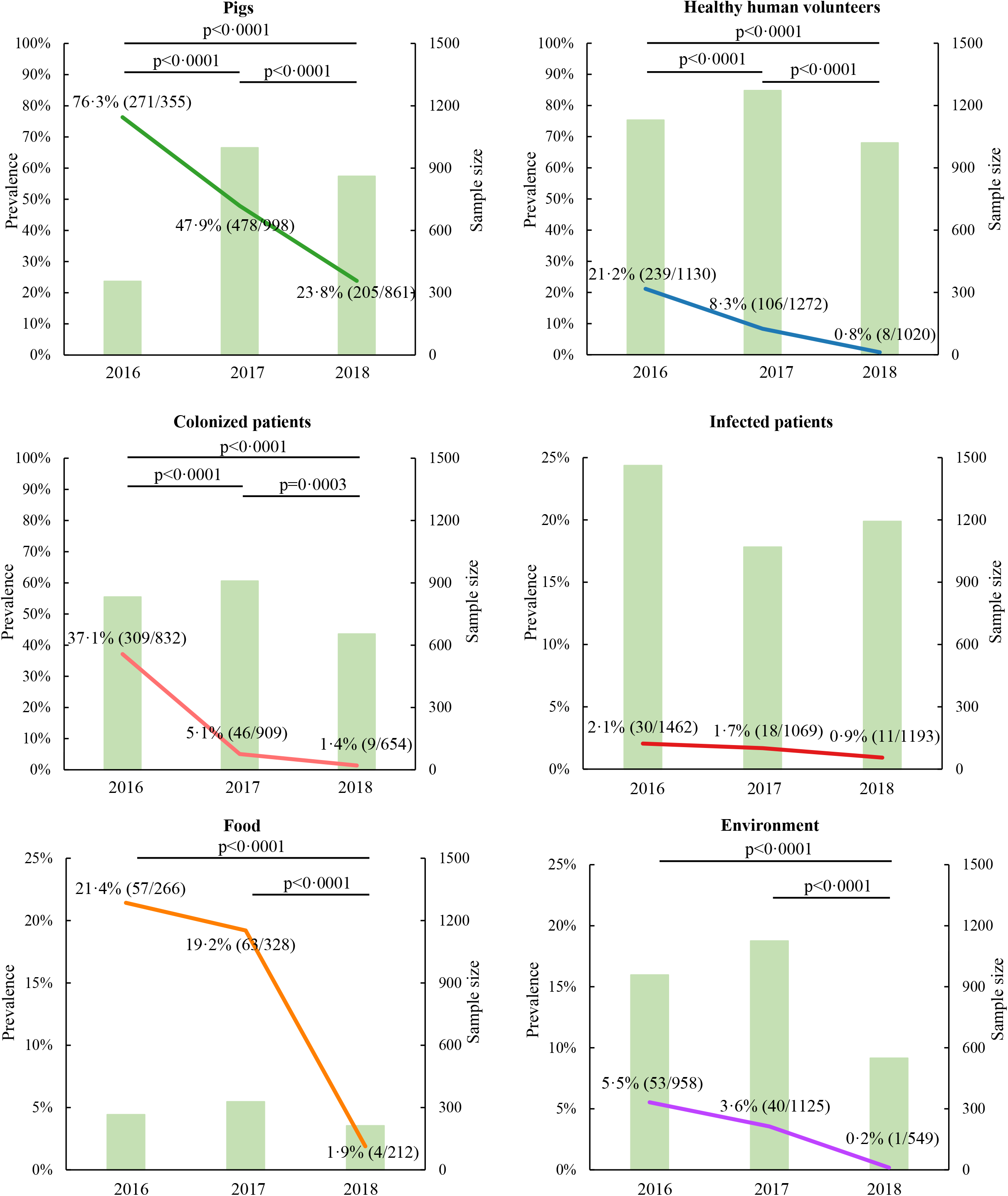
*mcr-1* prevalence in ecological samples in Guangzhou 2016-2018.

**Table 2.**
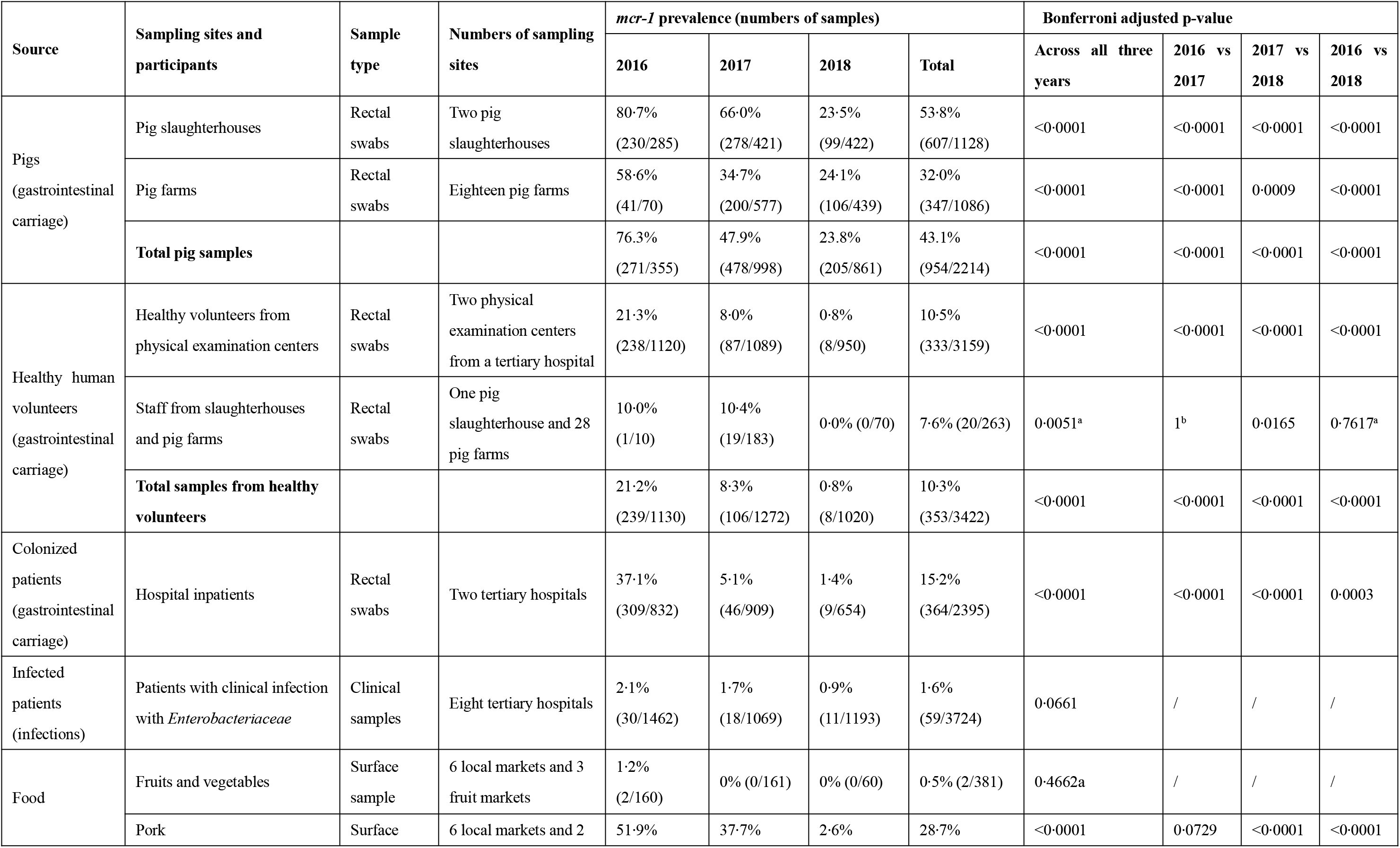
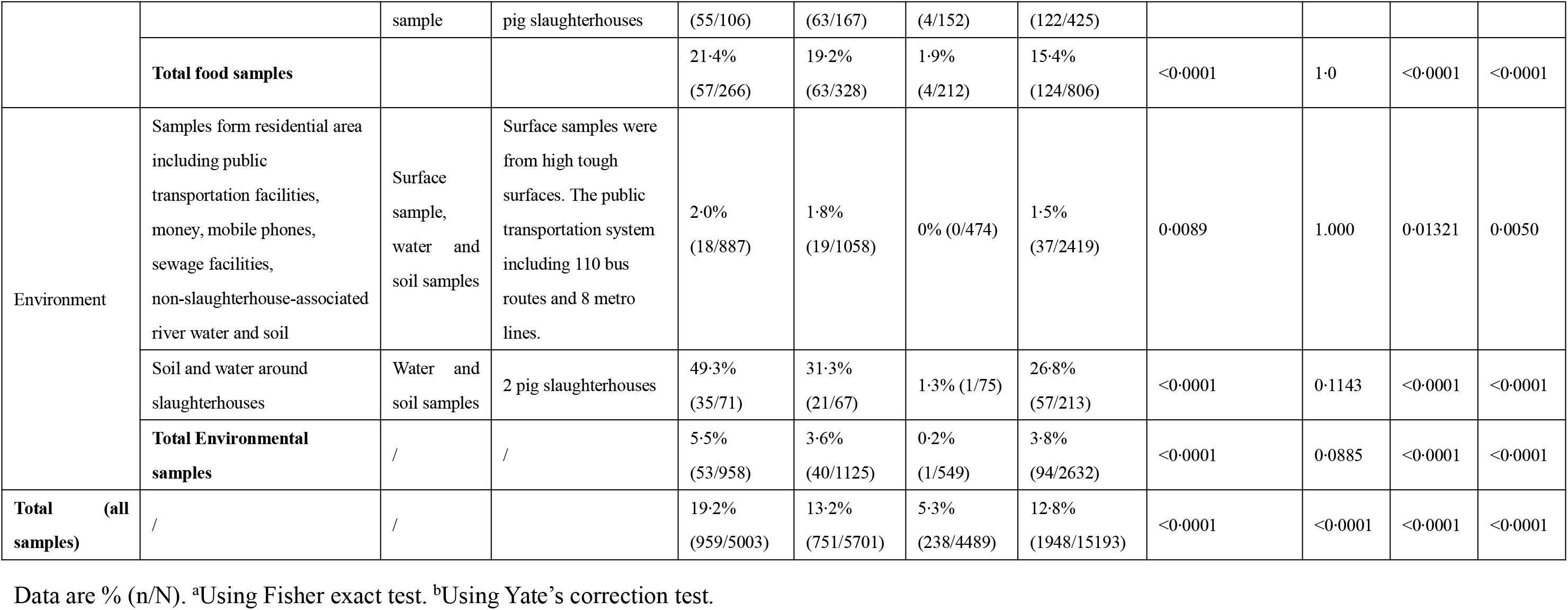
Characteristics of sample collection and prevalence for regional prospective epidemiological study.

To determine the genetic dynamics of *mcr-1*-positive bacteria after the intervention, we focused on *mcr-1*-positive *E. coli* (MCRPEC) cultured from *mcr-1*-positive samples in Guangzhou, as *E. coli* was the predominant bacterial host of *mcr-1* identified across all sample types in this study. We performed whole genome sequencing (WGS) of MCRPEC to evaluate several genetic characteristics, including strain relatedness (core genome single-nucleotide-polymorphism [cgSNPs]), and accessory genes, plasmids, and transposons present. In total, 755/1948 available MCRPEC were selected for sequencing based on the sample size of each epidemiological group. We included all MCRPEC for WGS analysis when the number of MCRPEC isolated across samples for an epidemiological group was ≤100 isolates per year; if the number of isolates cultured for an epidemiological group was >100/year, we randomly selected ~25% (~70) MCRPEC for WGS analysis (appendix table 1).

### Inclusion and exclusion criteria

For inclusion in the group defined as patients with infections caused by Enterobacterales, individuals had to have been admitted for ≥2 days on any of the medical, surgical, gynaecologic/obstetric, paediatric, psychiatric units of the two study hospitals. Clinical samples and cultured isolates were collected as part of clinical management and hospital surveillance. The source of infection was determined as pneumonia, urinary tract infection, surgical site infection, intra-abdominal infection, catheter-related infection, or bacteremia as defined by the Centers for Disease Control and Prevention.^13^ Patients with all forms of gastroenteritis and those with active bleeding per rectum and/or anal fissures were excluded.

For inclusion in the human inpatient gastrointestinal carriage group, inpatients hospitalized for ≥2 days had to be sampled within 24hrs of hospitalization; for inclusion in the healthy gastrointestinal carriage group, individuals had to be sampled within 12 hours of a normal physical exam. Exclusion criteria for enrolment in the human gastrointestinal carriage groups (both inpatients and healthy volunteers) included age ≤28 days, pregnancy, and clinical presentations of gastroenteritis, gastrointestinal cancer (esophageal cancer, gastric cancer, colon cancer, etc.), peptic ulcer, gastrointestinal bleeding, inflammatory bowel disease (Crohn’s disease, ulcerative colitis, etc.), intestinal polyps, intestinal fistula, anal fistula or anal fissure or requiring gastrointestinal surgery.

### Ethics and consent

Ethical approval was sought and given by Sun Yat-Sen University ZhongShan School of Medicine (November 1^st^, 2014). Individual consent forms were translated into Mandarin Chinese and consent was obtained for all patients and healthy volunteers. Patients were individually contacted, either face-to-face or by phone and consent sought to use these samples/data for this study described herein. All participants had the right to withdraw from the study at any stage.

### Procedures

### Sample collection, sample culture, *mcr-1* screening and isolate sub-culture and identification

For the national pig farm study, rectal swabs were collected from individual pigs; up to three non-duplicate samples were collected from pigs each pigsty. Feed samples were collected simultaneously for measurement of colistin concentrations to corroborate the effective implementation of the ban over the study time period.

For the study in Guangzhou, rectal swabs from live pigs were collected as for the national study, or using rectal swabs to obtain samples of endorectal faeces from individual pigs in slaughterhouses. Human gastrointestinal carriage samples (both inpatient and healthy volunteers) were collected using rectal swabs. Samples from patients with infections with *Enterobacteriaceae* were collected as part of routine clinical management and/or hospital surveillance. For food samples, surface samples were obtained using cotton swabs moistened with sterile saline. For environmental samples, surface samples were obtained as for food; in addition, water and soil samples were collected in separate sterile containers. All samples were obtained using sterile swabs/containers, refrigerated after collection, and transported for laboratory processing within two hours of collection.

All samples were cultured on non-selective nutrient broth media overnight at 37°C; DNA was extracted from sweeps of cultured colonies, which were also stored. Clinical samples (urine, blood, sputum, wound samples) from patients with infections were plated on Columbia blood agar (CBA) with 5% sheep blood (Luqiao, Beijing, China); if *Enterobacteriaceae* were identified, sweeps of cultured growth from the original CBA plates were both stored and underwent DNA extraction. Total DNA was extracted using the boiling method and screened for the presence of *mcr-1* by polymerase chain reaction (PCR) as previously described.^14^ *mcr-1*-positive samples were re-cultured on selective agar (MacConkey + colistin [2mg/L]) and incubated for 18-24 hours, 37°C). Subsequently, up to 10 *Enterobacteriaceae* colonies were sub-cultured to MacConkey agar with colistin (2 mg/L); species identification was first confirmed by MALDI-TOF MS (BrukerDaltonik GmbH, Bremen, Germany). Where the species could not be reliably assigned by MALDI-TOF, 16S rDNA sequencing was applied (appendix methods). Only one suspected *mcr-1*-positive isolate, chosen at random, was analyzed per sample.

### Phenotypic characterization

Minimum inhibitory concentrations (MICs) for 16 antimicrobial agents were determined for all *mcr-1*-positive isolates by agar dilution, except for colistin and polymyxin B MICs, which were determined using the broth dilution method (EUCAST breakpoints, version 9.0; Clinical and Laboratory Standards Institute, document M100-S29). Conjugation experiments were performed using streptomycin-resistant *E. coli* C600 as the recipient (appendix methods).

### WGS procedures and analyses

DNA was extracted from MCRPEC isolates and sequenced using the Illumina HiSeq 2000 platform. Trimmed and quality-controlled reads were assembled using SPAdes.^15^ *In silico* MLST, plasmid replicon, virulence gene, insertion sequence and resistance gene identification were determined using BLASTn based comparisons with reference databases, and/or SRST2 and ABRicate (https://github.com/tseemann/abricate).^16,17^ Draft genomes were annotated using Prokka.^18^ Clusters of orthologous genes (COGs) were annotated using eggNOG.^19^ Core genome extraction and pan-genome analysis was performed using Roary,^20^ and core genome single nucleotide polymorphisms (cgSNPs) extracted from this concatenated alignment using SNP-sites.^21^ Due to the limitations of short-read assemblies in directly assigning the location of *mcr-1* to particular plasmids, we developed a new method to approximate the association of *mcr-1* with plasmid background and replicon type (appendix methods and appendix figure 9). Presence/absence of Tn*6330* (IS*Apl1-mcr-1-pap2-ISApl1*) in association with *mcr-1* was determined by BLASTn comparisons using *mcr-1*-positive contigs as the query sequence, and confirmed with PCR/Sanger sequencing in cases of uncertainty. Genome-and pan-genome wide association analyses, to identify: (a) cgSNPs associated with pre-/post-intervention time periods; (b) accessory genes associated with pre-/post-intervention time periods; (c) accessory genes associated with epidemiological groups, were done using Scoary.^22^ Details of WGS procedures and analyses, including software versions and parameters used, can be found in the appendix methods. Sequencing data have been deposited with the NCBI (BioProject accession: PRJNA593695).

### Statistical analyses

Statistical analyses and random selection were performed using R. Differences in *mcr-*prevalence and antimicrobial resistance rates were assessed using the Chi-square test, Fisher’s exact test or Yates’ correction with Bonferroni-adjusted p-values. Antibiotic MICs were log-transformed (base-2) and differences in distributions assessed using Kruskal-Wallis or Wilcoxon-Mann-Whitney tests with Bonferroni correction of p-values for multiple comparisons. p-values <0.05 were considered significant, unless otherwise specified.

## Results

### *mcr-1* prevalence in national pig farms

To observe the nationwide changes in *mcr-1* prevalence among pigs in response to the ban on colistin use as a feed additive in animals, we analyzed rectal swabs from 3675 pigs from 66 farms in 14 provinces in China (figure 1 and table 1). Overall, 1068/3675 (29·1%) rectal swabs were *mcr-1*-positive; *mcr-1* prevalence significantly decreased from 45·0% in 2016 (308/684 samples) to 30·9% in 2017 (486/1575 samples, Bonferroni-adjusted p<0.0001), and further to 19·4% in 2018 (274/1416, Bonferroni-adjusted p<0.0001) (table 1). *mcr-1* prevalence was decreased in most of the surveyed provinces after the intervention (12/14 provinces [86%]), including significant decreases observed in ten provinces (p<0.0001) and downward trends in two others (figure 1 and table 1).

Pig farms sampled in provinces colored green demonstrated significant declines in mcr-1 prevalence pre-and post-colistin ban; those colored blue demonstrated non-significant declines; those colored yellow demonstrated no obvious trends. No samples were obtained from sites in provinces colored grey.

### *mcr-1* prevalence amongst pigs, humans, food and environmental samples in Guangzhou

To more fully investigate the wider impact of the ban on changes in *mcr-1* prevalence across an ecosystem, we collected a total of 15193 samples from different sources in Guangzhou (figure 2 and table 2). Of these, 1948 (12o8%) were *mcr-1*-positive (table 2); with decreases in *mcr-1* prevalence pre-and post-ban mirroring those seen in pig farms nationally, namely from 19·2% in 2016 (959/5003 samples) to 13·2% in 2017 (751/5701, Bonferroni-adjusted p<0·0001) and 5·3% in 2018 (238/4489, Bonferroni-adjusted p<0·0001) (table 2). Specifically, as expected, the *mcr-1* prevalence in Guangzhou pigs decreased significantly from 76·3% in 2016 (271/355 samples) to 47·9% in 2017 (478/998, Bonferroni-adjusted p<0·0001) and 23·8% in 2018 (205/861, Bonferroni-adjusted p<0·0001). In addition, *mcr-1* prevalence was very high amongst rectal swab samples from healthy volunteers (21·3%, 238/1120), colonized inpatients (37·1%, 309/832), pork for consumption (51o9%, 55/106) and river water and soil samples around slaughterhouses (49·3%, 35/71) in 2016, whereas it was rarely identified in these sample types in 2018, with significant linear decreases observed across all these groups (table 2). In contrast, *mcr-1* prevalence amongst Enterobacteriaceae isolates causing infection was low at baseline (2·1%, 30/1462 isolates in 2016), with only slight reductions observed following the ban (figure 2 and table 2). As to the food and environmental samples, the *mcr-1* prevalence decreased significantly in slaughterhouse-associated samples such as pork, river water and soil around slaughterhouse, whereas a low and continuous decreasing prevalence of *mcr-1* was observed in other food samples such as fruits and vegetables and other environmental samples such as surface swabs from public transportation facilities, money, mobile phones, sewage facilities, and non-slaughterhouse-associated river water and soil (table 2).

Left y-axis and line plot represent mcr-1 prevalence for each year. Right y-axis and histogram show sampling size for each year. Data are % (n/N). All p-values are Bonferroni-adjusted.

### Species distribution of *mcr-1*-positive isolates obtained from samples

All 1948 *mcr-1*-positive samples from Guangzhou were sub-cultured to identify *mcr-1*-positive isolates. 1914 (98·3%) *mcr-1*-positive isolates were *Escherichia coli*, followed by 28 (1·4%) *Klebsiella pneumoniae*, two each of *Citrobacter freundii, Enterobacter cloacae*, and one each of *Enterobacter asburiae* and *Enterobacter aerogenes*.

### Antibiotic susceptibility profile of *mcr-1*-positive *E. coli* (MCRPEC)

Antibiotic susceptibility testing against 16 antibiotics was performed for the 1948 MCRPEC cultured from Guangzhou (figure 3, appendix figure 1, and appendix table 2-3). Overall, 1944 (99·8%) isolates were non-susceptible to colistin or polymyxin B, followed by ampicillin, amoxicillin-clavulanic acid and ciprofloxacin. The antibiotics to which isolates were most frequently susceptible were imipenem, ertapenem, meropenem, and amikacin. The resistance rates and MICs of gentamicin, ampicillin, amoxicillin-clavulanic acid, cefotaxime, nitrofurantoin, and ceftazidime increased significantly from 2016 to 2018, while decreased resistance from 2016 to 2018 was observed for fosfomycin and tigecycline (figure 3 and appendix table 2).

**Figure 3.**
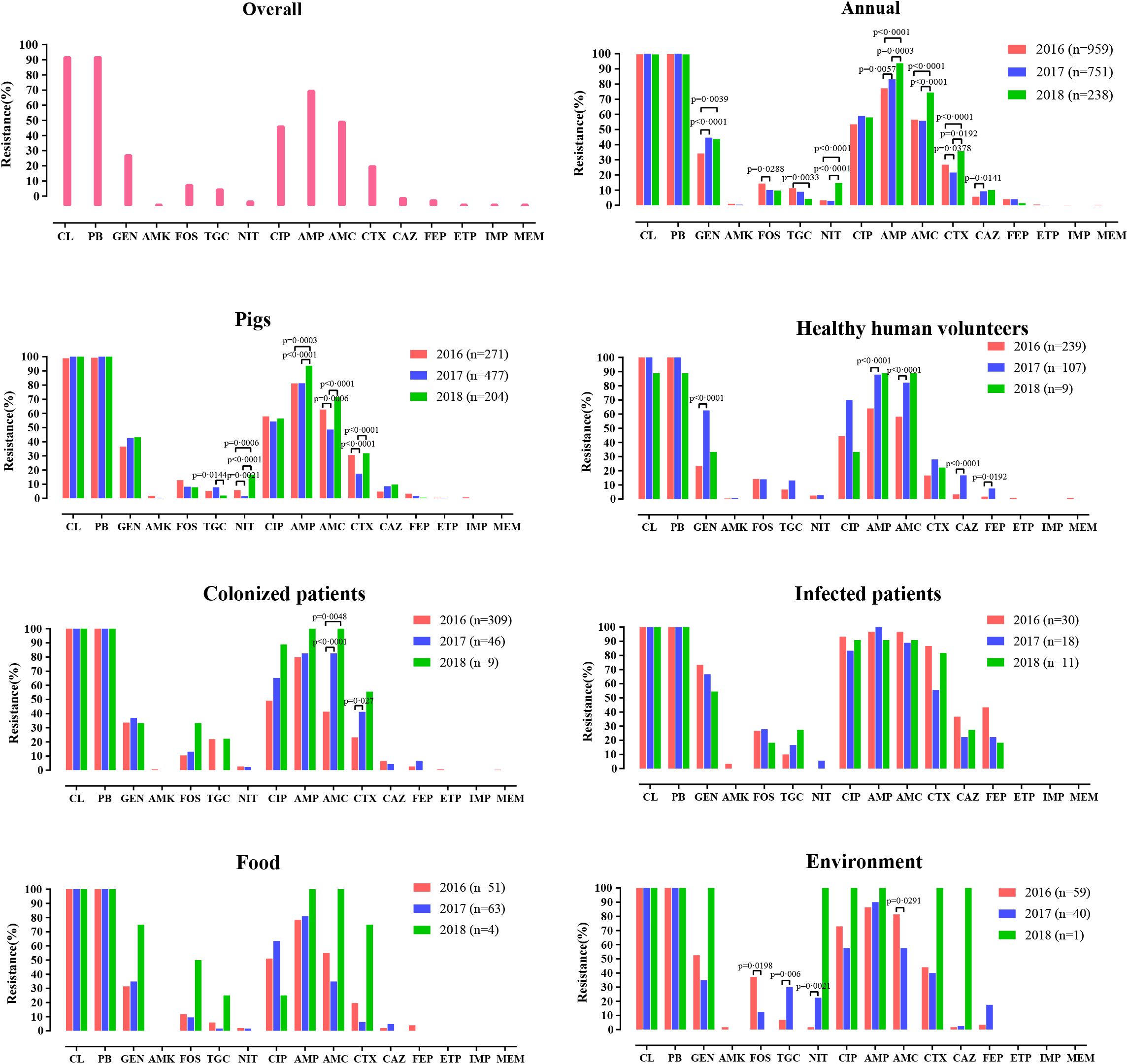
Antimicrobial susceptibility profiles of 1948 *mcr-1* positive *E. coli* (MCRPEC) cultured in Guangzhou by sampling year and source.

MIC values were analysed to determine the resistance level of antimicrobials (appendix figure 1 and appendix table 3-4). Generally, the MIC values of most antimicrobials were higher in the post-intervention population (appendix figure 1). Unexpectedly, compared with 2016, MIC values of colistin and polymyxin B were higher in 2017; but lower in 2018 (appendix figure 1). Remarkably, pairwise comparisons of MIC values of *mcr-1-*positive isolates from different sources revealed that *mcr-1-*positive isolates collected from patients with infections were more multi-drug resistant and more persistently drug-resistant across all three study years, (appendix table 4).

Bonferroni-adjusted p-values <0·05 are shown. CL = colistin, PB = polymyxin B, TGC=tigecycline, AMP=ampicillin, AMC= amoxicillin-clavulanate, CTX=cefotaxime, CAZ=ceftazidime, FEP=cefepime, GEN=gentamicin, AMK=amikacin, ETP=ertapenem, IMP=imipenem, MEM=meropenem, FOS=fosfomycin, NIT=nitrofurantoin, CIP=ciprofloxacin.

### Genomic population structure and distribution of characteristics of *mcr-1*-positive *E. coli* (MCRPEC)

We performed WGS for 755/1948 (38·8%) cultured MCRPEC from Guangzhou, with 688/755 (91·1%) genomes available following quality control. Of these, 330 were collected in 2016, 269 in 2017 and 89 in 2018, including 199 from pigs, 144 from healthy people, 110 from colonization inpatients, 55 from patients with Enterobacteriaceae infections, 107 from food samples, and 73 from environmental samples (appendix table 1).

The 688 sequenced MCRPEC were assigned to 198 distinct STs. The most prevalent STs were ST10 (n=100), ST101 (n=46), ST48 (n=28), ST69 (n=26), ST744 (n=22), ST1244 (n=18), ST410 (n=15), ST165 (n=14), ST206 (n=14), ST117 (n=11), ST641 (n=11) and ST58 (n=10), which accounted for 45·8% of MCRPEC. A total of 247 MCRPEC belonged to the ST10 clonal complex (CC), which represented the predominant group across the three years. Notably, ST10 CC were only detected in samples from animals (n=15) and the environment (n=1) in 2018. Forty-two new STs were identified in 53 isolates; whereas 112 STs were uniquely represented by one isolate (appendix table 5).

### Characteristics of resistance genes and virulence factors of *mcr-1*-positive *E. coli* (MCRPEC)

Antimicrobial resistance genes (ARGs) and virulence factors (VFs) were identified for 688 sequenced MCRPEC (appendix figure 2-4). ARG analysis showed that all MCRPEC harbored *mcr-1* or its variants *(mcr-1.2* (n=5), *mcr-1.6* (7), *mcr-1.12* (1), *mcr-1.13* (1), and *mcr-1.14* (1)), and 13 coexisted with *mcr-3* or its variants *(mcr-3.5* (7) and *mcr-3.21* (1)); no other *mcr* variant was found (appendix figure 2 and 4). There was no significant difference in the average number of ARGs across the three years (13.7±4.6 in 2016, 13.1±4.4 in 2017, 14.1±4.3 in 2018, p=0.0724, Chi-squared test). However, the average number of VFs in 2017 (159.6±24.9) and 2018 (171.4±25.4) was significantly higher than in 2016 (172.3±26.2) (2016 *vs* 2017, t=-5.708244, p<0·0001; 2016 *vs* 2018, t=-4.207219, p<0·0001; 2017 vs 2018, t=-0.278680, p=0·78). Notably, the average number of both ARGs and VFs was significantly higher in isolates associated with infection than from other sources (appendix figure 2 and 3).

### Core genome-based and accessory genome-based population structure of *mcr-1*-positive *E. coli* (MCRPEC)

We analyzed the population structure using both core genes and accessory genes of 688 MCRPEC. As a result, a high correlation was observed between the nucleotide divergence of core genes and content differences of accessory genes in MCRPEC (p<0·0001, r=0.8108, Mantel test) (appendix figure 5), indicating that core genome and accessory genome correlate for each given MCRPEC.

Pan-genome analysis of 688 MCRPEC identified 2290 core genes representing a 1.2 Mb alignment in ≥99% of genomes. hierBAPS analysis of the MCRPEC population structure based on 43648 cgSNPs identified 23 sequence clusters (SCs), of which two (SC8 and SC23) were polyphyletic in the ML phylogeny (figure 4). SC8 was bifurcated as two distinct monophyletic clades; whereas SC23 was comprised of low-frequency genotypes. The MCRPEC had diverse population structures and did not form specific host/source-related clusters (figure 4 and 5A); the majority of the clusters were detected in isolates from all sources. MCRPEC from pigs were however typically found in discrete and dominant clades interspersed among isolates from other sources. The prevalence and widespread distribution of MCRPEC from pigs among most of the clusters represent a major reservoir for the spread of MCRPEC to other hosts. SC8, SC11, SC15-17, and SC19-21 were the leading lineages harboring MCRPEC from all sources (figure 4 and 5A). Compared with 2016 and 2017, the number of MCRPEC in each cluster reduced in 2018, leading to a decrease of diversity within each cluster (figure 5A). Notably, most MCRPEC causing infection were concentrated in SC1 to SC4, and were genetically distant from MRCPEC from other sources (figure 4 and 5A).

**Figure 4.**
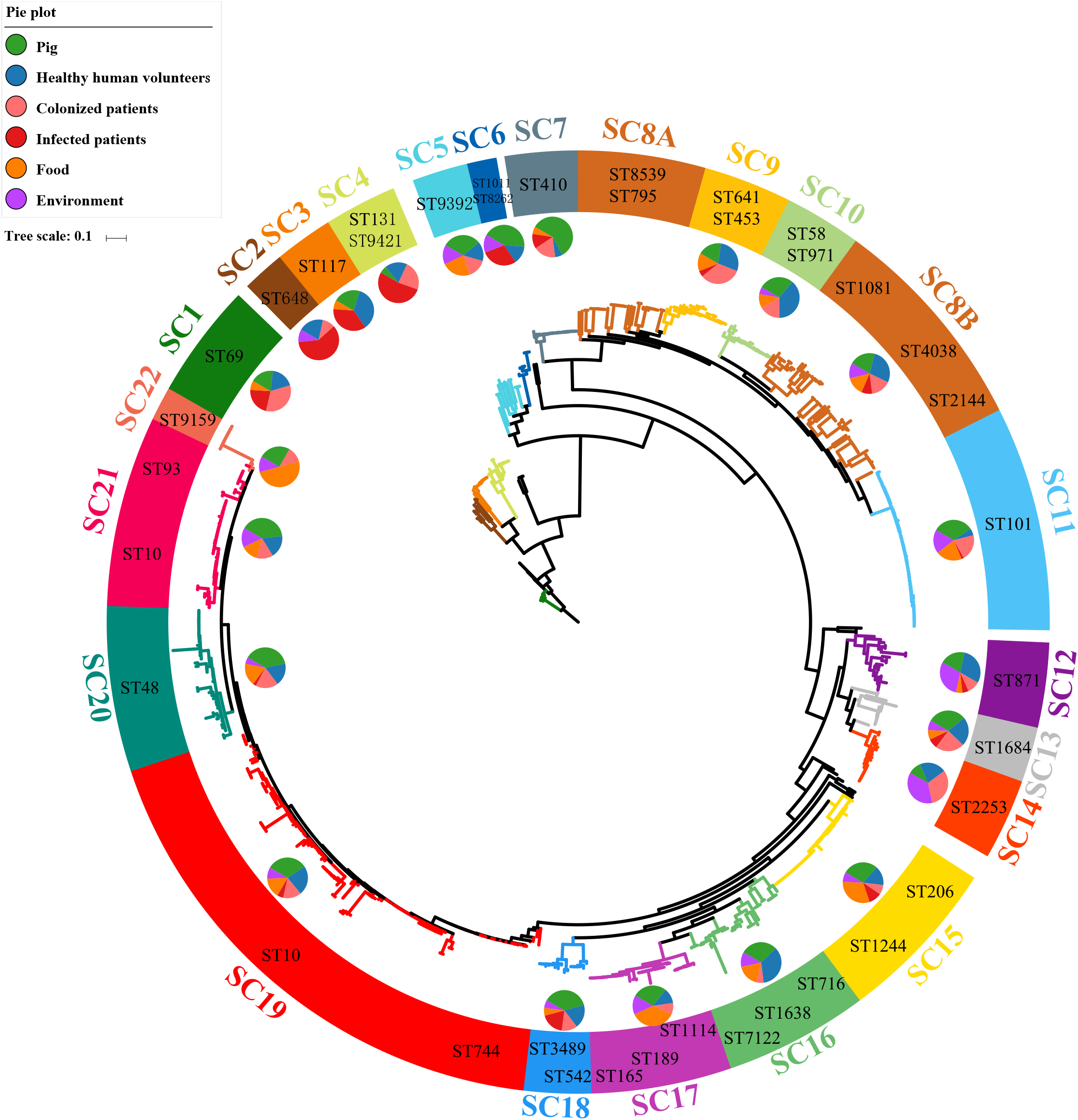
Population structure of mcr-1-producing E. coli (MCRPEC).

**Figure 5.**
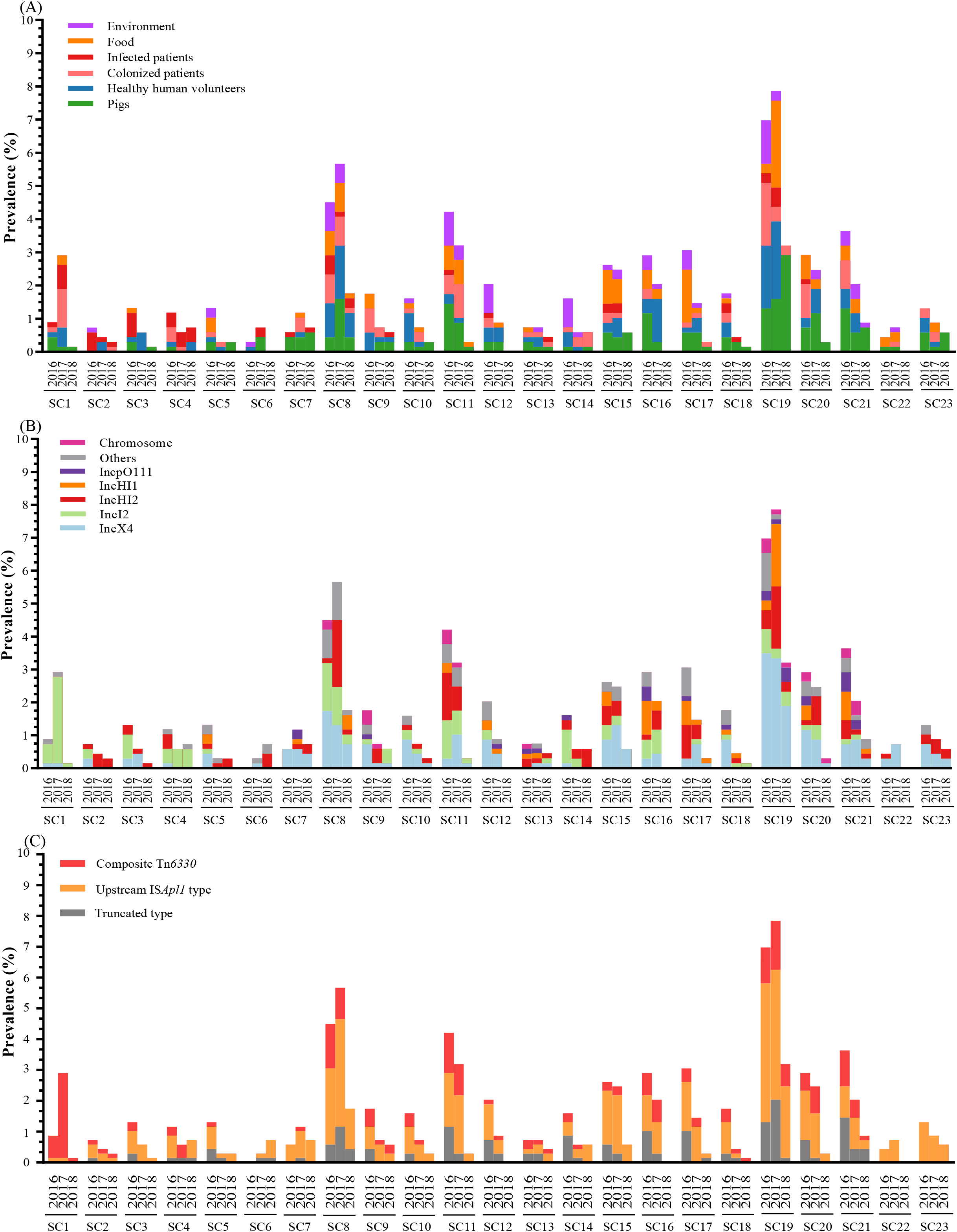
Distribution of (A) sources, (B) plasmid Inc types and (C) mcr-1-associated flanking sequences for each mcr-1-producing E. coli (MCRPEC) sequence cluster (SC) by sampling year.

The frequency of occurrence for most accessory genes across isolates was low, with 26009 (75·6%) of 34388 accessory genes found in <5% MCRPEC and 8781 (25o5%) of 34388 in only one isolate. We included 5700 accessory genes which were contained in 5%-95% of MCRPEC for pan-genome analysis. Intersection analysis showed that 230/5700 accessory genes were lacking in MCRPEC causing infection and 110/5700 accessory genes were lacking in MCRPEC from environmental samples (figure 6A). Network analysis showed that the included 5700 accessory genes was not significantly correlated with source/host and that MCRPEC accessory genes were shared amongst most hosts, except for MCRPEC isolates causing infection which clustered in SC1-SC4 (figure 6B). Moreover, enrichment analysis showed that 5·5% (313/5700) accessory genes or gene clusters were enriched in infection samples (Benjamini-Hochberg-adjusted p<0·0001), whereas 97/5700 (1·7%) were enriched in food samples, only one gene in environmental samples, and none in the other sources (figure 6C).

**Figure 6.**
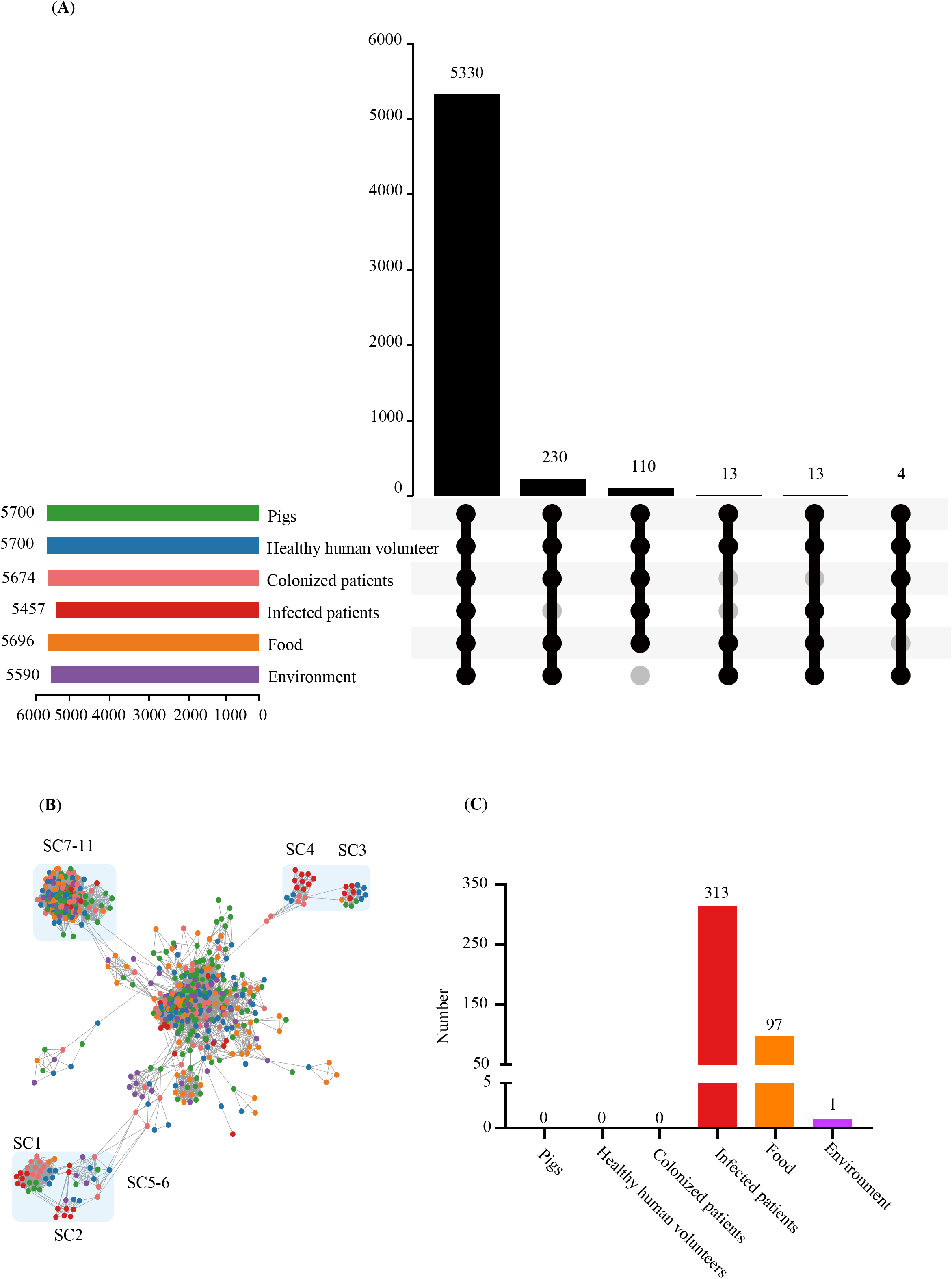
(A) Accessory genes by source for 688 *w*c*/o-/*-producing E. coli (MCRPEC), (B) per-isolate accessory gene network analysis and (C) accessory gene enrichment in different sources. (A) Upset plot of intersection of 5700 accessory genes by isolate source. The total number of accessory genes by source is represented in the left barplot. The intersection plot on the right demonstrates the number of genes shared across a subset of source categories. (B) Network graph of pairwise distances of accessory gene content between isolates. Each node represents an isolate, color-coded to indicate the isolate source. Edges indicate ?50% shared accessory gene content; edges representing <50% shared accessory gene content were removed. (C) The histogram represents the number of enriched accessory genes for each source (p<0·000/).

Maximum likelihood phylogeny of 688 MRCPEC inferred from an alignment of 2290 core COGs including 43648 polymorphic sites. Clades are colored by sequence clusters (SCs; derived from hierBAPS) on the outside ring; SC8 is further divided into monophyletic sub-clusters A, and B. SC23 was comprised of low-frequency genotypes and not show in the figure. Pie plots represent the source distribution of isolates for each SC; for SC8A and SC8B, this has been represented as part of SC8B. Dominant MLST types for each SC are labeled.

Sequence clusters (SCs) are derived from the phylogenetic analysis summarized in Fig.4, and represented along the x-axis. Clustered bars represent proportions of isolates in each category, namely isolate source (panel A), plasmid Inc type (panel B) and mcr-1-associated flanking sequences type (C), by sampling year. SC8 and SC23 are polyphyletic. SC8 has two distinct sub-clades, and SC 23 includes polyphyletic lineages represent at minor frequencies in the population.

### Genetic contexts and dynamics of *mcr-1*

Using our approach to plasmid Inc typing from the genomic data and S1-PFGE with Southern blot, we assigned the location of *mcr-1* in 656/688 MCRPEC isolates, with *mcr-1* present in a plasmid context in 632 MCRPEC and in a chromosomal location in 24 MCRPEC (figure 7, appendix figure 6). *mcr-1*-harbouring plasmids in 632 MCRPEC represented ten Inc types including IncX4 (36·9%, n=233), IncI2 (21·4%, n=135), IncHI2 (20·4%, n=129), IncHI1 (9·3%, n=59), IncF (4·6%, n=29), IncpO111 (4·1%, n=26), IncP (1·9%, n=12), IncY (1·3%, n=8), IncHI2+IncX4 (0·6%, n=4), and IncI2+IncX4 (0·2%, n=1) (figure 7, appendix figure 6).

**Figure 7.**
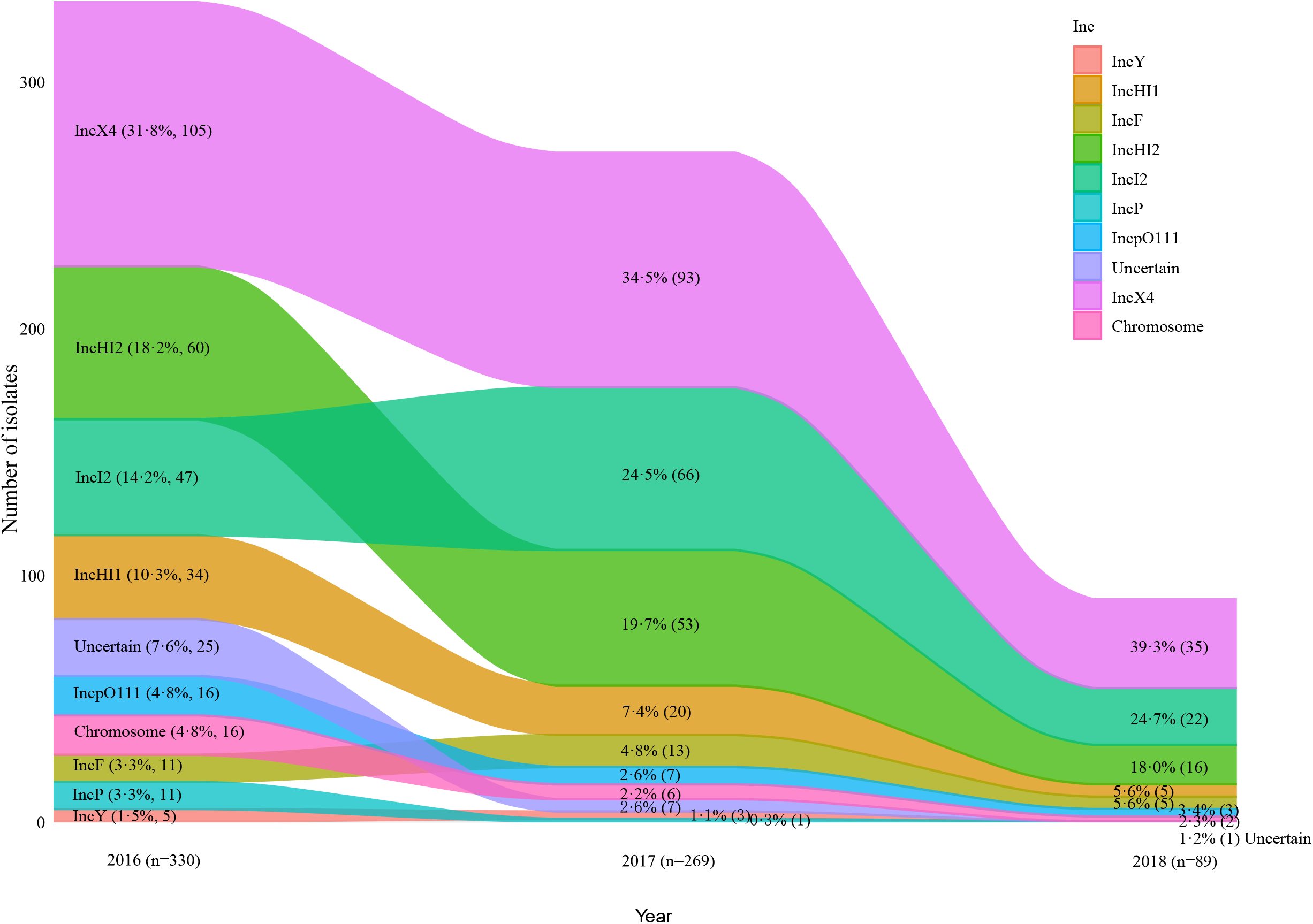
Dynamics of the genetic context of *mcr-1* in 688 sequenced *mcr-1*-producing E. coli (MCRPEC) by sampling year.

IncX4, IncI2 and IncHI2 plasmids were the predominant location for *mcr-1*, accounting for 76·5% (502/656) MCRPEC cases. These plasmids were prevalent in various hosts and STs, indicating that they highly adaptive and mobilisable plasmid vectors of *mcr-1* transmission (figure 7 and appendix figure 7). The proportion of MCRPEC with IncX4 and IncHI2-associated *mcr-1* did not change significantly over the three-year study period. Compared with 2016 (47/330 [14·2%]), however, the proportion of IncI2-associated *mcr-1* was significantly higher in 2017 (66/269 [24·5%], Bonferroni-adjusted p=0.0014) and 2018 (22/80 [24.7%], Bonferroni-adjusted p=0·018) (figure 7). Notably, when we pooled other types of the *mcr-1* gene locations, their prevalence decreased significantly after the intervention (2016 *vs* 2017, Bonferroni adjusted p=0.0061). In addition, the diversity of Inc types associated with *mcr-1* decreased after the colistin ban (e.g. loss of IncP and IncY) (figure 7). These results indicated that the intervention may have influenced non-adaptive plasmids more than adaptive plasmids harboring *mcr-1* (figure 7). We also analyzed the distribution of *mcr-1* location by source, and the results showed that IncX4 was the dominant *mcr-1* plasmid type for most of the sources and STs; IncHI2 however was the dominant *mcr-1* plasmid type for MCRPEC causing infection (25 [45·5%] of 55) (appendix figure 7).

Alluvial diagram representing proportional changes in the genetic context (chromosome, plasmid type, uncertain) of *mcr-1* in sampled MCRPEC over time. Y-axis shows the number of isolates. Numbers in brackets for each category represent the percentage and number of isolates for each *m*c*r-1*-associated genetic context by year. The “Uncertain” category includes five isolates which harbored two *mcr-1* plasmids of different Inc types.

To more fully analyze the structures of *mcr-1*-harbouring plasmids, we extracted the coding sequences (CDSs) and performed CDS clustering analysis for the three dominant types of *mcr-1* plasmids, namely IncX4, IncI2 and IncHI2 (figure 8). We obtained 79 CDSs among 232/233 analyzable IncX4 plasmids (median: 40 [range: 35-47] CDSs per plasmid), 134 CDSs among 135 IncI2 plasmids (median 77 [range: 68-91]), and 645 CDSs among 124/129 analyzable IncHI2 plasmids (median 243 [range: 156-286]). Clustering results demonstrated that IncX4 plasmids were generally stable, compared with both IncHI2 and IncI2 plasmids, which were more diverse (figure 8). Interestingly, we found 20/124 IncHI2 plasmids and 34/135 IncI2 plasmids had a specific lineage lacking ~50 CDSs in IncHI2 and ~15 CDSs in IncI2, respectively (figure 8 and table 3). Further analysis indicated that some of absent CDSs are well-known to involve the components of type IV secretion system (T4SS) and play a role in plasmid transformation, such as *tra, dsbC*, and *mauD* in IncHI2, and *virB, pilR* and *hicAB* in IncI2 (table 3). Therefore, we performed conjugation experiments for IncHI2-carrying isolates and IncI2-carrying isolates with/without these specific CDSs. Surprisingly, all 20 IncHI2 plasmids from the lineage lacking these CDSs failed in to conjugate, whereas 13/15 randomly selected IncHI2 plasmids from other lineages successfully conjugated (figure 8). We also noted that the proportion of IncHI2 plasmids without these 50 CDSs were significantly increased after the colistin ban (4/62 [6·5%]) in 2016 vs 16/71 [22·5%] in 2017/2018). These results indicate that the transferability of this IncHI2 plasmid in post-ban MCRPEC may be diminished, which could lead to restricted spread of this plasmid. These findings were not observed for the IncI2 plasmids, where the presence/absence of the 15 CDSs had no discernible impact on conjugation efficiency (19/34 [55·9%] IncI2 plasmids without the CDSs successfully conjugated vs 20/43 [46·5%] plasmids with the CDSs; [p=0·4141]), which indicated that conjugation ability in IncI2 plasmids was most likely unrelated to these CDSs.

**Figure 8.**
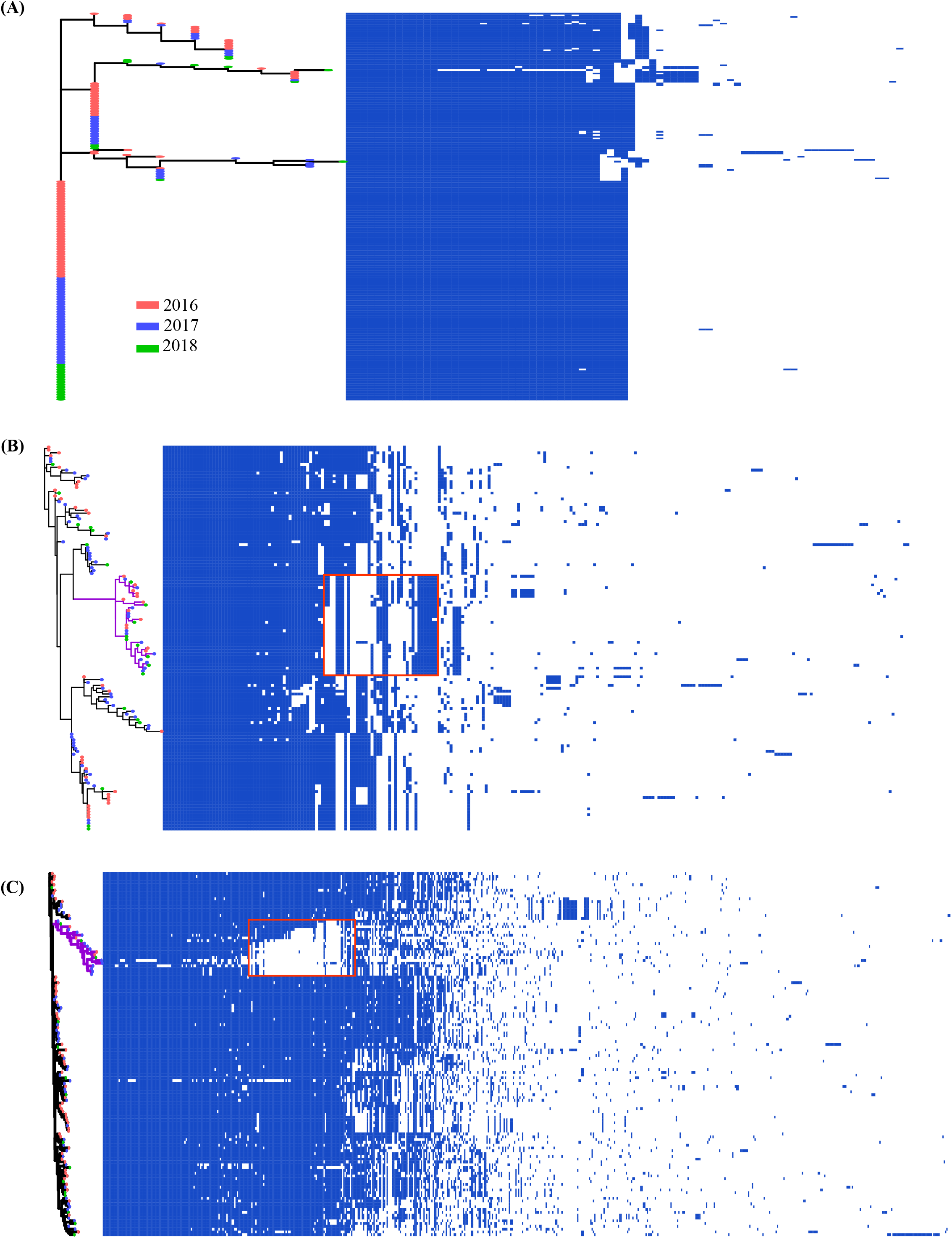
Clustering analysis of coding sequences (CDSs) for (A) IncX4, (B) IncI2 and (C) IncHI2 plasmid.

**Table 3.** CDSs present/absent in IncHI2 and IncI2 type plasmids pre-and post-colistin ban

| Location | Status | CDS | COG annotation |
| --- | --- | --- | --- |
| IncHI2 | Absent | *traD* | TraM recognition site |
|  |  | *traG* | TraM recognition site |
|  |  | *traG* | TraG protein |
|  |  | *traH* | TraH Conjugative relaxosome accessory transposon protein |
|  |  | *dsbC* | Required for disulfide bond formation in some periplasmic proteins.<br>A dedicated oxidative folding enzyme is important for conjugative plasmid transfer. |
|  |  | *mauD* | Plasmid transfer operon protein |
|  |  | *HNH* | HNH nucleases |
|  |  | *Mrr* | Mrr N-terminal domain |
|  |  | Others | Forty-two Hypothetical protein |
| IncI2 | Absent | *trbL/virB6* | TrbL/VirB6 plasmid conjugal transfer protein |
|  |  | *hicA* | HicA toxin of bacterial toxin-antitoxin |
|  |  | *hicB* | HicB_like antitoxin of bacterial toxin-antitoxin system |
|  |  | *pilR* | Protein transport across the cell outer membrane |
|  |  | *xseA* | exodeoxyribonuclease VII activity |
|  |  | *dnaJ* partial | DnaJ molecular chaperone homology domain |
|  |  | Others | Nine hypothetical protein |
|  | Present | *cbix* | cobalamin biosynthesis protein CbiX |
|  |  | Others | Six hypothetical proteins |

The CDSs in this table presents the marked area of gene presence/absence for figure 7.

The CDS presence/absence dendrogram for MCRPEC isolates harboring mcr-1 on (A) IncX4, (B) IncI2 and (C) IncHI2-associated plasmids, representing CDSs extracted from draft plasmid sequences. The red rectangles highlight plasmid clusters lacking distinct groups of CDSs, as discussed in the text and table 3. Purple branches on the phylogeny represent the lineages affected. Tip colors represent sampling year.

Because recombination of transposons enables *mcr-1* to transfer across plasmids and isolates, we then focused on the *mcr-1* flanking sequences (figure 9 and appendix figure 8). We extracted the composite transposon *Tn6330* or flanking sequences of *mcr-1* from each of the 688 sequenced MCRPEC to clarify the genetic context of *mcr-1*, and any changes over time. Tn*6330* (IS*Apl1-mcr-1-pap2-ISApl1*) has been thought to be the primary vehicle for transmission of *mcr-1* ^23–25^. We found that in 133 MCRPEC (19·3%, n=394) *mcr-1* was part of a complete Tn*6330*, whilst 161 (23·4%) contained only the upstream *ISApl1*, and the majority (57·3%) contained no IS*Apl1* (“truncated type”) (figure 9), indicating the likely importation and stabilization of *mcr-1* differently in different genomic backgrounds.^23,25^ Furthermore, we found that 113/129 (87·6%) IncHI2 plasmids harbored IS*Apl1* upstream of *mcr-1* (i.e. *ISApl1-mcr-1-pap2)*, while only 5/233 (2·1%) IncX4 and 19/135 (14·1%) IncI2 plasmids contained IS*Apl1* upstream (appendix figure 8). Of note, we found that percentage of IS*Apl1*-lacking MCRPEC increased significantly from 2016 (174/330 [52·7%]) to 2018 (67/89 [75.3%]), Bonferroni-adjusted p=0.0005) (figure 9). Also, we noticed that the upstream type *(\SApl1-mcr-1-pap2)* decreased in 2018 only, which suggests that the Tn*6330* composite transposon firstly lost its downstream *ISApl1*, becoming a truncated composite transposon (figure 9B). In addition, no difference was observed in distribution of Tn*6330* among sources (appendix figure 8).

**Figure 9.**
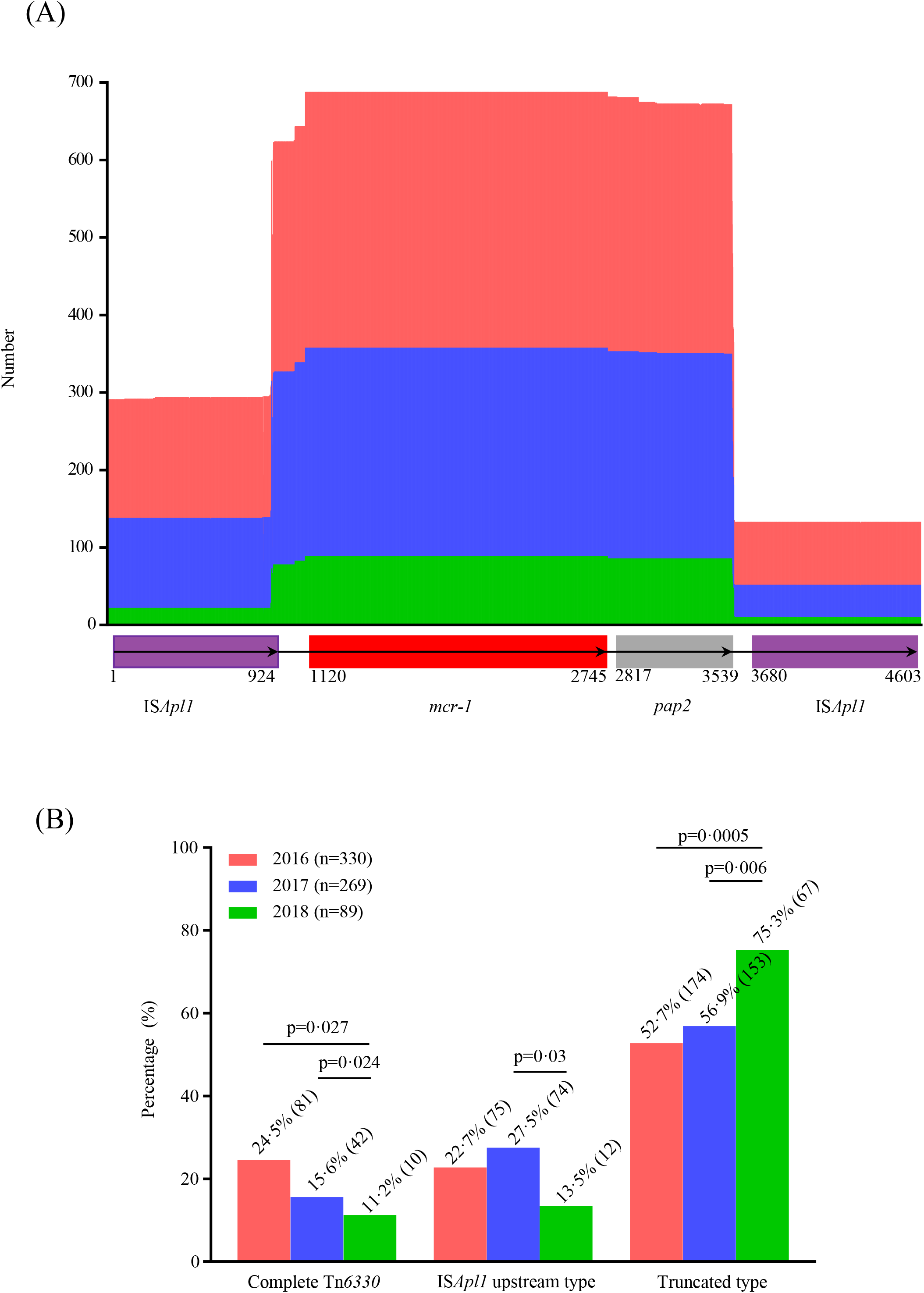
The structural dynamics of the Tn*6330* genetic element carrying *mcr-1* by sampling year. (A) Length distribution of the sequence alignments with respect to the reference *Tn6330* sequence (x-axis) across the three sampling years. (B) Histogram of prevalence for each type of *Tn6330* element (complete, upstream *ISApl1* sequence only, truncated type [i.e. no *ISApl1]*) across the three sampling years.

### The frequency of cgSNPs and accessory genes before and after the colistin ban

As seen above, both the prevalence and distribution of specific resistance genes, transposons and plasmids associated with *mcr-1* in MCRPEC were significantly altered after the banning of colistin use as a feed additive. We postulated that this intervention might also affect distributions of cgSNPs and other accessory genes, and therefore, performed genome-wide association analyses to investigate the frequency changes for both cgSNPs and accessory genes before and after the intervention (2016 *vs* 2017 and 2018) (figures 10–11). We found that the frequencies of 63 cgSNPs, located on 33 genes, were significantly different pre-and post-intervention (Benjamini-Hochberg-adjusted p<0·05 and empirical p<0·05) (figure 10 and table 4). Among them, 7 SNPs, located on *dnaJ, nhaA, hycD* and *yjjW* genes, respectively, led to missense mutations (figure 10 and table 4). Specifically, *dnaJ* is known to be associated with plasmid DNA replication, *nhaA* is associated with the adaptation to high salinity at alkaline pH, *hycD* has an oxidoreductase activity, and *yjjW* encodes a pyruvate formate-lyase activating enzyme. Of 5700 accessory genes, 30 genes or gene clusters were significantly differently distributed pre-and post-intervention (figure 11 and table 5). Remarkably, 35/49 genes were associated with either plasmid maintenance or transfer (table 5). In addition, 23/30 genes and gene clusters were known to be located on IncI2 plasmids and around *mcr-1* (figure 11B and table 5). The other six genes and one gene cluster had no specific genetic location; two of these were found with higher frequency before the intervention and four of them after the intervention (figure 11A and table 5).

**Figure 10.**
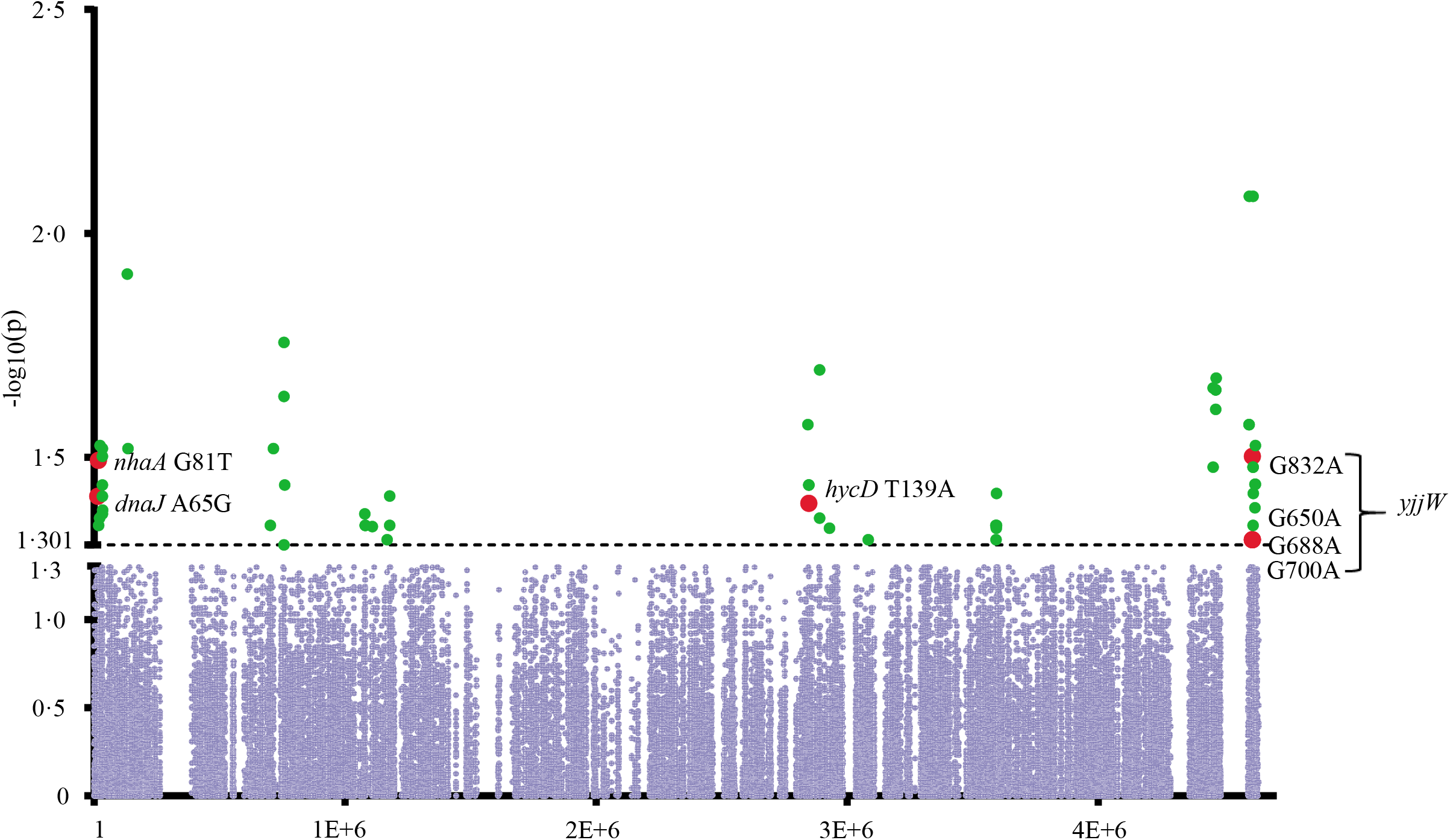
Manhattan plot for the cgSNP genome-wide association analysis of pre-and post-colistin ban mcr-1-producing E. coli (MCRPEC).

**Figure 11.**
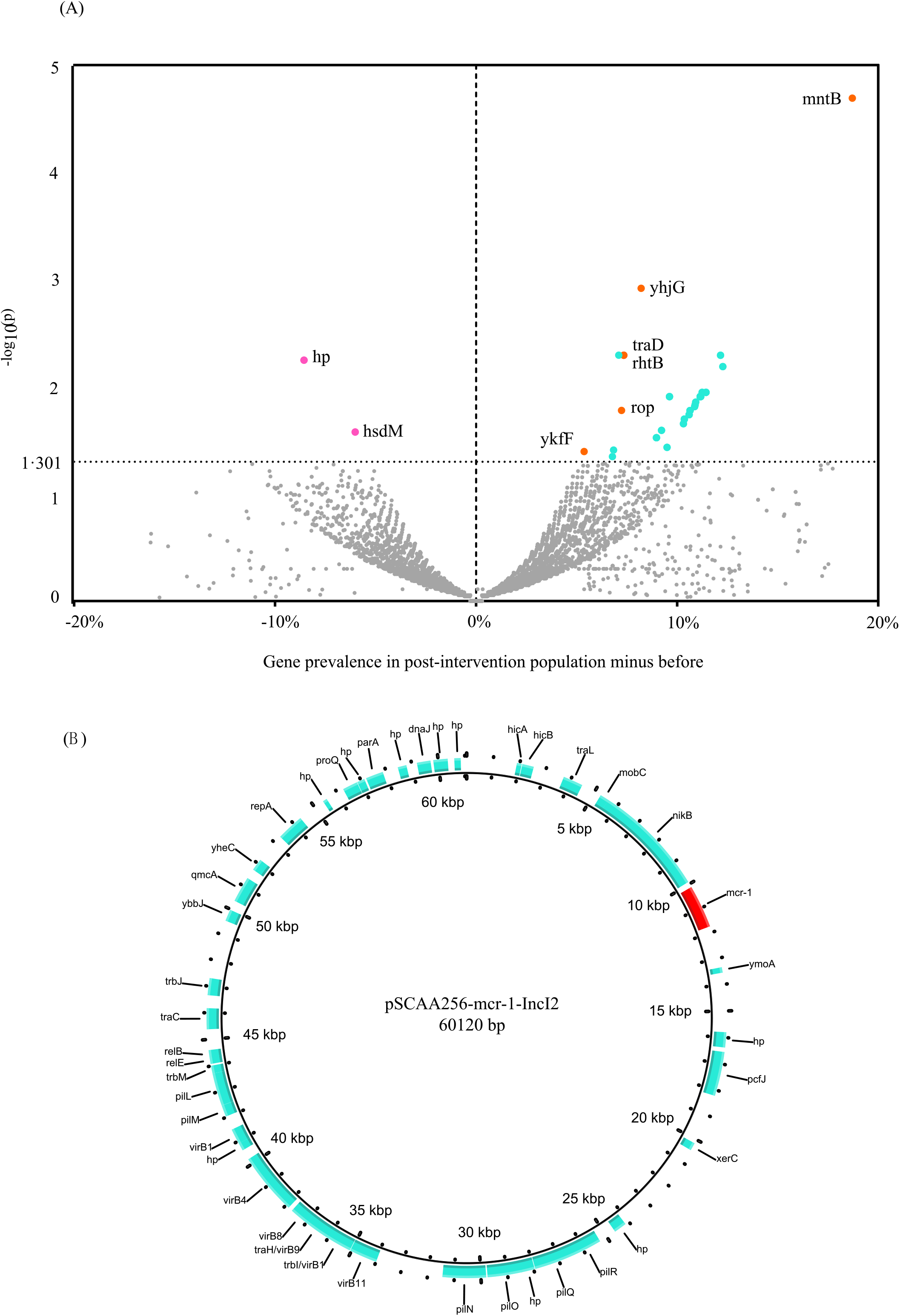
Genome-wide association analysis results of accessory genes for pre-and post-colistin ban *mcr*-/-producing *E coli* (MCRPEC). (A) Volcano plot of different accessory genes between pre-and post-colistin ban MCRPEC populations. The y-axis represents the −log10(p)-values derived from the genome-wide association analysis. The x-axis represents the accessory gene prevalence of the post-ban population minus the gene prevalence pre-ban. Blue dots represent genes located on IncI2 plasmids. Orange dots with text annotation represent genes not thought to be located on IncI2 plasmids. (B) Circle map of a representative mcr-1-harboring IncI2 reference plasmid. Blue CDSs match blue dots in figure 10A.

**Table 4.**
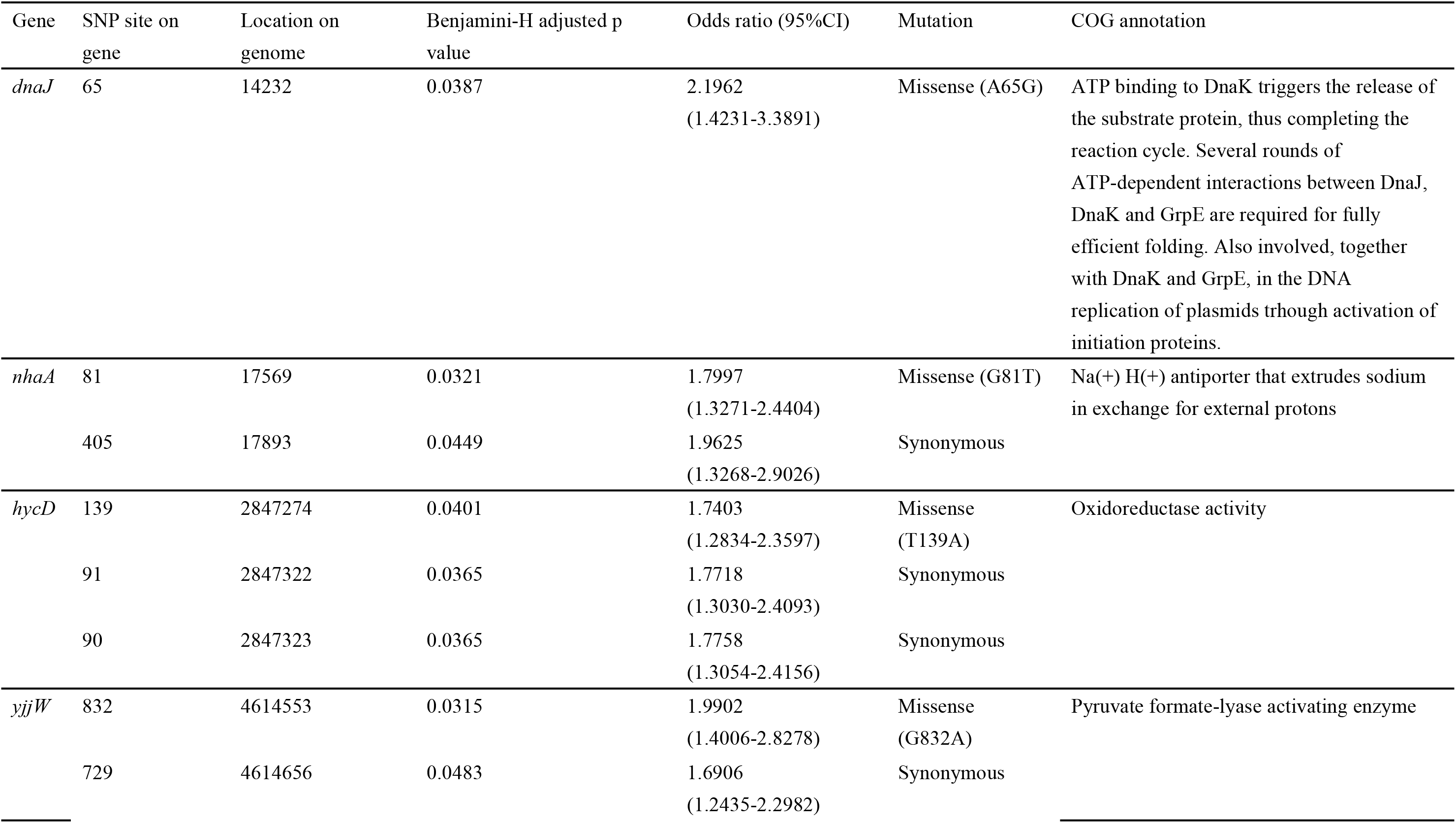
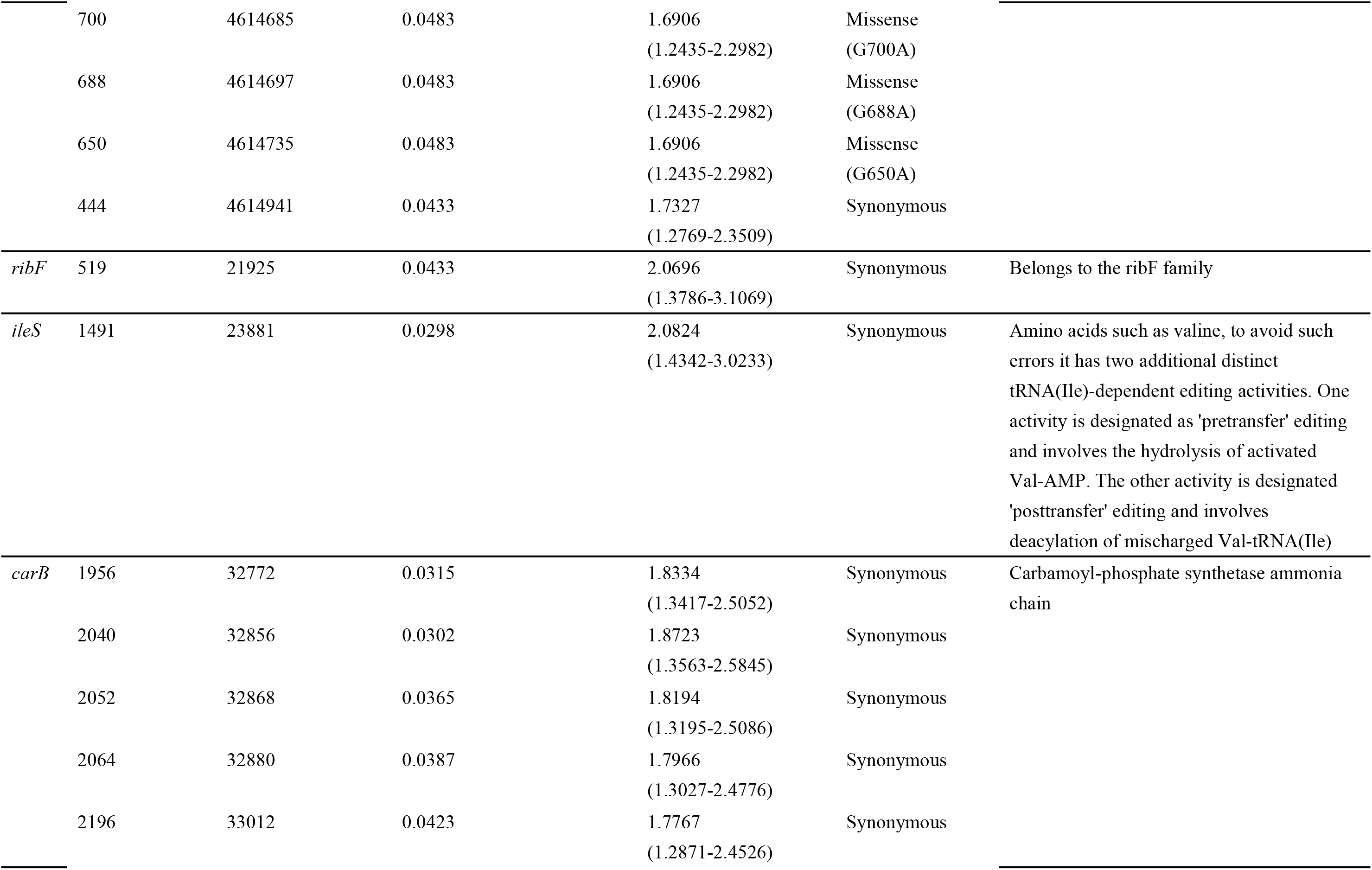
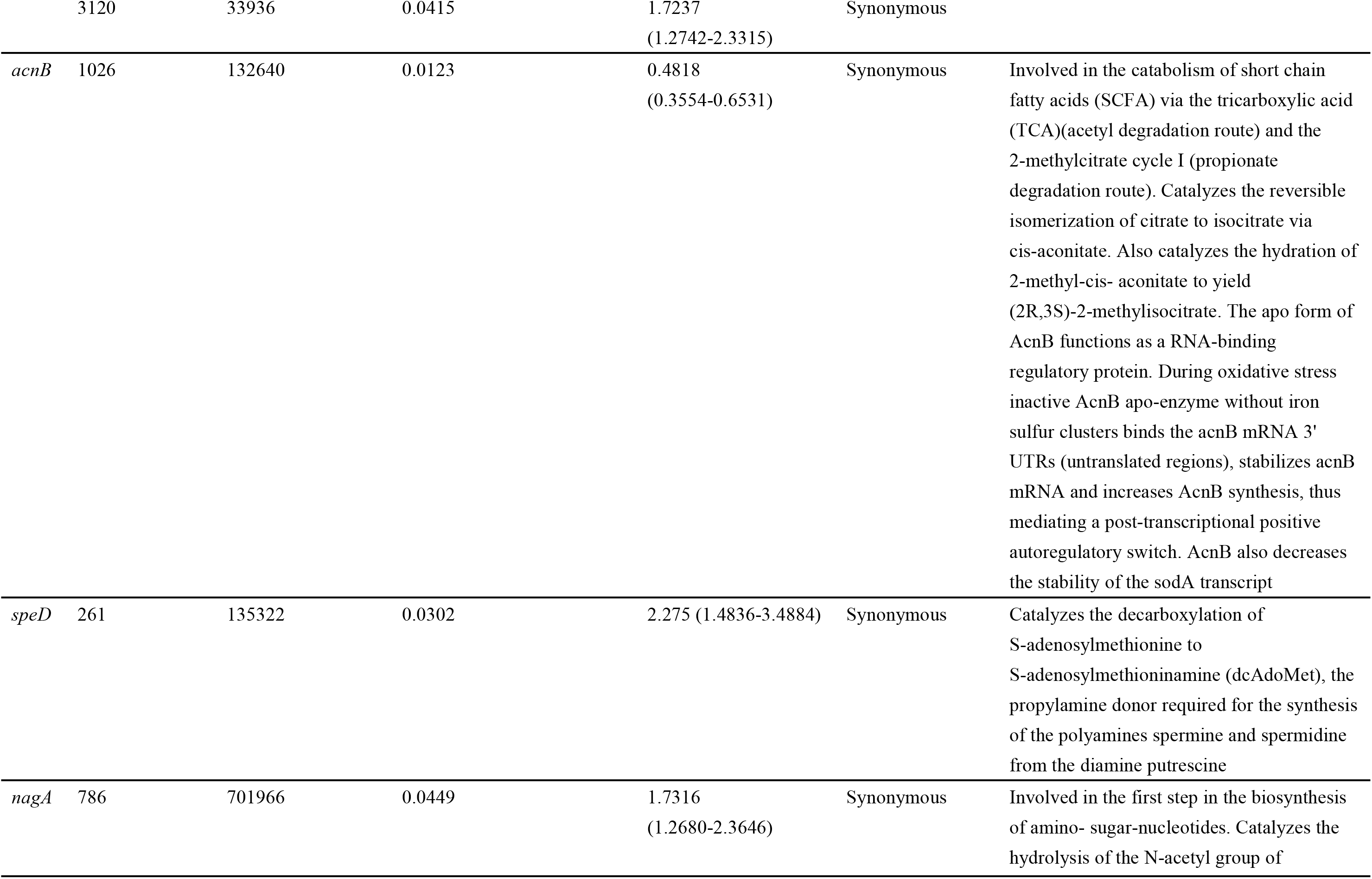
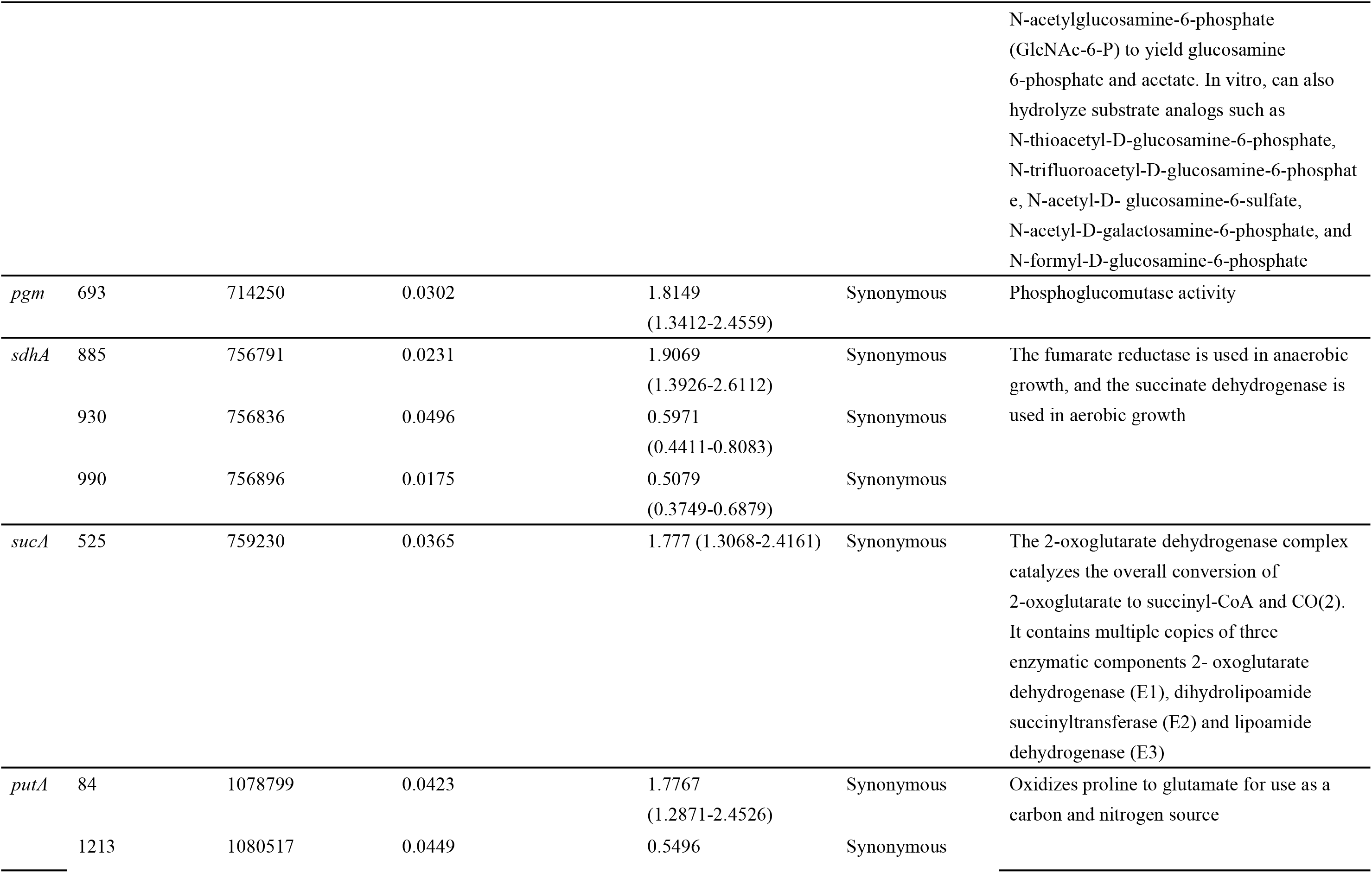
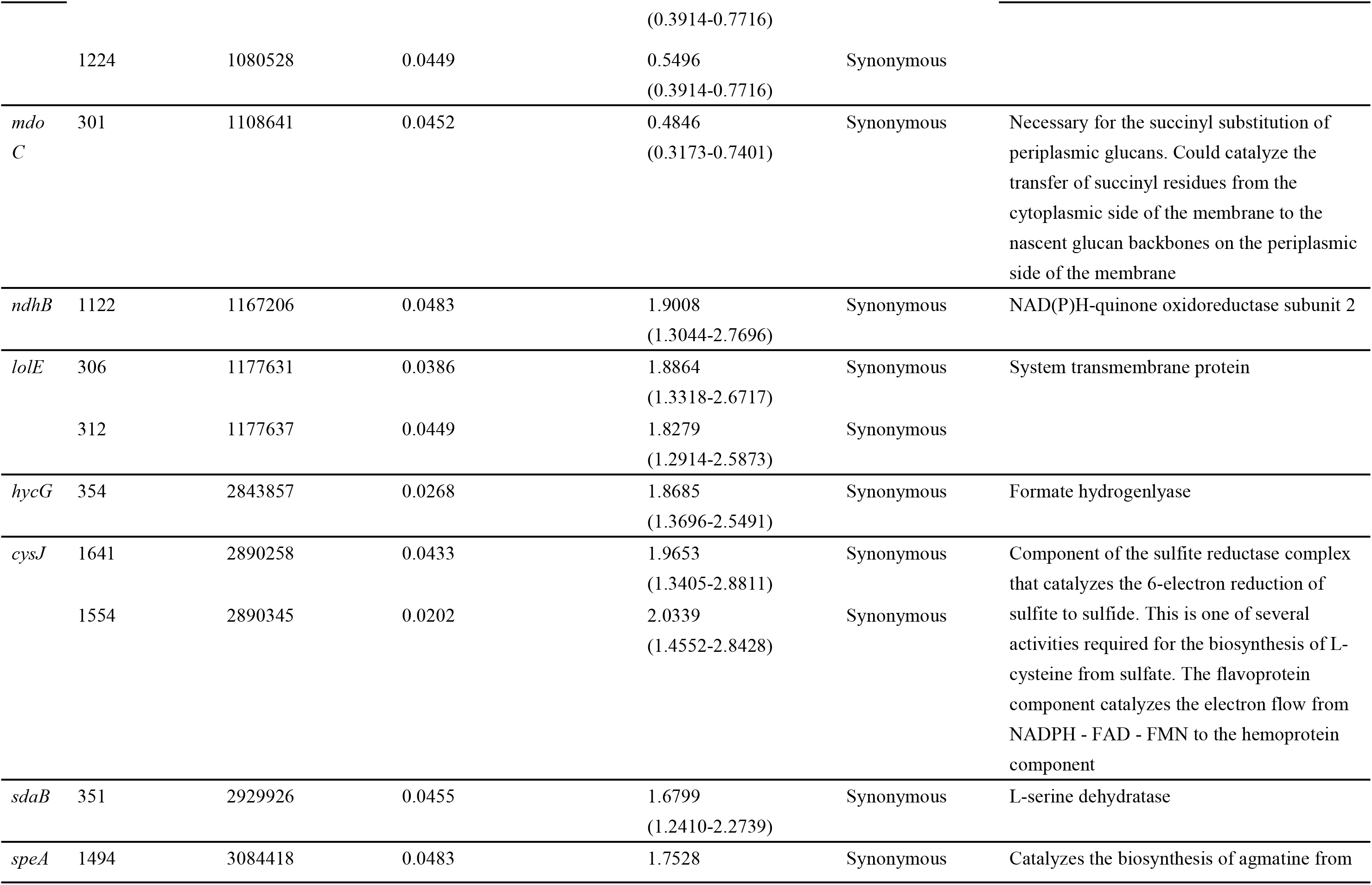
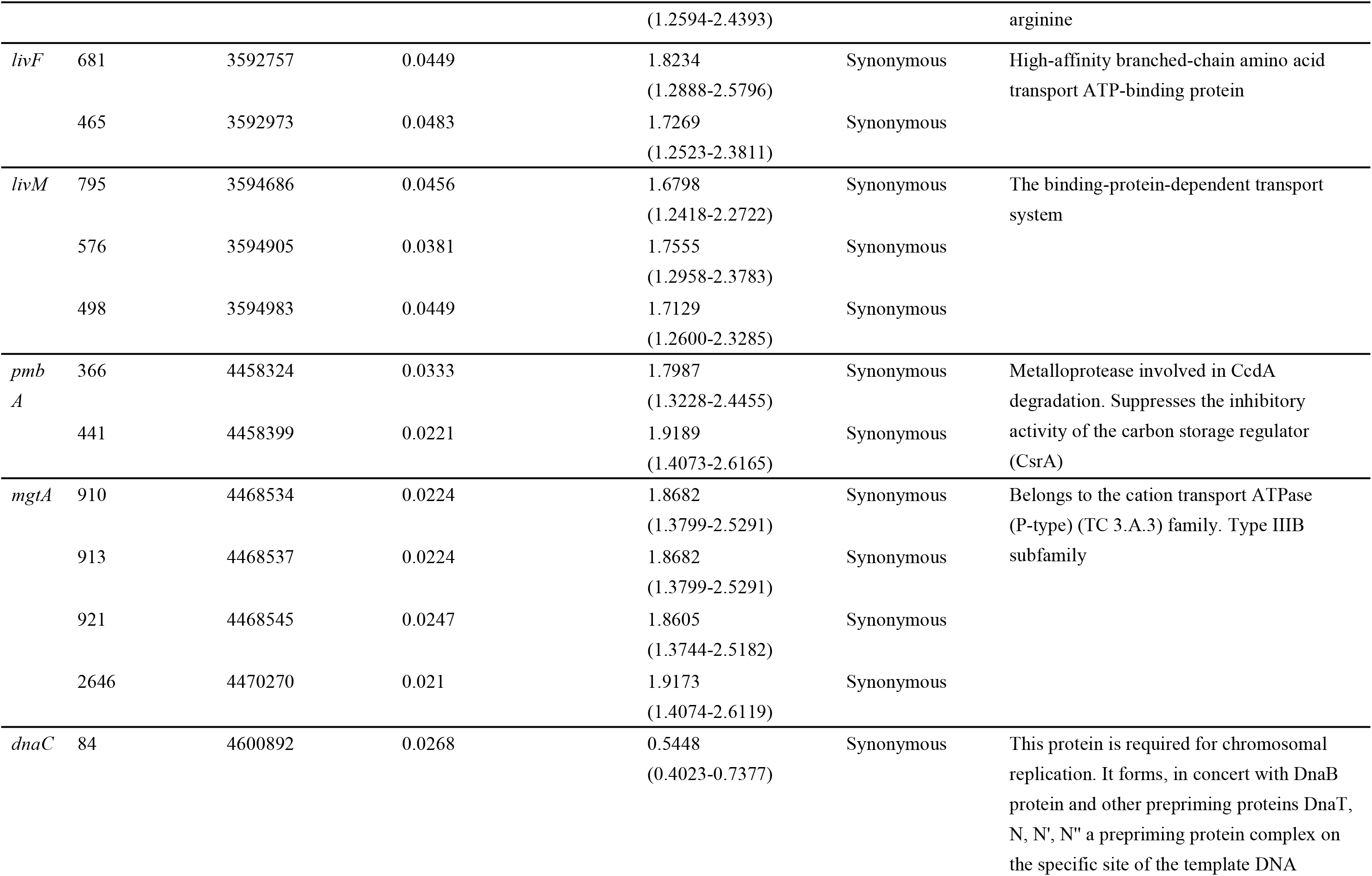
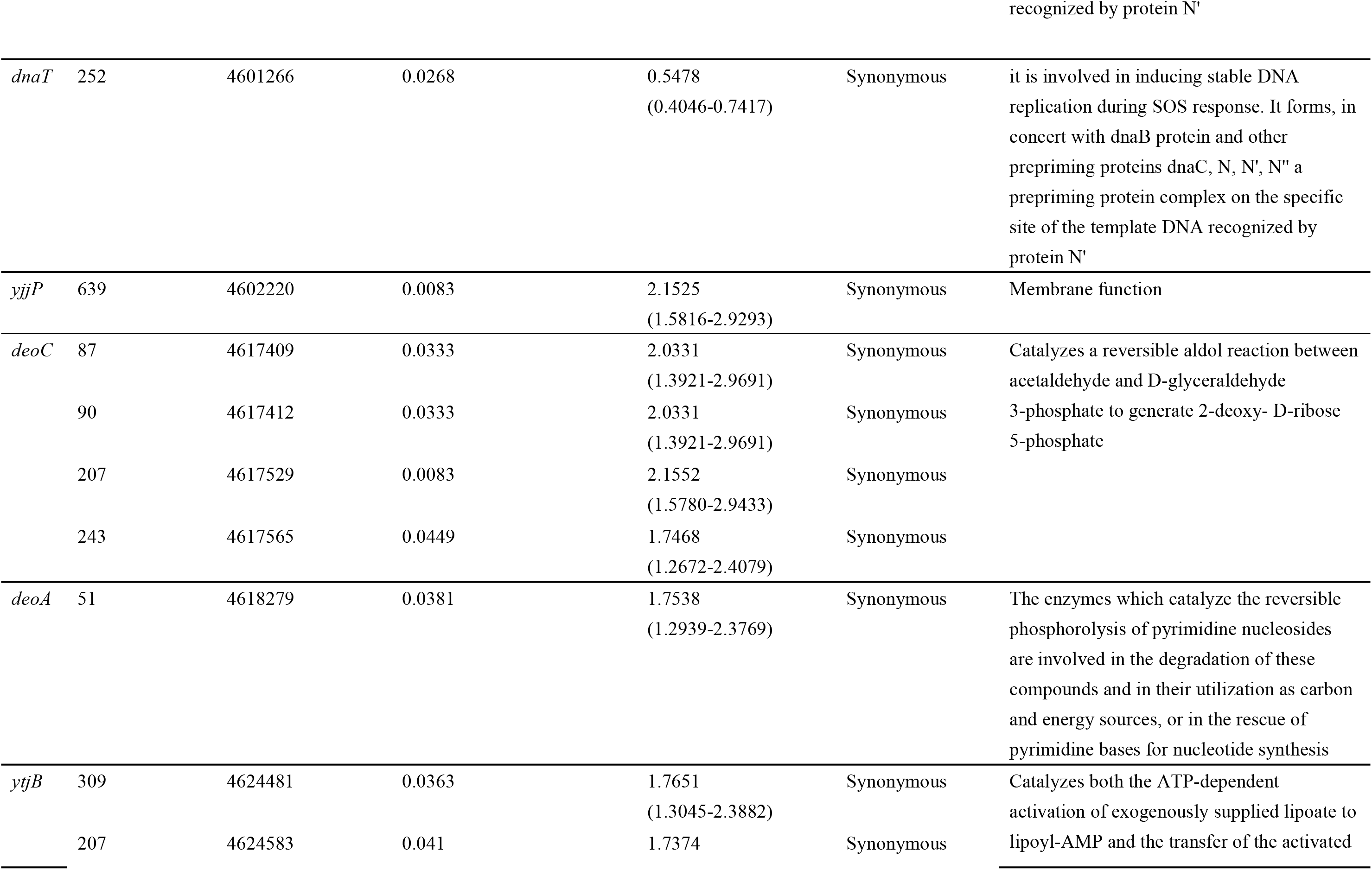
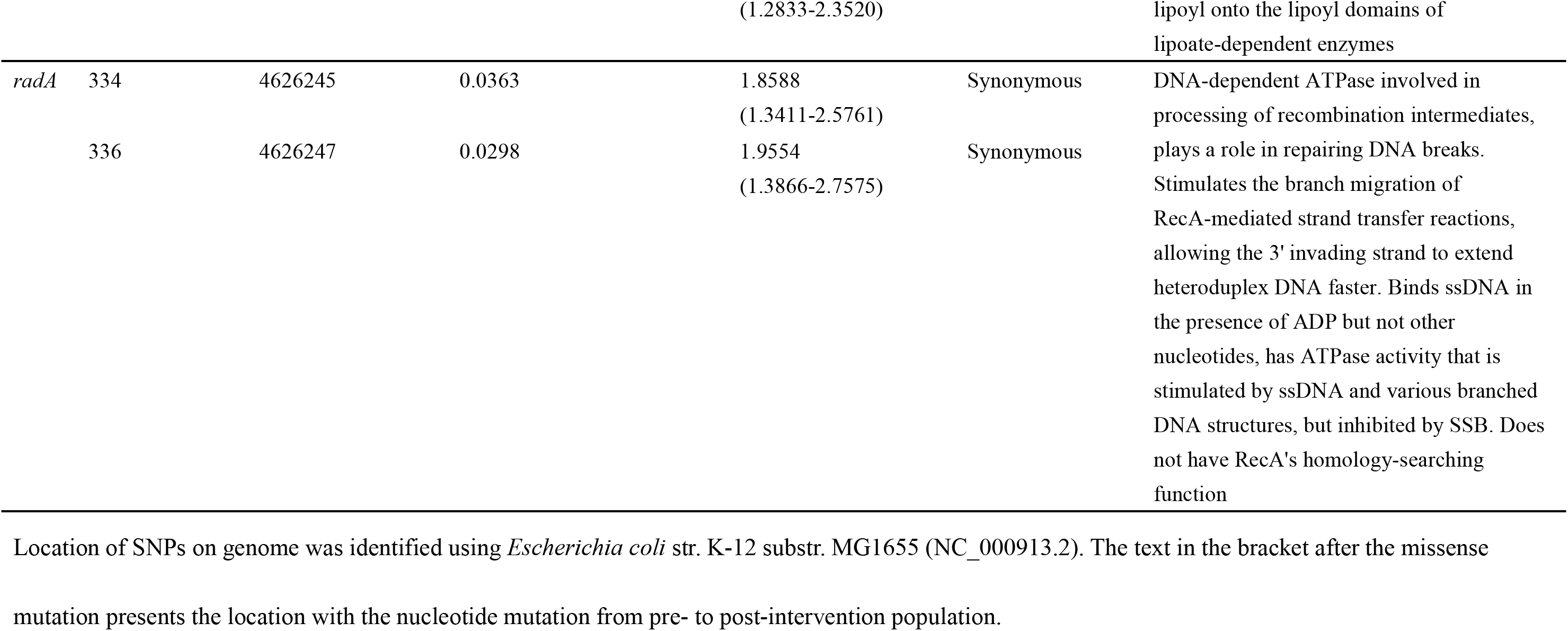
Genetic information on significantly different cgSNPs identified in MCRPEC pre-and post-colistin ban.

**Table 5.**
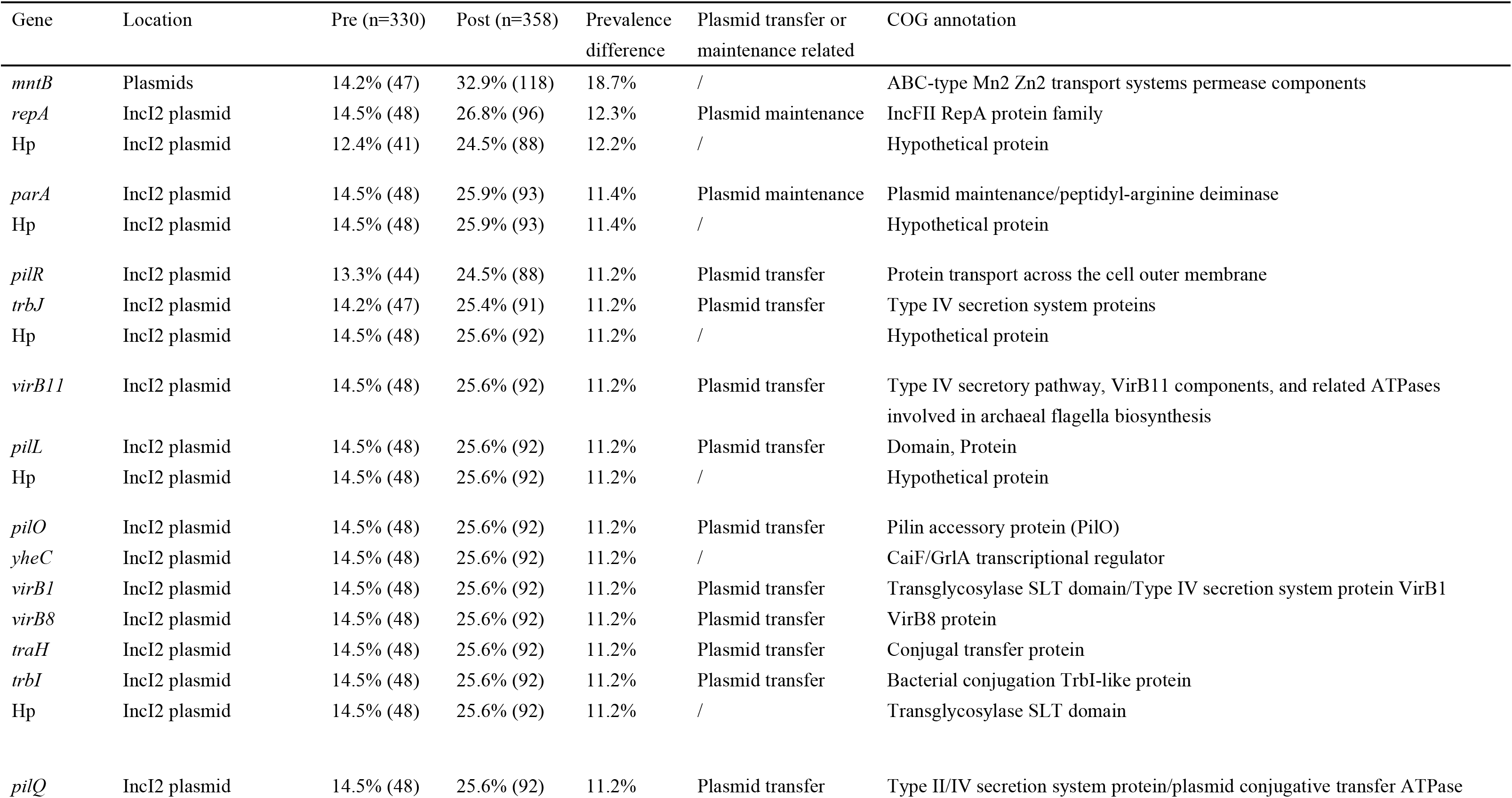
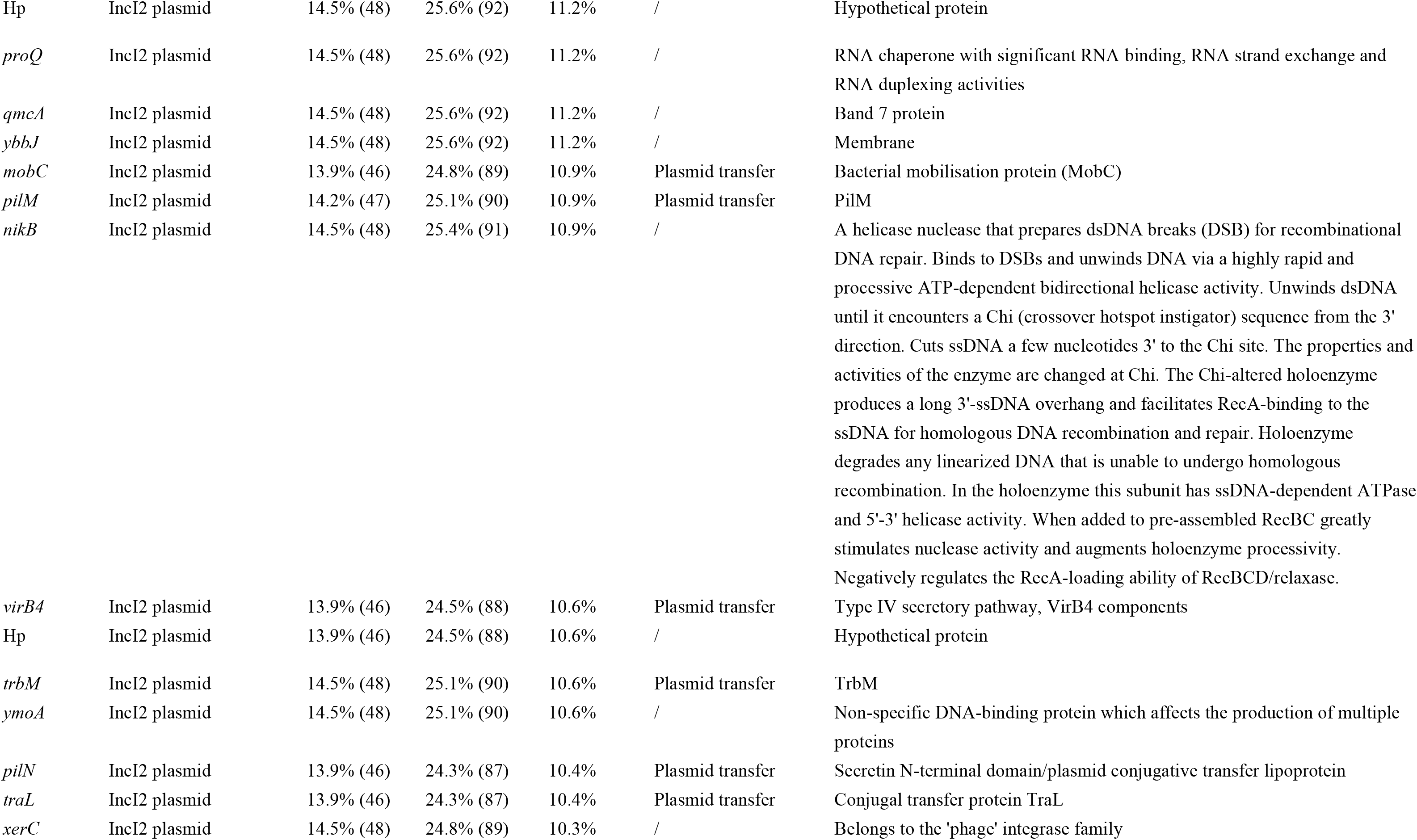
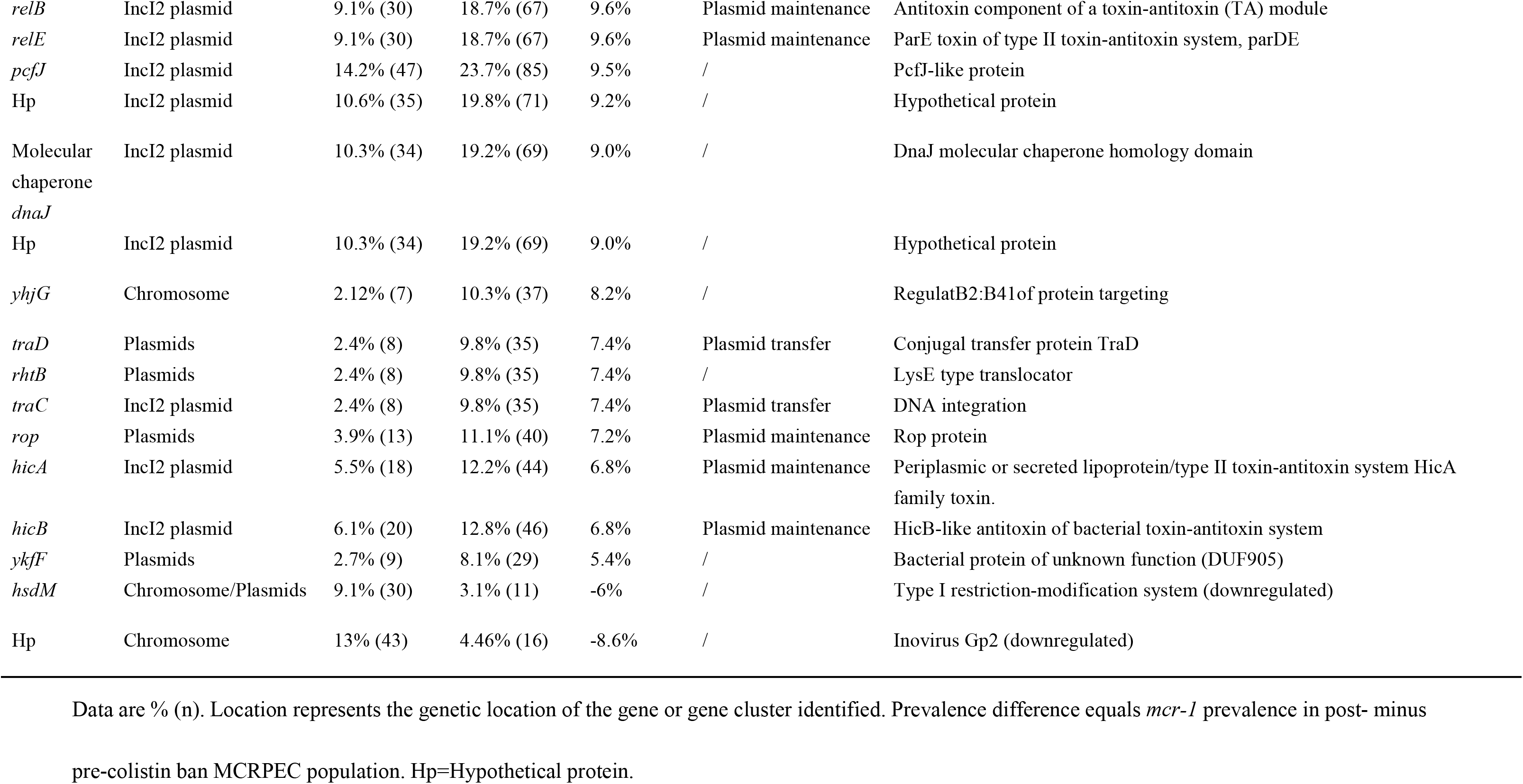
Description and annotation of differential accessory gene or gene clusters between pre-and post-intervention.

Dots represent cgSNPs. The y-axis represents log10(p)-values derived from the genome-wide association analysis. The x-axis represents nucleotide positions with respect to the reference strain Escherichia coli str. K-12 substr. MG1655 (NC_000913.2). The plot above the dashed horizontal line (y=1.301) represents cgSNPs that have met the pre-specified significance threshold (Benjamini-Hochberg-adjusted p<0.05). Green dots represent cgSNPs causing synonymous mutations. Red dots with text annotation represent cgSNPs causing non-synonymous mutations, with annotations of the locus affected (where known), the nucleotide change and the nucleotide position with respect to the locus of interest.

## Discussion

Colistin has been extensively used in animals in China and other countries, and this has likely contributed to the increase and transmission of colistin resistance including the emergence of *mcr-1*.^5^ The compulsory ban of colistin use as a feed additive for animals in May 1^st^ 2017 in China afforded a clear opportunity to understand the impact of this ban on *mcr-1* prevalence and the genetic dynamics of *mcr-1*-positive bacterial populations. We show clearly that the ban appears to have led to dramatic declines in *mcr-1* prevalence in farmed pig populations nationally, and over short timeframes (two years). With our detailed study across both porcine and non-porcine associated niches in Guangzhou over the same timeframe, we have also demonstrated concomitant sharp decreases in *mcr-1* prevalence in human carriers (healthy and inpatient), slaughterhouse-associated river and environmental samples, and pork foodstuffs by 2018. Previous studies have shown that rates of *mcr-1* in various host populations/sources including pigs, humans, environment and foods in China were very much increasing prior to the colistin ban^8,10,11^ which strongly suggests that the declines we have observed are directly attributable to this intervention. In addition, our observation of the specific declines in *mcr-1* prevalence in pig-associated sources (e.g. pigs, pork, and slaughterhouse-associated soil and water samples) indicates that food animals represent the principal reservoir of *mcr-1* in this setting.

In our large, regional study, where *mcr-1* is mostly associated with *E. coli*, we have demonstrated that both clonal and horizontal transmission are likely important contributors to the dissemination of *mcr-1* across sources. Indeed, the predominant STs in our study, such as ST10, ST101 and ST48, were found in various hosts/sample types. MCRPEC from various hosts (except, perhaps, to an extent, isolates causing infection in hospitalized inpatient) did not represent distinct genetic subpopulations, which indicated that at least some MCRPEC including those recognized as widely distributed isolates can transfer and proliferate in multiple niches, also providing an opportunity for genetic exchange with a broad range of other bacterial species.

Interestingly, the *mcr-1* gene was first identified in an IncI2 plasmid, and then it was identified in different plasmid families.^3,4^ Herein, the IncX4, IncI2 and IncHI2 plasmids were the predominant *mcr-1*-harboring vectors. A recent nationwide epidemiological study in China revealed that IncX4 was the dominant *mcr-1*-harbouring plasmid in healthy humans.^26^ Remarkably, the diversity of *mcr-1*-associated plasmid replicon types decreased following the colistin ban; and the proportion of IncI2 was significantly higher after the intervention which may suggest that IncI2 could more stable than other Inc types in MCRPEC in the absence of colistin selection pressures. In addition, we noticed that *mcr-1* was located on the chromosome in 24 (3·4%) of 688 MCRPEC, which is a potential threat as this integration may confer enhanced genetic stability for AMR genes.^27,28^ Interestingly, however, we found the proportion of chromosomally-located *mcr-1* essentially halved after the colistin ban from 4.8% in 2016 to 2.3% in 2018.

Notably, our results have shown *mcr-1* in MCRPEC causing infections in hospitalized inpatients had distinguishing genetic features. In these cases, MCRPEC were condensed mainly in lineages SC1-SC4 rather than being diffusely distributed; 25/55 (45.5%) *mcr-1* genes were located on IncHI2 plasmids; and 313/5700 (5.5%) accessory genes were enriched whereas 230/5700 (4%) were absent, indicating these may represent a relatively independent population that are clonally transmitted amongst patients. In keeping with this, a nationwide colistin ban in animal feed would not be anticipated to exert any effect on *mcr-1* prevalence in this group, which was in line with our results (low prevalence, no discernible change over time). We did not survey isolates obtained from community-associated infections - this would be interesting future work.

It has been hypothesized that after the loss of one or two copies of *ISApl1* constituting the ancestral Tn*6330* composite transposon involved in *mcr-1* mobilization, *mcr-1* is immobilized in a diverse range of plasmid backgrounds where insertions and deletions are less likely.^23–25^ Herein, we found that the prevalence of complete composite Tn*6330* sequences was significantly decreased in MCRPEC after the colistin ban; and by 2018 most *mcr-1* was located on a truncated Tn*6330* variant, probably no longer capable of transposition. IS*Apl1* genes were mostly present in IncHI2 plasmids, indicating that *mcr-1* in IncHI2-type plasmids is still potentially mobilizable through the IS*Apl1-*mediated recombination or mobilization.^23^

A sharp reduction in MCRPEC population size and the genetic changes in mobile genetic elements associated with *mcr-1* following the colistin ban could be interpreted as the imposition of a population bottleneck that could lead to a variation in the gene pool of such population.^29–31^ In this study, we found that frequencies of some cgSNPs and accessory genes were higher in the post-ban MCRPEC population, and these might confer an ability to adapt to and survive following such a wholesale change in a major selection pressure. Of note, although colistin is banned as an animal feed additive, it has been approved for the treatment of human infections in China, which might result in a switch in selection from animal to human compartments, leading to the emergence of novel or residual *mcr-1*-producing populations. We are not able to anticipate new, adaptive changes that might occur over the next few years - additional surveillance would be warranted.

Our study has several limitations. First, although we have demonstrated the dramatic decline of *mcr-1* prevalence in our national survey of pig populations and across sources in our study in Guangzhou, and in association with clear genomic changes in MCRPEC following the ban, the duration of our sampling periods was limited by the considerable resource required to undertake this detailed work. We are not able to predict whether these changes will be sustained in the medium-to long-term. We sampled over a three-month period each year, and may have missed seasonal differences; we were also only able to undertake our large study across sources in a single city (Guangzhou), and our genomic and multi-source findings may not be generalizable. We realize that more wide-ranging studies involving the surveillance of *mcr-1* and other important resistance genes in *Enterobacteriaceae*, for prolonged periods, are needed to clarify the long-term effects of China’s national colistin ban. Secondly, we were not able to assess whether the reduction in *mcr-1* was mediated by completely reducing the *mcr-1*-producing population or through the loss of *mcr-1*-harbouring plasmids because we only focused on *mcr-1*-positive isolates at the expense of looking at *mcr-1*-negative strains inhabiting the same ecological niche. Further metagenomic, or culture-based investigation of our study samples would be helpful to address this gap, and is part of future work. Finally, although we found some cgSNPs and accessory genes that might be associated with the colistin ban and *mcr-1* transmission, further experimental analysis would be needed to confirm the function of these genes and causality.

Despite these limitations, our data provide an unprecedented, comprehensive understanding of *mcr-1* prevalence across potential reservoirs, its reduction in response to a national stewardship intervention, and the molecular basis for these changes amongst the most common *mcr-1-*positive bacterial species in our setting, *E. coli.* Cessation of colistin use in animal feed appears to have dramatically reduced *mcr-1* prevalence in pigs and across other niches, including humans, food and the environment, resulting in a major “One Health” effect, over short timeframes. Clonal transmission, plasmid transfer and horizontal gene transfer by IS*Apl1-*mediated recombination have all played a role in the dissemination of MCRPEC in China. Continued surveillance of the genomic dynamics of *mcr-1*-producing bacteria will generate further knowledge for optimizing the design of *mcr-1* control in the future, and our approach could potentially be used as a template for monitoring the impact of such interventions on drug resistance in *Enterobacteriaceae* more widely.

## Research in context

### Evidence before this study

Karen Tang, *et al.* reported a systematic review and meta-analysis in Jan 27, 2017 summarizing the impact of interventions reducing antibiotic use in food-producing animals on the presence of antibiotic-resistant bacteria in animals and humans.^32^ Across 94 studies evaluated, reducing antibiotic use in animals decreased the prevalence of antibiotic-resistant bacteria in animals and humans by 15-24%. These data were however limited by their heterogeneity, quality, and lack of longitudinal data. In this study, we aimed to directly test the hypothesis that a national ban on colistin use as a feed additive for animals would be effective in reducing *mcr-1* prevalence in farmed animals, and subsequently in human populations, foodstuffs and the environment. We searched in PubMed using the terms “colistin” [MeSH]/[All Fields] and “animal” [All Fields] AND (“intervention”[All Fields] OR ban[All Fields]) for articles published before Nov 25, 2019 without restriction by language of publication, and identified only one study on *mcr-1* prevalence reduction in farm-animals after a colistin intervention which focused only a small number of farms and animals over a short timeframe. No large-scale, comprehensive study of *mcr-1* prevalence and genetic dynamics across multiple niches has been published to assess the effectiveness of such an antimicrobial stewardship intervention in animal and human populations.

### Added value of this study

To our knowledge, this study is the first nationwide study to assess the effectiveness of the ban on colistin use as a feed additive for animals in pig farms in China; the first study to assess in detail the effectiveness of the intervention across different potential reservoirs, including pigs, animals, food and the environment; and the first and largest study to use whole genome sequencing to characterise the genetic dynamics of *mcr-1* in this context.

### Implications of all the available evidence

Our study provides evidence that banning the use of colistin as a feed additive in animals dramatically reduced *mcr-1* prevalence in animals, environmental samples, human gastrointestinal carriage and pork foodstuffs, with the benefits seen over short timeframes. The findings demonstrate that antimicrobial stewardship in farmed animals can have a major impact on resistance rates across human and animal populations, in this case highlighting that restricted animal use of colistin should be a key feature in controlling *mcr-1*-mediated colistin resistance in both animal and human populations. Our detailed genomic assessment has identified core and accessory genome changes associated with bacterial adaptations to the changing colistin selection pressures. The introduction of colistin for human medicinal use in China might increase colistin resistance rates in the community and hospitals. Therefore, continued surveillance of colistin resistance and *mcr-1* is warranted.

### Contributors

CS, LLZ and YQY contributed equally in this study. GBT, CS, LLZ and YD designed the study. CS, LLZ and FRM did the literature search. GBT, CS, YQY and YD wrote the manuscript, which was reviewed and edited by NS. CS, LLZ, MAEEA and FRM performed the experiments and produced the tables and figures. CS, FRM, LLZ and YQY contributed to data analysis and interpretation. All authors (except NS) contributed to sample collection and data collection. All authors reviewed, revised, and approved the final submission.

## Acknowledgement

This work was supported by the National Natural Science Foundation of China (grant numbers 81722030, 81830103, 81902123), National Key Research and Development Program (grant number 2017ZX10302301), Guangdong Natural Science Foundation (grant number 2017A030306012), 111 Project (grant number B12003), Open project of Key Laboratory of Tropical Disease Control (Sun Yat-sen University), Ministry of Education (grant number 2018kfkt01/02) and China Postdoctoral Science Foundation (grant number 2019M653192). We acknowledge the valuable contribution of Xingui Wu (Zhongshan School of Medicine, Sun Yat-sen University, Guangzhou, China), Rui Ma (The First Affiliated Hospital of Sun Yat-sen University, Guangzhou, China) and Cheng Gong (School of Public Health, Sun Yat-sen University, Guangzhou, China) in providing statistical support and guidance. NS is funded through a Public Health England/University of Oxford Lectureship.

