## appendix for "Dynamics of *mcr-1* prevalence and *mcr-1*-positive *Escherichia coli* after the cessation of colistin use as a feed additive for animals in China: a prospective cross-sectional and whole genome sequencing based molecular epidemiological study"

**Appendix table 1. The number of MCRPEC isolates for WGS and subsequent analysis**

| Source | Status | 2016 | 2017 | 2018 | Sum |
|---|---|---|---|---|---|
| Pigs (gastrointestinal carriage) | MCRPEC | 271 | 478 | 205 | 954 |
| | WGS | 81 | 69 | 73 | 223 |
| | Included | 78 | 63 | 58 | 199 |
| Healthy volunteers (gastrointestinal carriage) | MCRPEC | 239 | 106 | 8 | 353 |
| | WGS | 63 | 78 | 8 | 149 |
| | Included | 61 | 75 | 8 | 144 |
| Hospital inpatients (gastrointestinal carriage) | MCRPEC | 309 | 46 | 9 | 364 |
| | WGS | 60 | 46 | 9 | 115 |
| | Included | 60 | 41 | 9 | 110 |
| Hospital inpatients (infections) | MCRPEC | 30 | 18 | 11 | 59 |
| | WGS | 30 | 18 | 11 | 59 |
| | Included | 27 | 17 | 11 | 55 |
| Environment | MCRPEC | 53 | 40 | 1 | 94 |
| | WGS | 53 | 40 | 1 | 94 |
| | Included | 50 | 22 | 1 | 73 |
| Food | MCRPEC | 57 | 63 | 4 | 124 |
| | WGS | 57 | 54 | 4 | 115 |
| | Included | 54 | 51 | 2 | 107 |

### Appendix table 2. Antimicrobial susceptibility results for 1948 *mcr-1*-positive isolates

| Source | Year | N | CL | PB | TGC | AMP | AMC | CTX | CAZ | FEP | GEN | AMK | ETP | IMP | MEM | FOS | NIT | CIP |
|---|---|---|---|---|---|---|---|---|---|---|---|---|---|---|---|---|---|---|
| Overall | | | | | | | | | | | | | | | | | | |
| | All | 1948 | 99.8% (1944) | 99.8% (1945) | 9.5% (185) | 81.5% (1588) | 58.4% (1138) | 25.9% (504) | 7.5% (147) | 3.64% (71) | 39.4% (767) | 0.7% (13) | 0.3% (6) | 0.1% (2) | 0.2% (3) | 12.1% (236) | 4.6% (89) | 56.1% (1092) |
| Annual | | | | | | | | | | | | | | | | | | |
| | 2016 | 959 | 99.7% (956) | 99.8% (957) | 11.3% (108) | 77.2% (740) | 56.5% (542) | 26.8% (257) | 5.6% (54) | 4.0% (38) | 34.2% (328) | 1.0% (10) | 0.5% (5) | 0.2% (2) | 0.3% (3) | 14.3% (137) | 3.3% (32) | 53.4% (512) |
| | 2017 | 751 | 100% (751) | 100% (751) | 8.9% (67) | 83.2% (625) | 55.8% (419) | 21.6% (162) | 9.2% (69) | 4.0% (30) | 44.6% (335) | 0.4% (3) | 0.1% (1) | 0% (0) | 0% (0) | 10.1% (76) | 2.9% (22) | 58.9% (442) |
| | 2018 | 238 | 99.6% (237) | 99.6% (237) | 4.2% (10) | 93.7% (223) | 74.4% (177) | 35.7% (85) | 10.1% (24) | 1.3% (3) | 43.7% (104) | 0% (0) | 0% (0) | 0% (0) | 0% (0) | 9.7% (23) | 14.7% (35) | 58.0% (138) |
| Pigs | | | | | | | | | | | | | | | | | | |
| | 2016 | 271 | 98.9% (268) | 99.3% (269) | 5.2% (14) | 81.2% (220) | 62.7% (170) | 30.6% (83) | 4.8% (13) | 3.3% (9) | 36.5% (99) | 1.8% (5) | 0.4% (1) | 0.7% (2) | 0% (0) | 12.9% (35) | 5.9% (16) | 57.9% (157) |
| | 2017 | 477 | 100% (477) | 100% (477) | 7.8% (37) | 81.3% (388) | 48.6% (232) | 17.4% (83) | 8.6% (41) | 1.7% (8) | 42.6% (203) | 0.4% (2) | 0.2% (1) | 0% (0) | 0% (0) | 8.2% (39) | 1.5% (7) | 54.3% (259) |
| | 2018 | 204 | 100% (204) | 100% (204) | 2.0% (4) | 93.6% (191) | 71.6% (146) | 31.9% (65) | 9.8% (20) | 0.5% (1) | 43.1% (88) | 0% (0) | 0% (0) | 0% (0) | 0% (0) | 7.8% (16) | 16.7% (34) | 56.4% (115) |
| Healthy volunteers | | | | | | | | | | | | | | | | | | |
| | 2016 | 239 | 100% (239) | 100% (239) | 6.7% (16) | 64.0% (153) | 58.2% (139) | 16.7% (40) | 3.3% (8) | 1.7% (4) | 23.4% (56) | 0.4% (1) | 0.8% (2) | 0% (0) | 0.8% (2) | 14.2% (34) | 2.5% (6) | 44.4% (106) |
| | 2017 | 107 | 100% (107) | 100% (107) | 13.1% (14) | 87.9% (94) | 82.2% (88) | 28.0% (30) | 16.8% (18) | 7.5% (8) | 62.6% (67) | 0.9% (1) | 0% (0) | 0% (0) | 0% (0) | 14.0% (15) | 2.8% (3) | 70.1% (75) |
| | 2018 | 9 | 88.9% (8) | 88.9% (8) | 0% (0) | 88.9% (8) | 88.9% (8) | 22.2% (2) | 0% (0) | 0% (0) | 33.3% (3) | 0% (0) | 0% (0) | 0% (0) | 0% (0) | 0% (0) | 0% (0) | 33.3% (3) |
| Colonization patients | | | | | | | | | | | | | | | | | | |
| | 2016 | 309 | 100% (309) | 100% (309) | 22.0% (68) | 79.9% (247) | 41.4% (128) | 23.3% (72) | 6.5% (20) | 2.6% (8) | 33.7% (104) | 0.6% (2) | 0.6% (2) | 0% (0) | 0.3% (1) | 10.4% (32) | 2.6% (8) | 49.2% (152) |
| | 2017 | 46 | 100% (46) | 100% (46) | 0% (0) | 82.6% (38) | 82.6% (38) | 41.3% (19) | 4.3% (2) | 6.5% (3) | 37.0% (17) | 0% (0) | 0% (0) | 0% (0) | 0% (0) | 13.0% (6) | 2.2% (1) | 65.2% (30) |
| | 2018 | 9 | 100% (9) | 100% (9) | 22.2% (2) | 100% (9) | 100% (9) | 55.6% (5) | 0% (0) | 0% (0) | 33.3% (3) | 0% (0) | 0% (0) | 0% (0) | 0% (0) | 33.3% (3) | 0% (0) | 88.9% (8) |
| Infection patients | | | | | | | | | | | | | | | | | | |
| | 2016 | 30 | 100% (30) | 100% (30) | 10% (3) | 96.7% (29) | 96.7% (29) | 86.7% (26) | 36.7% (11) | 43.3% (13) | 73.3% (22) | 3.3% (1) | 0% (0) | 0% (0) | 0% (0) | 26.7% (8) | 0% (0) | 93.3% (28) |
| | 2017 | 18 | 100% (18) | 100% (18) | 16.7% (3) | 100% (18) | 88.9% (16) | 55.6% (10) | 22.2% (4) | 22.2% (4) | 66.7% (12) | 0% (0) | 0% (0) | 0% (0) | 0% (0) | 27.8% (5) | 5.6% (1) | 83.3% (15) |
| | 2018 | 11 | 100% (11) | 100% (11) | 27.3% (3) | 90.9% (10) | 90.9% (10) | 81.8% (9) | 27.3% (3) | 18.2% (2) | 54.5% (6) | 0% (0) | 0% (0) | 0% (0) | 0% (0) | 18.2% (2) | 0% (0) | 90.9% (10) |
| Food | | | | | | | | | | | | | | | | | | |
| | 2016 | 51 | 100% (51) | 100% (51) | 5.9% (3) | 78.4% (40) | 54.9% (28) | 19.6% (10) | 2.0% (1) | 3.9% (2) | 31.4% (16) | 0% (0) | 0% (0) | 0% (0) | 0% (0) | 11.8% (6) | 2.0% (1) | 51.0% (26) |
| | 2017 | 63 | 100% (63) | 100% (63) | 1.6% (1) | 81.0% (51) | 34.9% (22) | 6.3% (4) | 4.8% (3) | 0% (0) | 34.9% (22) | 0% (0) | 0% (0) | 0% (0) | 0% (0) | 9.5% (6) | 1.6% (1) | 63.5% (40) |
| | 2018 | 4 | 100% (4) | 100% (4) | 25.0% (1) | 100% (4) | 100% (4) | 75.0% (3) | 0% (0) | 0% (0) | 75.0% (3) | 0% (0) | 0% (0) | 0% (0) | 0% (0) | 50% (2) | 0% (0) | 25.0% (1) |
| Environment | | | | | | | | | | | | | | | | | | |
| | 2016 | 59 | 100% (59) | 100% (59) | 6.8% (4) | 86.4% (51) | 81.4% (48) | 44.1% (26) | 1.7% (1) | 3.4% (2) | 52.5% (31) | 1.7% (1) | 0% (0) | 0% (0) | 0% (0) | 37.3% (22) | 1.7% (1) | 72.9% (43) |
| | 2017 | 40 | 100% (40) | 100% (40) | 30% (12) | 90% (36) | 57.5% (23) | 40% (16) | 2.5% (1) | 17.5% (7) | 35.0% (14) | 0% (0) | 0% (0) | 0% (0) | 0% (0) | 12.5% (5) | 22.5% (9) | 57.5% (23) |
| | 2018 | 1 | 100% (1) | 100% (1) | 0% (0) | 100% (1) | 0% (0) | 100% (1) | 100% (1) | 0% (0) | 100% (1) | 0% (0) | 0% (0) | 0% (0) | 0% (0) | 0% (0) | 100% (1) | 100% (1) |

Data are % (n/N). CL = colistin, PB = polymyxin B, TGC=tigecycline, AMP=ampicillin, AMC=amoxicillin-clavulanate, CTX=cefotaxime, CAZ=ceftazidime, FEP=cefepime, GEN=gentamicin, AMK=amikacin, ETP=ertapenem, IMP=imipenem, MEM=meropenem, FOS=fosfomycin, NIT=nitrofurantoin, CIP=ciprofloxacin.

**Appendix table 3. Median and interquartile ranges for log2 transformed MIC values**

| Source | Year | n | CL | PB | TGC | AMP | AMC | CTX | CAZ | FEP | GEN | AMK | ETP | IMP | MEM | FOS | NIT | CIP |
|---|---|---|---|---|---|---|---|---|---|---|---|---|---|---|---|---|---|---|
| Overall | | | | | | | | | | | | | | | | | | |
| | All | 1948 | 4 (3-4) | 4 (3-4) | -2 (-2--1) | 9 (7-9) | 4 (3-4) | 0 (0-2) | 0 (0-0) | -1 (-1--1) | 1 (0-6) | 2 (1-2) | -3 (-3--3) | -3 (-3--3) | -3 (-3--3) | 3 (3-3.25) | 4 (3-5) | 0 (-2-3) |
| | 2016 | 959 | 4 (3-4) | 4 (3-4) | -2 (-2--1) | 9 (6-9) | 4 (3-4) | 0 (0-2) | 0 (0-0) | -1 (-1--1) | 1 (0-5) | 2 (1-2) | -3 (-3--3) | -3 (-3--3) | -3 (-3--3) | 3 (3-3) | 4 (3-5) | 0 (-2-2) |
| | 2017 | 751 | 4 (4-4) | 4 (3-4) | -2 (-2--1.5) | 9 (8-9) | 4 (3-4) | 0 (0-0) | 0 (0-1) | -1 (-1--1) | 2 (1-6) | 2 (1-3) | -3 (-3--3) | -3 (-3--3) | -3 (-3--3) | 3 (3-3) | 4 (3-5) | 0 (-2-4) |
| | 2018 | 238 | 3 (3-4) | 3 (3-4) | -2 (-3--2) | 9 (8-9) | 4 (4-5) | 0 (0-3) | 0 (0-0) | -1 (-1--1) | 3 (0-6) | 2 (1-3) | -3 (-3--3) | -3 (-3--3) | -3 (-3--3) | 3 (3-4.75) | 4 (4-6) | 0 (-1.75-5) |

Data are median (IQR). IQR=interquartile range. Drug abbreviations are as per Appendix Table 2.

**Appendix table 4. Pairwise Wilcoxon-Mann-Whitney test of log2-transformed MICs for 1948 MCRPEC isolates from all sources**

| Comparison | CL | PB | TGC | AMP | AMC | CTX | CAZ | FEP | GEN | AMK | ETP | IMP | MEM | FOS | NIT | CIP |
|---|---|---|---|---|---|---|---|---|---|---|---|---|---|---|---|---|
| Pig vs. Healthy volunteer | 0.3919 | 0.4241 | <0.0001 | 0.1527 | 0.0004 | 0.0602 | 0.1655 | 0.0804 | 0.4796 | 0.1933 | 0.1933 | 0.0004 | 0.0499 | 0.2026 | <0.0001 | 0.2262 |
| Pig vs. Colonization patient | 0.4044 | <0.0001 | <0.0001 | 0.1281 | <0.0001 | 0.1348 | 0.6841 | 0.0016 | 0.0213 | 0.3871 | 0.3871 | 0.0769 | 0.3838 | 0.3062 | <0.0001 | 0.3049 |
| Pig vs. Infection patient | 0.0009 | 0.4241 | <0.0001 | <0.0001 | <0.0001 | <0.0001 | <0.0001 | <0.0001 | <0.0001 | 0.3871 | 0.3871 | <0.0001 | 0.2337 | <0.0001 | 0.0841 | <0.0001 |
| Pig vs. Food | 0.0113 | 0.0065 | <0.0001 | 0.0256 | <0.0001 | 0.024 | 0.3899 | 0.0107 | 0.3732 | 0.4764 | 0.4764 | 0.1007 | 0.6419 | 0.3816 | 0.0336 | 0.0454 |
| Pig vs. Environment | 0.4044 | 0.0053 | 0.0001 | 0.0097 | <0.0001 | <0.0001 | 0.3446 | <0.0001 | 0.5051 | 0.0041 | 0.0041 | <0.0001 | 0.115 | <0.0001 | 0.0163 | 0.2262 |
| Healthy volunteer vs. Colonization patient | 0.4044 | 0.0002 | 0.0768 | 0.0529 | <0.0001 | 0.024 | 0.2298 | 0.0268 | 0.1127 | 0.0781 | 0.0781 | 0.0002 | 0.0244 | 0.4106 | 0.1652 | 0.131 |
| Healthy volunteer vs. Infection patient | 0.0024 | 0.4241 | 0.0001 | <0.0001 | <0.0001 | <0.0001 | <0.0001 | <0.0001 | <0.0001 | 0.7407 | 0.7407 | <0.0001 | 0.7939 | <0.0001 | 0.0003 | <0.0001 |
| Healthy volunteer vs. Food | 0.0084 | 0.0082 | <0.0001 | 0.0861 | <0.0001 | 0.1321 | 0.6012 | 0.0048 | 0.4349 | 0.1986 | 0.1986 | 0.0028 | 0.115 | 0.4317 | 0.1142 | 0.1668 |
| Healthy volunteer vs. Environment | 0.3919 | 0.0063 | 0.0621 | 0.0048 | 0.0127 | <0.0001 | 0.13 | <0.0001 | 0.4796 | 0.0007 | 0.0007 | 0.0001 | 0.0174 | <0.0001 | <0.0001 | 0.131 |
| Colonization patient vs. Infection patient | 0.0017 | 0.047 | 0.0005 | <0.0001 | <0.0001 | <0.0001 | <0.0001 | <0.0001 | <0.0001 | 0.2979 | 0.2979 | <0.0001 | 0.1452 | <0.0001 | 0.0015 | <0.0001 |
| Colonization patient vs. Food | 0.012 | <0.0001 | <0.0001 | 0.0096 | 0.1239 | 0.0117 | 0.3899 | 0.0002 | 0.4349 | 0.7407 | 0.7407 | 0.1903 | 0.8448 | 0.4816 | 0.257 | 0.03 |
| Colonization patient vs. Environment | 0.4044 | <0.0001 | 0.1282 | 0.0502 | <0.0001 | 0.0005 | 0.354 | <0.0001 | 0.1343 | 0.0224 | 0.0224 | <0.0001 | 0.2613 | <0.0001 | <0.0001 | 0.3746 |
| Infection patient vs. Food | <0.0001 | 0.0385 | <0.0001 | <0.0001 | <0.0001 | <0.0001 | <0.0001 | <0.0001 | <0.0001 | 0.3218 | 0.3218 | <0.0001 | 0.2235 | <0.0001 | 0.0128 | <0.0001 |
| Infection patient vs. Environment | 0.0026 | 0.0277 | 0.0056 | 0.0037 | 0.0003 | <0.0001 | 0.0006 | <0.0001 | 0.0015 | 0.0167 | 0.0167 | 0.0032 | 0.0525 | 0.2259 | 0.3315 | <0.0001 |
| Food vs. Environment | 0.0791 | 0.4241 | <0.0001 | 0.0009 | <0.0001 | <0.0001 | 0.2298 | <0.0001 | 0.3732 | 0.0685 | 0.0685 | <0.0001 | 0.3078 | <0.0001 | 0.0015 | 0.0313 |

The colored frames represent the source which has the higher MIC values for a given pairwise comparison, if statistically significant. Green color represents a pig source; blue represents a healthy volunteer source; pink color represents a colonized inpatient source; red color represents a hospital inpatient with infection source; orange color represents a food source; and purple represents an environmental source.

**Appendix table 5. MLST distributions amongst 688 MCRPEC isolates**

| MLST | ST clonal complex (CC) | Count | Percentage | MLST | ST clonal complex (CC) | Count | Percentage |
|---|---|---|---|---|---|---|---|
| ST10 | ST10 | 100 | 14.53% | ST2345 | ST10 | 3 | 0.44% |
| ST101 | ST101 | 46 | 6.69% | ST34 | ST10 | 3 | 0.44% |
| ST48 | ST10 | 28 | 4.07% | ST162 | ST469 | 3 | 0.44% |
| ST69 | ST69 | 26 | 3.78% | ST1602 | ST10 | 3 | 0.44% |
| ST744 | ST10 | 22 | 3.20% | ST746 | ST10 | 3 | 0.44% |
| ST1244 | ST206 | 18 | 2.62% | ST2705 | / | 3 | 0.44% |
| ST410 | ST23 | 15 | 2.18% | ST3489 | ST10 | 3 | 0.44% |
| ST165 | ST165 | 14 | 2.03% | ST46 | ST46 | 3 | 0.44% |
| ST206 | ST206 | 14 | 2.03% | ST226 | ST226 | 3 | 0.44% |
| ST117 | / | 11 | 1.60% | ST457 | / | 3 | 0.44% |
| ST641 | ST86 | 11 | 1.60% | ST648 | ST648 | 3 | 0.44% |
| ST58 | ST155 | 10 | 1.45% | ST9392 | / | 3 | 0.44% |
| ST9159 | ST10 | 8 | 1.16% | ST345 | / | 3 | 0.44% |
| ST2253 | ST168 | 8 | 1.16% | ST5503 | ST10 | 3 | 0.44% |
| ST189 | ST165 | 8 | 1.16% | ST9412 | ST95 | 3 | 0.44% |
| ST1114 | ST165 | 7 | 1.02% | ST8262 | / | 3 | 0.44% |
| ST542 | / | 7 | 1.02% | ST3274 | ST10 | 2 | 0.29% |
| ST93 | ST168 | 6 | 0.87% | ST5995 | ST165 | 2 | 0.29% |
| ST871 | / | 6 | 0.87% | ST2935 | ST10 | 2 | 0.29% |
| ST453 | ST86 | 6 | 0.87% | ST2466 | / | 2 | 0.29% |
| ST2144 | ST23 | 6 | 0.87% | ST6712 | ST168 | 2 | 0.29% |
| ST25 | / | 6 | 0.87% | ST1258 | / | 2 | 0.29% |
| ST795 | ST278 | 5 | 0.73% | ST13 | ST13 | 2 | 0.29% |
| ST131 | ST131 | 5 | 0.73% | ST167 | ST10 | 2 | 0.29% |
| ST9378 | ST10 | 5 | 0.73% | ST224 | / | 2 | 0.29% |
| ST1684 | / | 5 | 0.73% | ST877 | ST86 | 2 | 0.29% |
| ST1081 | / | 5 | 0.73% | ST3203 | ST10 | 2 | 0.29% |
| ST95 | ST95 | 5 | 0.73% | ST2179 | / | 2 | 0.29% |
| ST156 | ST156 | 5 | 0.73% | ST1193 | ST14 | 2 | 0.29% |
| ST716 | ST10 | 4 | 0.58% | ST354 | ST354 | 2 | 0.29% |
| ST4038 | / | 4 | 0.58% | ST57 | ST350 | 2 | 0.29% |
| ST971 | ST155 | 4 | 0.58% | ST1968 | ST10 | 2 | 0.29% |
| ST4014 | ST101 | 4 | 0.58% | ST1642 | / | 2 | 0.29% |
| ST1196 | / | 4 | 0.58% | ST5871 | ST10 | 2 | 0.29% |
| ST398 | ST398 | 4 | 0.58% | ST359 | ST101 | 2 | 0.29% |
| ST1638 | ST10 | 4 | 0.58% | ST9403 | ST10 | 2 | 0.29% |
| ST617 | ST10 | 4 | 0.58% | ST1249 | / | 2 | 0.29% |
| ST1716 | / | 4 | 0.58% | ST3997 | ST10 | 2 | 0.29% |
| ST7122 | ST10 | 4 | 0.58% | ST88 | ST23 | 2 | 0.29% |
| ST8539 | / | 4 | 0.58% | ST9405 | / | 2 | 0.29% |
| ST3944 | ST10 | 4 | 0.58% | ST1485 | ST648 | 2 | 0.29% |
| ST7056 | / | 4 | 0.58% | ST7652 | ST10 | 2 | 0.29% |
| ST1011 | / | 4 | 0.58% | ST7452 | / | 2 | 0.29% |

| MLST | ST complex | Count | Percentage |
|---|---|---|---|
| ST9373 | / | 1 | 0.15% |
| ST2854 | ST10 | 1 | 0.15% |
| ST7331 | ST10 | 1 | 0.15% |
| ST9381 | / | 1 | 0.15% |
| ST9377 | ST10 | 1 | 0.15% |
| ST2064 | / | 1 | 0.15% |
| ST2526 | / | 1 | 0.15% |
| ST993 | ST10 | 1 | 0.15% |
| ST5215 | ST10 | 1 | 0.15% |
| ST793 | ST206 | 1 | 0.15% |
| ST9380 | / | 1 | 0.15% |
| ST9375 | ST101 | 1 | 0.15% |
| ST2514 | / | 1 | 0.15% |
| ST2913 | ST206 | 1 | 0.15% |
| ST1040 | / | 1 | 0.15% |
| ST1695 | ST10 | 1 | 0.15% |
| ST218 | ST10 | 1 | 0.15% |
| ST2708 | ST10 | 1 | 0.15% |
| ST6272 | / | 1 | 0.15% |
| ST9400 | / | 1 | 0.15% |
| ST5951 | / | 1 | 0.15% |
| ST361 | / | 1 | 0.15% |
| ST9379 | ST10 | 1 | 0.15% |
| ST6815 | ST10 | 1 | 0.15% |
| ST3288 | ST206 | 1 | 0.15% |
| ST6449 | / | 1 | 0.15% |
| ST9382 | ST10 | 1 | 0.15% |
| ST191 | / | 1 | 0.15% |
| ST770 | / | 1 | 0.15% |
| ST4541 | / | 1 | 0.15% |
| ST5542 | ST10 | 1 | 0.15% |
| ST5222 | / | 1 | 0.15% |
| ST2509 | / | 1 | 0.15% |
| ST9383 | / | 1 | 0.15% |
| ST3634 | / | 1 | 0.15% |
| ST155 | ST155 | 1 | 0.15% |
| ST9384 | ST10 | 1 | 0.15% |
| ST1286 | ST10 | 1 | 0.15% |
| ST9387 | / | 1 | 0.15% |
| ST129 | / | 1 | 0.15% |
| ST4741 | ST10 | 1 | 0.15% |
| ST9385 | / | 1 | 0.15% |
| ST9386 | ST206 | 1 | 0.15% |
| ST73 | ST73 | 1 | 0.15% |
| ST3941 | ST168 | 1 | 0.15% |
| ST9388 | / | 1 | 0.15% |
| ST6706 | ST10 | 1 | 0.15% |
| ST9391 | / | 1 | 0.15% |

| MLST | ST complex | Count | Percentage |
|---|---|---|---|
| ST9394 | / | 1 | 0.15% |
| ST9395 | ST206 | 1 | 0.15% |
| ST799 | / | 1 | 0.15% |
| ST6335 | ST165 | 1 | 0.15% |
| ST1308 | ST86 | 1 | 0.15% |
| ST9398 | ST10 | 1 | 0.15% |
| ST767 | ST155 | 1 | 0.15% |
| ST9396 | ST10 | 1 | 0.15% |
| ST9402 | / | 1 | 0.15% |
| ST9397 | / | 1 | 0.15% |
| ST7054 | / | 1 | 0.15% |
| ST9404 | / | 1 | 0.15% |
| ST9399 | ST165 | 1 | 0.15% |
| ST2521 | ST446 | 1 | 0.15% |
| ST29 | ST29 | 1 | 0.15% |
| ST4214 | / | 1 | 0.15% |
| ST4358 | / | 1 | 0.15% |
| ST1136 | / | 1 | 0.15% |
| ST9401 | ST155 | 1 | 0.15% |
| ST707 | / | 1 | 0.15% |
| ST9415 | ST10 | 1 | 0.15% |
| ST9406 | ST10 | 1 | 0.15% |
| ST6908 | / | 1 | 0.15% |
| ST9407 | ST10 | 1 | 0.15% |
| ST3856 | / | 1 | 0.15% |
| ST3014 | / | 1 | 0.15% |
| ST192 | / | 1 | 0.15% |
| ST602 | ST446 | 1 | 0.15% |
| ST657 | / | 1 | 0.15% |
| ST847 | / | 1 | 0.15% |
| ST207 | ST10 | 1 | 0.15% |
| ST484 | ST168 | 1 | 0.15% |
| ST5487 | ST155 | 1 | 0.15% |
| ST7073 | / | 1 | 0.15% |
| ST9410 | ST69 | 1 | 0.15% |
| ST2309 | ST38 | 1 | 0.15% |
| ST9411 | ST206 | 1 | 0.15% |
| ST9408 | ST405 | 1 | 0.15% |
| ST1589 | / | 1 | 0.15% |
| ST154 | / | 1 | 0.15% |
| ST9409 | ST10 | 1 | 0.15% |
| ST8540 | / | 1 | 0.15% |
| ST6438 | / | 1 | 0.15% |
| ST3937 | ST10 | 1 | 0.15% |
| ST4623 | / | 1 | 0.15% |
| ST710 | / | 1 | 0.15% |
| ST656 | ST10 | 1 | 0.15% |
| ST642 | ST278 | 1 | 0.15% |

| | | | |
|---|---|---|---|
| ST2216 | / | 1 | 0.15% |
| ST6756 | / | 1 | 0.15% |
| ST5086 | ST206 | 1 | 0.15% |
| ST1415 | ST10 | 1 | 0.15% |
| ST2936 | ST10 | 1 | 0.15% |
| ST9390 | / | 1 | 0.15% |
| ST9389 | ST20 | 1 | 0.15% |
| ST9393 | / | 1 | 0.15% |
| ST9413 | ST156 | 1 | 0.15% |
| ST3933 | / | 1 | 0.15% |
| ST773 | ST168 | 1 | 0.15% |
| ST2223 | ST10 | 1 | 0.15% |
| ST3076 | / | 1 | 0.15% |
| ST9414 | / | 1 | 0.15% |
| ST711 | / | 1 | 0.15% |
| ST8154 | / | 1 | 0.15% |

**Appendix table 6. Reference plasmids for BLAST analysis**

| Plasmids name | Source | Size(bp) | Inc-type | Accession number | References |
|---|---|---|---|---|---|
| pHNSHP45 | *E. coli* | 64015 | IncI2 | KP347127.1 | Liu YY, *et al*. Lancet Infect Dis. 2016. |
| pHNSHP45-2 | *E. coli* | 251493 | IncHI2 | KU341381.1 | Zhi C, *et al*. Lancet Infect Dis. 2016. |
| pMCR1_IncX4 | *K. pneumoniae* | 33287 | IncX4 | KU761327.1 | Li A, *et al*. Antimicrob Agents Chemother. 2016. |
| pKH457-3-BE | *E. coli* | 79798 | IncP | KU353730.1 | Malhotra-Kumar S , et al. Lancet Infect Dis. 2016. |
| pCQ02-121 | *E. coli* | 48350 | IncN | KU647721.2 | / |
| pMR0516mcr | *E. coli* | 225707 | IncF | KX276657.1 | McGann P, *et al*. Antimicrob Agents Chemother. 2016. |
| pKP81-BE | *E. coli* | 91041 | IncFII | KU994859.1 | Xavier BB, *et al*. J Antimicrob Chemother. 2016. |
| pH226B | *E. coli* | 209401 | IncHI1 | KX129784.1 | Zurfluh K, *et al*. Antimicrob Agents Chemother. 2016. |
| pEC2-4 | *E. coli* | 235403 | IncHI1A | CP016184.1 | / |
| p1108-IncFIB | *E. coli* | 193873 | IncFIB | MG825378.1 | / |
| pHKSHmcr1_P2_p1 | *E. coli* | 47818 | IncP2 | MF136778.1 | Chan WS, *et al.* BMC Infect Dis. 2018. |
| pZR78 | *E. coli* | 91281 | IncpO111 | MF455226.1 | Shen Y, *et al.* Environ Int. 2019. |
| pEC2_1-4 | *E. coli* | 230278 | IncHI1B | CP016183.1 | / |
| pHYEC7-mcr1 | *E. coli* | 97559 | IncY | KX518745.1 | Li R, *et al*. J Antimicrob Chemother. 2017. |

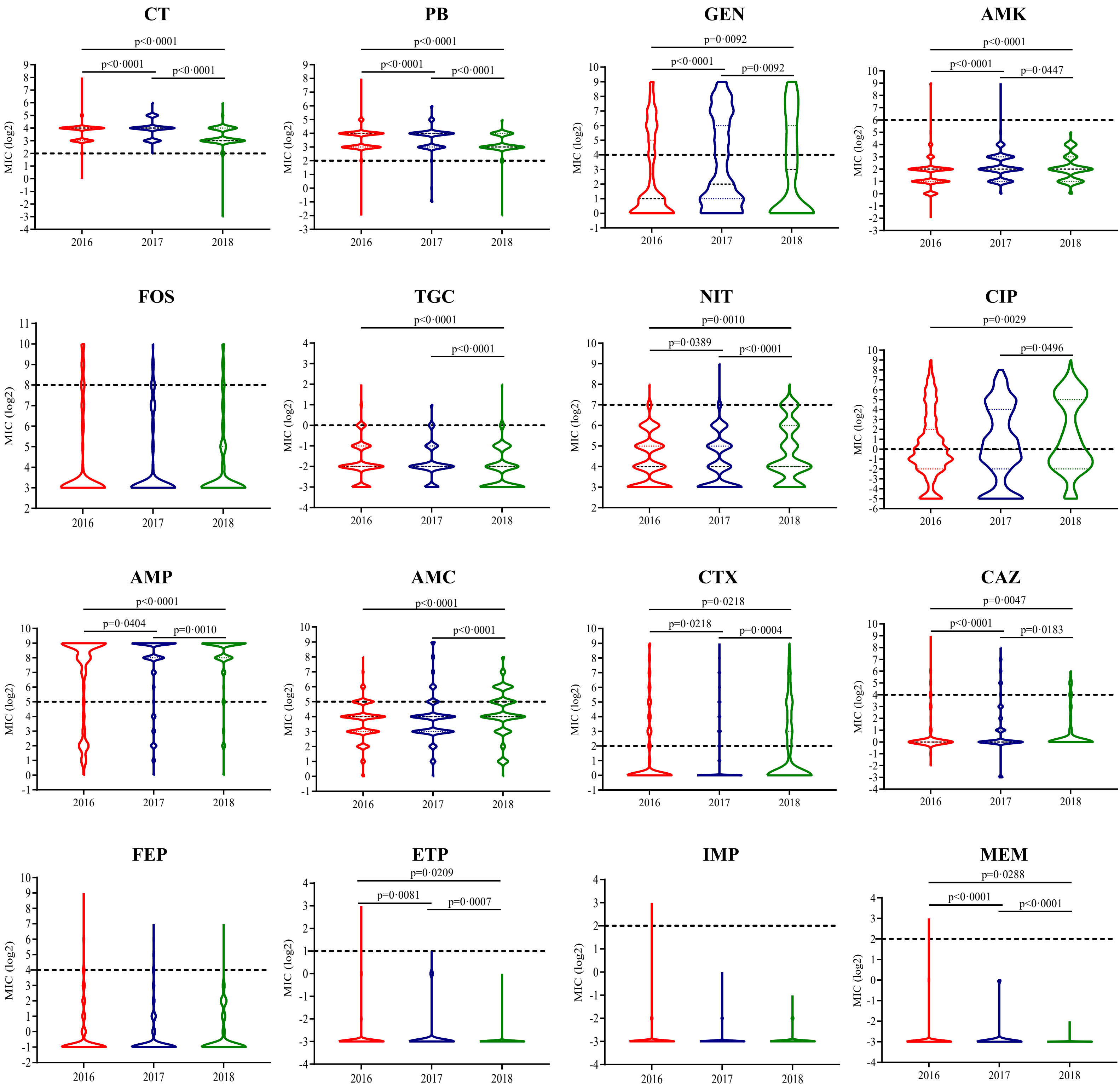


Appendix figure 1. Violin plots of MIC values of 1948 mcr-1 -positive E. coli isolates for each antimicrobial by year of sampling.

The red color represents data for 2016, blue for 2017, and green for 2018. Bonferroni-adjusted p-values less than 0·05 are highlighted. CL = colistin, PB = polymixin B, TGC=tigecycline, AMP=ampicillin, AMC=amoxicillin-clavulanate, CTX=cefotaxime, CAZ= ceftazidime, FEP=cefepime, GEN=gentamicin, AMK=amikacin, ETP=ertapenem, IMP=imipenem, MEM=meropenem, FOS= fosfomycin, NIT=nitrofurantoin, CIP=ciprofloxacin.

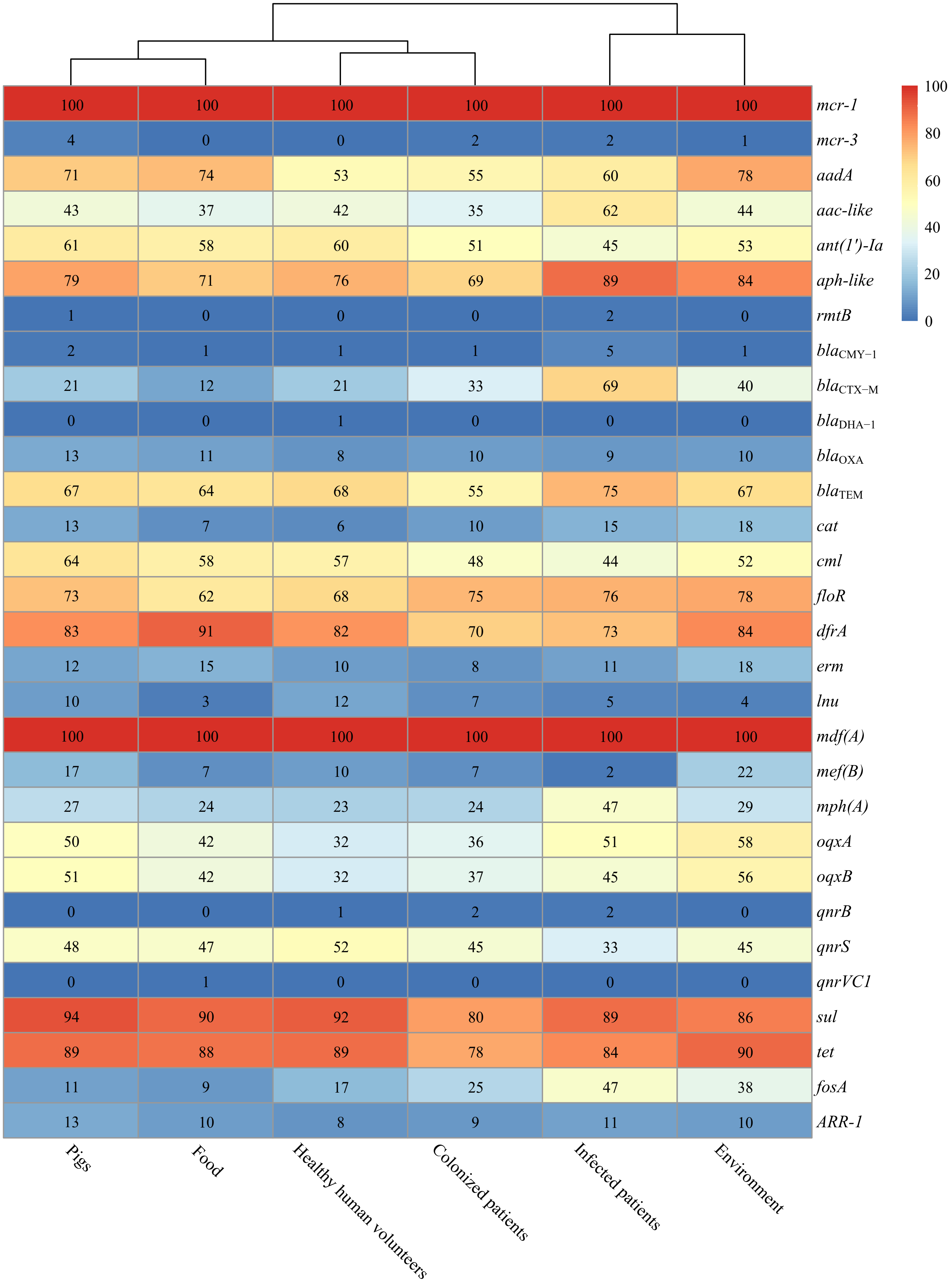


Appendix figure 2. Detection of antimicrobial resistance (AMR) genes in 688 mcr-1-positive E. coli (MCRPEC) by isolate source.

Each cell in the table indicates the percentage of MCRPEC isolates by source that contain the respective AMR gene e.g.. top left cell indicates that 100% of pig isolates carried the mcr-1 gene.

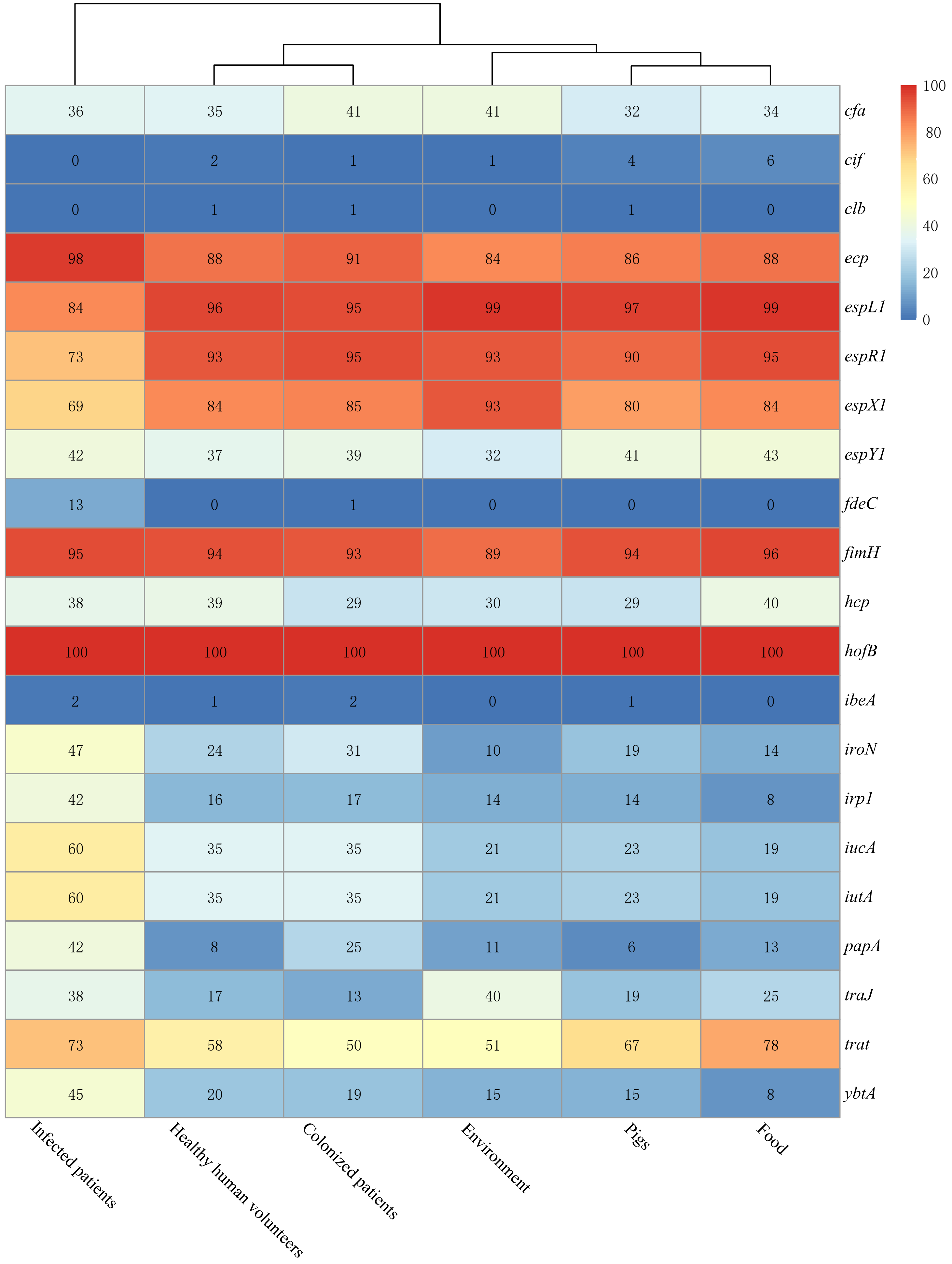


Appendix figure 3. Detection of virulence genes in 688 mcr-1-positive E. coli (MCRPEC) by isolate source.

Each cell in the table indicates the percentage of MCRPEC isolates by source that contain the respective virulence gene e.g. top left cell indicates that 36% of isolates from patients with MCRPEC infections contained cfa.

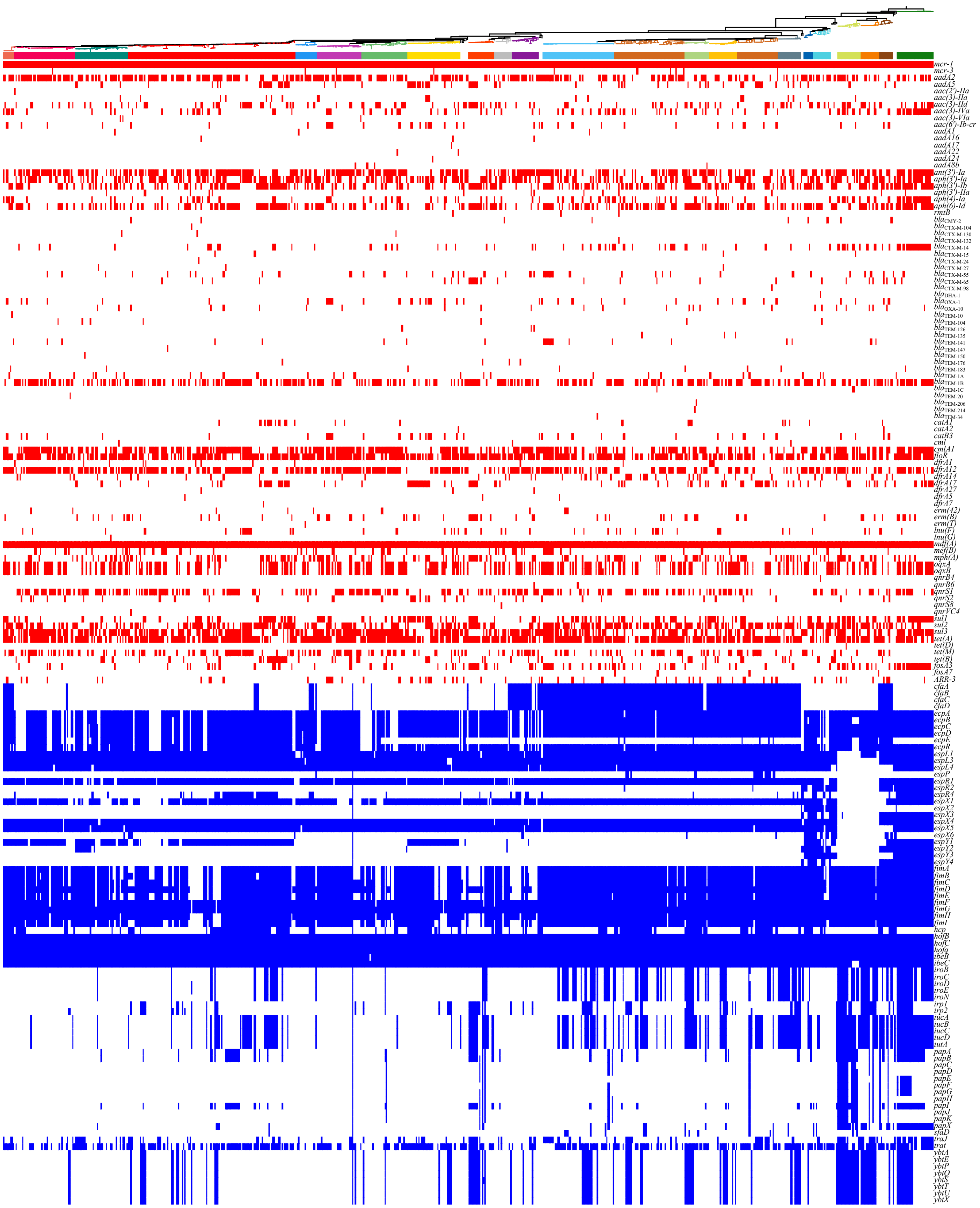


Appendix figure 4. Phylogenetic distribution of antimicrobial resistance (AMR) and virulence genes for 688 mcr-1-positive E. coli (MCRPEC).

The ML phylogeny derived from cgSNPs is at the top of the plot; lineage colors denote sequence clusters (SCs; as in Figure 4). In the heatmap, red blocks denote the presence of an AMR gene, and blue blocks the presence of a virulence gene. Individual genes identified in at least one isolate are listed on the left of the panel.

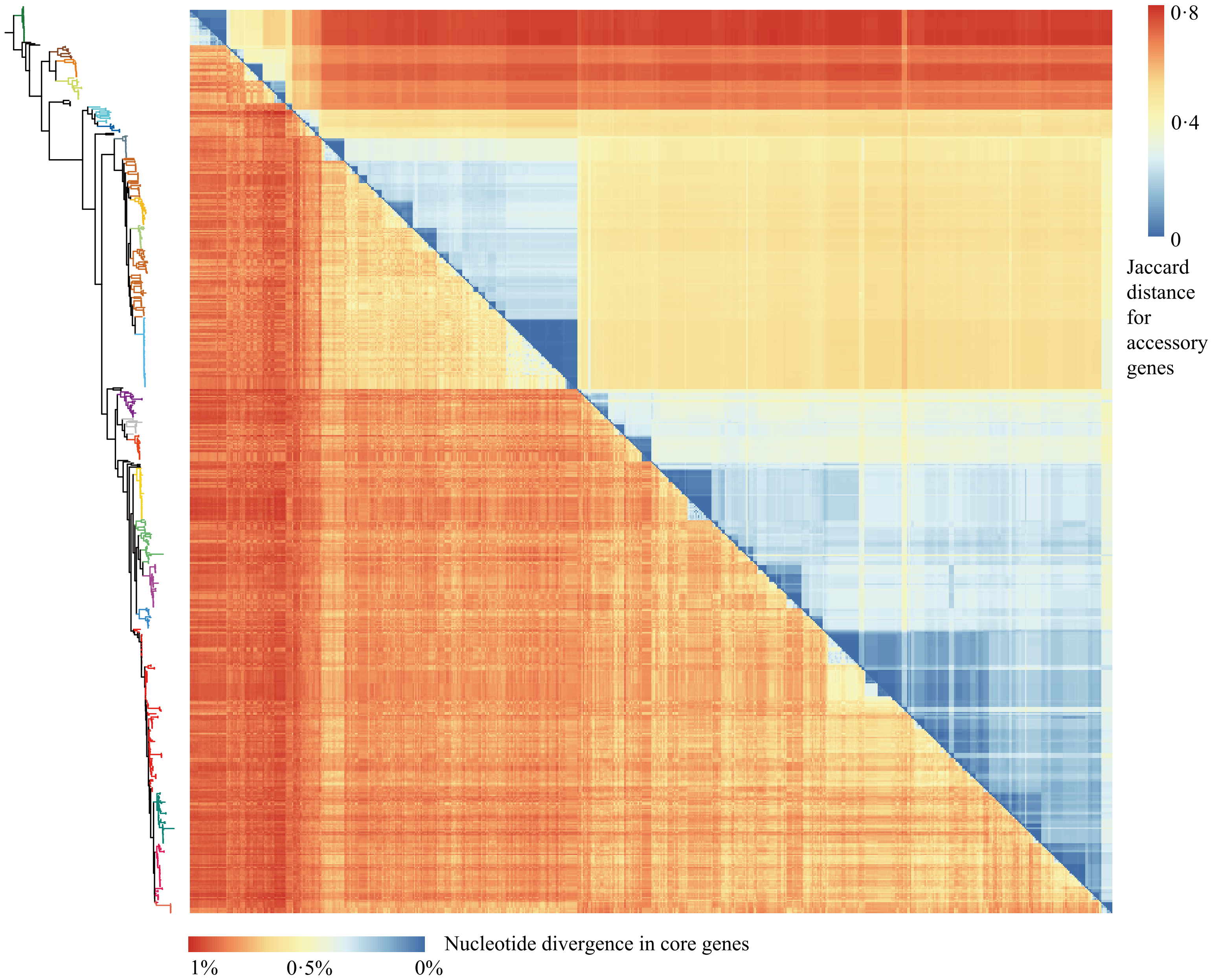


Appendix figure 5. Matrices of pairwise distances of core genes and accessory genes for 688 mcr-1-positive E. coli (MCRPEC).

Pairwise accessory gene differences (Jaccard distance, upper right triangle) and nucleotide divergence in core genes (lower left triangle), ordered against the maximum likelihood phylogeny generated from cgSNPs (left-hand side of panel). The two matrices were highly correlated (p<0·0001, r=0·8107, Mantel test with 1000 permutations).

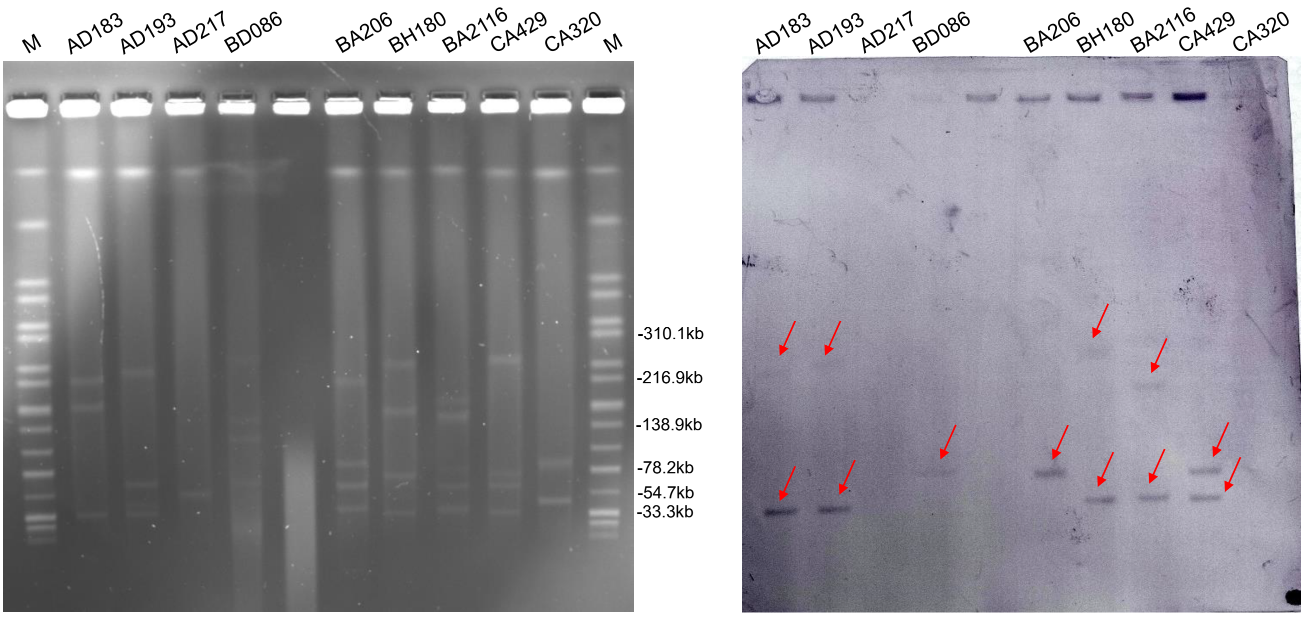


Appendix figure 6. (A) S1-PFGE and (B) Southern blot for nine isolates which harbored more than one common mcr-1-associated plasmid Inc type.

The red arrows indicate the plasmid location of mcr-1, confirming the dual location of mcr-1 in two plasmids in five isolates.

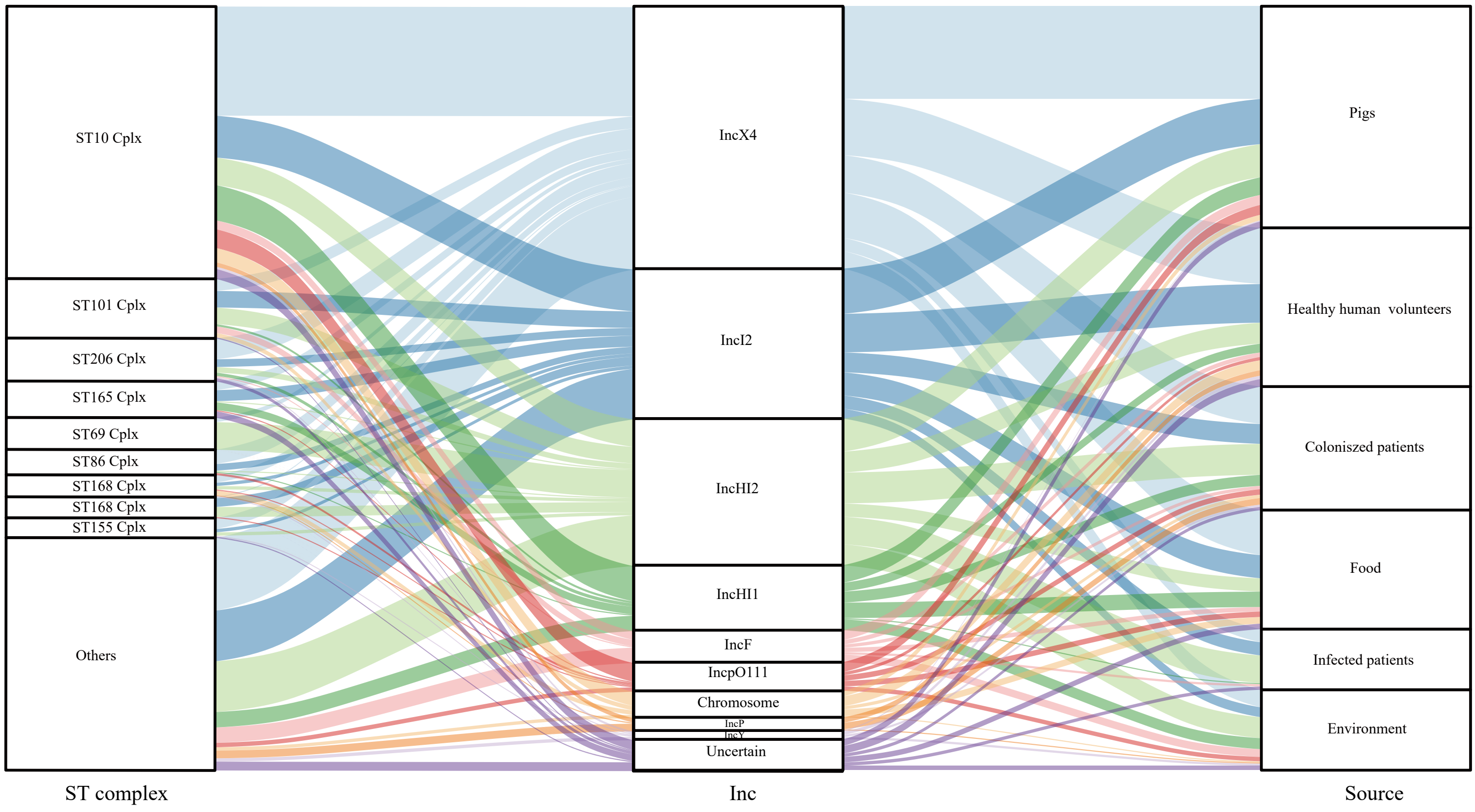


Appendix figure 7. Alluvial diagram of *mcr-1*-positive *E. coli* (MCRPEC) ST clonal complex, *mcr-1* context, and isolate source.

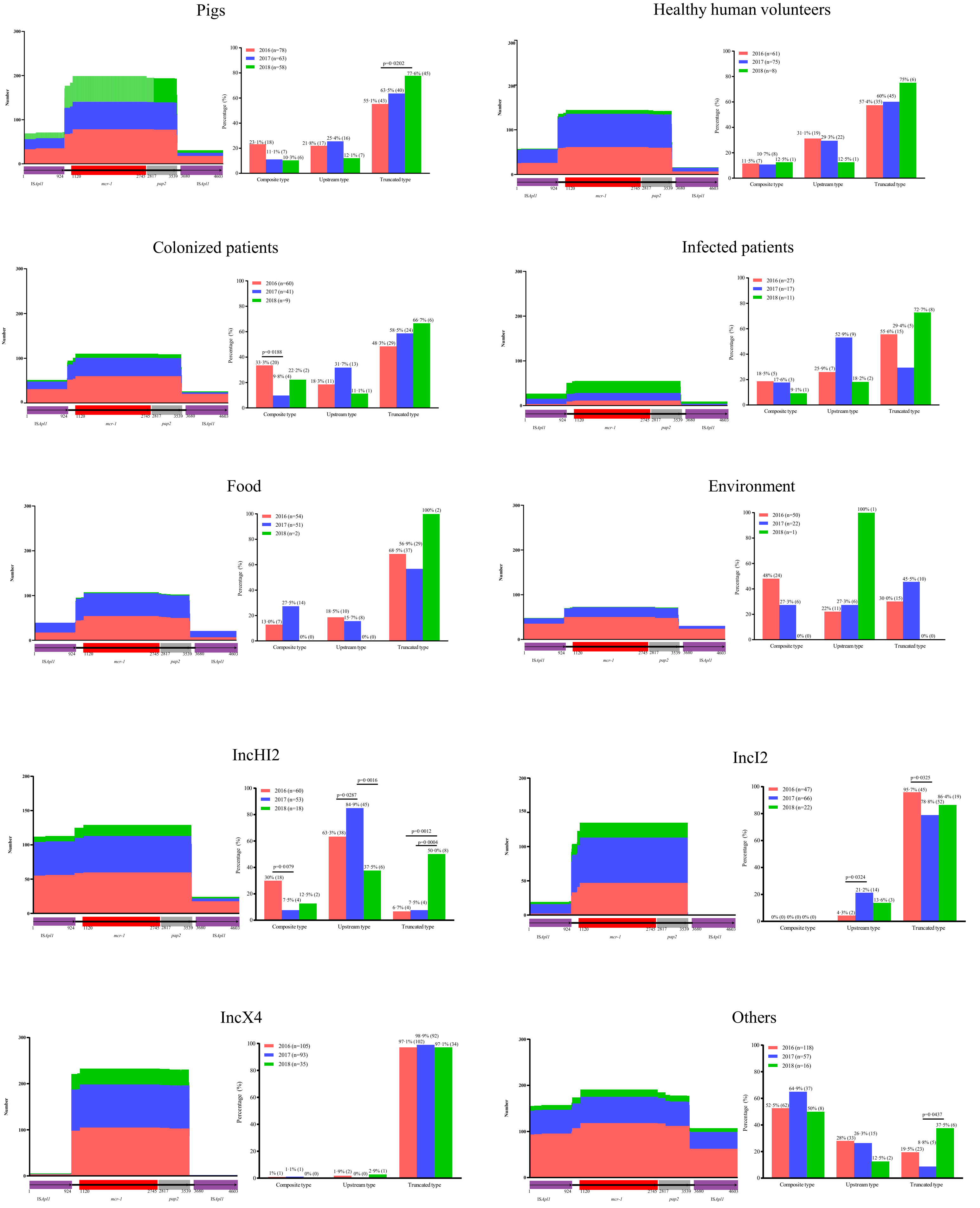


Appendix figure 8. The structural dynamics of the Tn*6330* genetic element carrying *mcr-1*.

Length distribution of the sequence alignments across sampling years by isolate source, and by *mcr-1* plasmid type. Histogram of prevalence for each genetic element across three years.

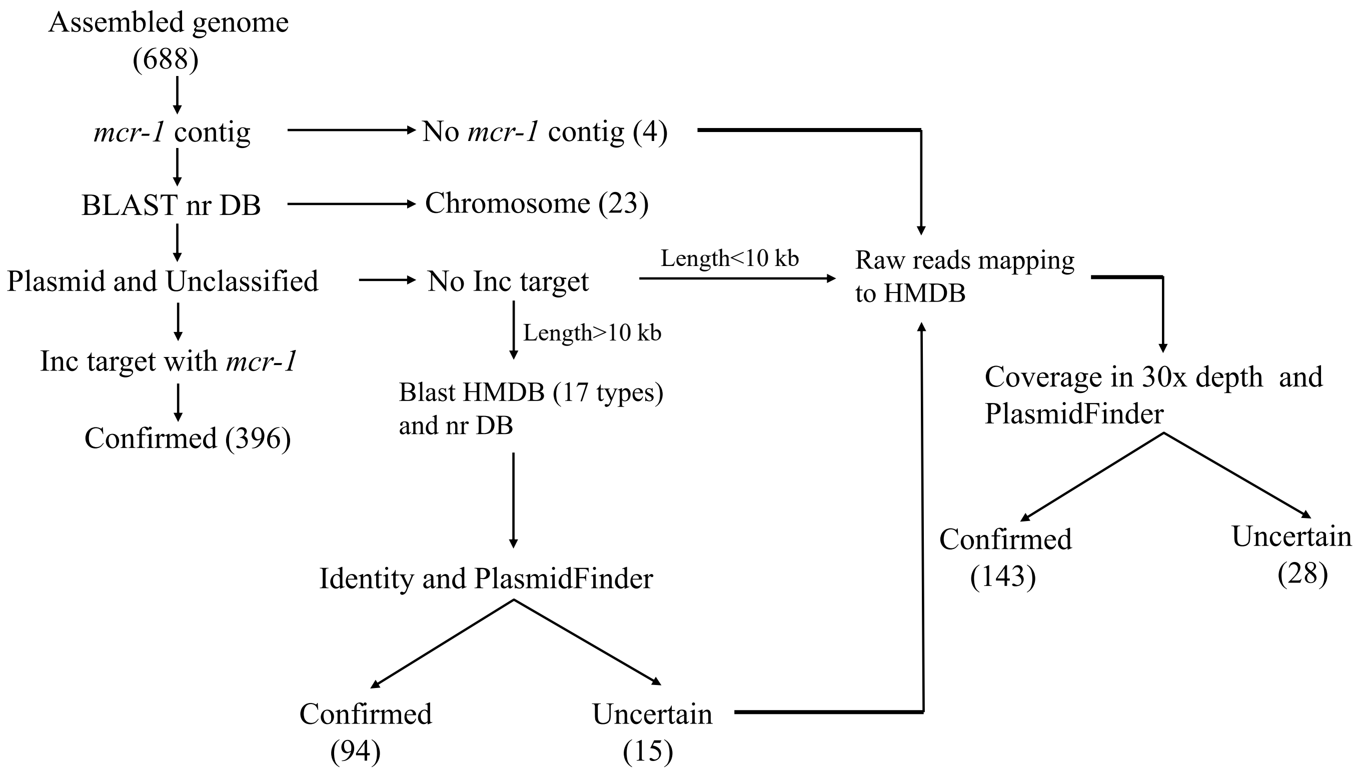


Appendix figure 9. Flowchart for identification of *mcr-1* location.

HMDB=homemade database which included reference plasmids in appendix table 6. Nr DB=nonredundant database.

### Appendix methods

#### Plasmid conjugation

Plasmid conjugation experiments were performed using streptomycin-resistant *E. coli* C600 as the recipient. Briefly, donor and recipient isolates were cultured overnight, set up as cell suspensions, and then 1:100 dilutions of the cell suspensions were sub-cultured for 2.5 hours at 37ºC, respectively. Donor and recipient populations were then mixed at a ratio of 1:9. Transconjugants were selected on LB agar plates supplemented with streptomycin (2000 mg/L) and colistin (2 mg/L). The presence of *mcr-1* in transconjugants was confirmed using PCR and Sanger sequencing.

#### S1-PFGE and Southern blot

The exact plasmid location of the *mcr-1* gene for nine isolates, which harbored more than one common *mcr-1*-associated plasmid replicon from WGS-based plasmid replicon typing, was determined by S1-nuclease digestion and pulsed-field gel electrophoresis (S1-PFGE) with Southern blot hybridizations. Whole-cell DNA of all nine *mcr-1*-producing isolates was extracted and embedded in gold agarose gel plugs (SeaKem® Gold Agarose, Lonza, USA). The plugs were digested with S1 nuclease (TaKaRa, Dalian, China), and the DNA fragments were separated by PFGE. Southern blot hybridizations of plasmid DNA were performed with a DIG-labeled *mcr-1* probe according to the manufacturer's instructions (Roche Diagnostics, Germany).

#### Whole-genome sequencing and bioinformatic analyses

##### *DNA extraction and whole-genome sequencing (WGS)*

A total of 755 presumed MCRPEC isolates (identified using MALDI-TOF MS +/- 16s rDNA sequencing), were subjected to whole genome sequencing. The Qiagen Blood and Tissue kit (Qiagen, Hilden, Germany) was used to extract genomic DNA for *mcr-1*-positive isolates according to the manufacturers' instructions. DNA libraries were constructed with 350bp paired-end fragments and sequenced using an Illumina HiSeq 2000 platform (Illumina, Inc, USA). The sequencing yielded >200-fold coverage sequences (1G) of raw paired-end (150bp) per isolate. Raw sequencing data have been deposited at the NCBI-Sequence Read Archive (Project accession number PRJNA593695).

***Raw read processing and de novo genome assembly***

Adaptor, unreliable reads and low-quality bases of raw reads were trimmed using trimmomatic v0·39 with default parameters.[1] SPAdes v3·13·1 with "--careful" parameter and automatically determined k-mer values (21,33,55,77) were used to perform draft genome *de novo* assembly.[2] Contigs less than 500bp in size were excluded for further analysis. WGS based species identification was performed using JSpeciesWS v3.2.7.[3] Isolates not confirmed as *E. coli* using WGS-based species identification, and contaminated/mixed sequences, were excluded from subsequent analyses.

***Prediction and annotation of open reading frames (ORFs)***

ORFs was predicted using prodigal v2·6·3 and annotated by Prokka v1·13·3.[4,5] Antimicrobial resistance genes, plasmid replicons and virulence factors were identified using SRST2 v0·2 by mapping reads to ResFinder, PlasmidFinder and VFDB database,[6-8] meanwhile, the locations of these traits in draft genome level were annotated using ABRicate 0·8·7 with "--minid 75 --mincov 60" parameters (https://github.com/tseemann/abricate). *In silico* multilocus sequence typing (MLST) was assigned using Enterobase (http://enterobase.warwick.ac.uk/). New alleles and STs were uploaded on Enterobase database. Insertion sequence elements were detected using ISFinder (https://www-is.biotoul.fr/). Clusters of orthologous genes (COG) were annotated using EggNOG v5·0.[9]

***Pan-genome analysis and phylogenetic construction***

Pan-genome analyses of sequences was performed using Roary,[10] and a concatenated alignment of genes shared among ≥99% of all isolates (core genome) was extracted using Mafft v7·407.[11] Core genome single nucleotide polymorphisms (cgSNPs) were extracted from this concatenated alignment using SNP-sites.[12] A maximum likelihood (ML) phylogeny for cgSNPs was reconstructed using RAxML v8·2·10 implementing a generalized time-reversible nucleotide substitution model with a Γ distribution (GTR+G).[13] Branch lengths and bootstrap supports for bipartitions were estimated using 1000 bootstrap replicates. Population structure was assessed using cgSNPs with hierBAPS, which was run four times using maximum clustering sizes of 20, 40, 60 and 80.[14] Sequence clusters (SCs) identified using hierBAPS were labeled on the core genome phylogeny on iTOL (https://itol.embl.de/). Accessory genes with a prevalence of between 5%-95%, were included for network analysis and PGWAS. Jaccard distance was used to assess relation between each isolate; a value of <0·5 was chosen to represent isolates were related. Genetic relationships between isolates were represented as a network using Cytoscape v3·7·2.[15]

***Identification of mcr-1 location and associated plasmid or chromosomal contexts***

The workflow of our approach to assigning *mcr-1* location in assembled MCRPEC is summarized in appendix figure 9. Contigs containing *mcr-1* were extracted from the isolate assemblies, and BLASTn with default parameters comparisons made to NCBI's non-redundant nucleotide database (nr DB). If the top match was chromosomal, the context was designated "chromosomal". If the top match was to a plasmid reference, the context was designated as "plasmid". For contigs containing a plasmid replicon following *in silico* plasmid typing (see above), *mcr-1* was deemed present in that plasmid type. If no replicon type was identified on an *mcr-1*-containing plasmid contig, we devised the following strategy. First, a homemade database (HMDB) of reference *mcr-1* plasmid sequences was created (appendix table 6), and used as a reference in conjunction with the nr DB. For contigs >10kb, BLASTn[16] was performed against both the HMDB and the nr DB, and the top match to either used for plasmid type assignation. For contigs <10kb, for isolates in which no *mcr-1*-containing contig was identified in the first instance, and for isolates that could not be assigned by the steps above, raw reads were mapped to the HMDB (appendix table 6) using bwa.[17] 30-folds depth identity and coverage were counted using samtools and bamdst (https://github.com/shiquan/bamdst), and assignations made on the basis of the highest 30-folds depth coverage result. For twenty-eight isolates the location of *mcr-1* remained uncertain (appendix figure 9).

All contigs large than 1000bp were aligned with identified plasmid reference sequence using BLASTn with default parameters.[16] The contigs, which aligned length large than 60% coverage, were defined as *mcr-1*-plasmid frame sequences. The extraction was assessed using QUAST.[18] ORF prediction and annotation for plasmid frame sequences were done as described before.

**Genetic context (Tn*6330*) of *mcr-1* extraction**

Tn*6330* (IS*Apl1*-*mcr-1*-*pap2*-IS*Apl1*, 4603bp, accession number KX084394·1) was screened for in 684 *mcr-1*-harboring contigs using BLASTn with default parameters,[16] and the genetic context of *mcr-1* was confirmed in 449 samples. For the remaining 235 *mcr-1*-contigs, in which *mcr-1* was located towards the end of the contig (distance <1·2kb upstream of *mcr-1*, or <2kb downstream of *mcr-1*) precluding the *in silico* identification of Tn*6330*/ IS*Apl1*, we performed PCR to verify presence of IS*Apl1* and obtained sequences using Sanger sequencing. The primers for “upstream” Tn*6330* screening were designed from the start of the upstream IS*Apl1* sequence up to and including 305bp of *mcr-1* (F:5’-ATGATTTTACTCGCACAGG-3’; R:5’-TCATAGACCGTGCCATAAG-3’, segment length 1424bp, annealing temperature 55℃, extension time 1min30s). The primers for “downstream” Tn*6330* screening were designed from the 304bp of *pap2* downstream of *mcr-1* to the end of the downstream IS*Apl1* sequence (F:5’-GTGCGATCATCATCGTG-3’; R:5’-TCAAACCAAGTGCAACG-3’, segment length 1228bp, annealing temperature 55℃, extension time

1min30s). Composite Tn*6330* sequences were identified in 4 additional isolates, using *mcr-1* and *pap2* full length sequencing as described previously. Sanger sequencing results were manually combined with *mcr-1*-harboring contigs.

**Core-genome-wide association analysis (CGWAS) and pan-genome-wide association analysis (PGWAS)**

Core genes and accessory genes were separated for CGWAS and PGWAS, respectively. Association analysis was performed by Scoary v1·6·16,[19] which uses a gene presence/absence dataset. 43648 cgSNPs derived from the concatenated alignment of core genes using Roary (see above) were converted to Scoary format for subsequent analysis. A total of 5700 accessory genes were included in PGWAS.

(1) For cgSNPs associated with pre- (2016) and post- intervention (combined 2017 and 2018) periods, Scoary was run with the ML tree produced above.

(2) For PGWAS pre- (2016) and post-intervention (combined 2017 and 2018), Scoary was run with the ML phylogeny produced by cgSNPs previously, collapsing genes that were identically distributed and using 10,000 empirical label-switching permutations. A gene cluster was considered significant by Scoary only if: (a) the Benjamini-Hochberg-corrected p-value was less than 0·05; (b) the worst p-value from the pairwise comparison algorithm was below 0·05, and (c) the empirical p-value based on permutations was lower than 0·05.

(3) For PGWAS of different epidemiological groups, our aim was to identify gene clusters that might be overrepresented in specific groups, thus, Scoary was run with "--nopair_wise" flag and a Benjamini-Hochberg-corrected p-value less than 0·0001 was used as the significance threshold.

12. Page AJ, Taylor B, Delaney AJ, et al. SNP-sites: rapid efficient extraction of SNPs from multi-FASTA alignments.

*Microb Genom* 2016; **2**(4): e000056.
